# The modification of DNA with indole-linked nucleotides alters its sensitivity to enzymatic cleavage

**DOI:** 10.1101/2025.01.26.634936

**Authors:** Suresh Lingala, Anastasiia Fisiuk, Michelle Stephen, Raja Mohanrao, Judah Klingsberg, Simon Vecchioni, Ealonah S Volvovitz, Sergei Rozhkov, Prabodhika Mallikaratchy

## Abstract

We describe the synthesis of C-5 indole-tagged pyrimidine and C-8 indole-tagged purine nucleoside phosphoramidites and their incorporation into double-stranded DNA 15 base pairs in length. Of the 23 sequence modifications tested, two induced the DNA duplex to adopt a Z-like left-handed conformation under physiological salt conditions, bypassing the specific sequences typically required for a left-handed Z-DNA structure. The impact of these modifications varied with the linker type: flexible propyl linkers exhibited distinct effects compared to rigid propargyl linkers. Notably, modifications positioned directly on or near a restriction site emphasized the pivotal role of linker rigidity in controlling DNA conformation. Specifically, the conformational change induced by the flexible linker impacted nuclease and restriction endonuclease cleavage, reducing sequence specificity. In contrast, the rigid linker suppressed this effect. Furthermore, our findings indicate that nucleic acid duplexes modified with indole-linked nucleotides using a flexible propyl linker have a pronounced tendency to form BZ or Z-like regions in longer DNA sequences. A higher density of modifications may even induce a full Z-like conformation throughout the duplex. These modified nucleotides hold potential for the development of novel antisense therapeutics and introducing valuable tools for in vitro screening of small molecules targeting distorted B-DNA, BZ-DNA, and Z-DNA structures.

## Introduction

Growing evidence suggests that noncanonical conformations (NCS) of nucleic acids play a major role in regulating cellular activities.^1, 2^ The important, yet not fully understood, role of NCS is further validated by data obtained from sequencing the human genome, which shows that more than 50% of it is composed of repeat DNA, out of which 13% of total DNA is comprised of simple sequence repeats that can form NCS.^3^ Owing to the poor understanding of NCS in regulating cellular function, the occurrence of NCS was initially considered to have no biological significance. However, recent discoveries related to NCS suggest otherwise. Repetitive sequences with a high propensity to form NCS have been identified in promoter regions, and replication origins suggesting their pivotal role in biological processes.^1^ Interestingly, while the occurrence of NCS is transient, driven by unfavorable folding, NCS still seems to play a significant role in the regulation of DNA replication and gene expression, altering DNA-protein interactions, regulating chromosomal organization, transcription and translation, and processing pre-mRNA.^4–9^ The most common NCS include G-quadruplex formation, i-motifs, R-loops, triplex and cruciform structures, supercoiling, bubbles, H-DNA, A-DNA, Z-DNA, and B-Z junction structures.^3, 10^ ^11, 12^

The left-handed form of DNA, termed Z-DNA, was initially discovered by Rich and coworkers.^13^ It is found in viruses, bacteria, yeast, flies, and mammals, including humans.^14–17^ Z-DNA formation is predominantly driven by alternating syn- and anti-base conformation of GC dinucleotide repeat sequences.^13^ The existence of Z-DNA conformation in the promoter sites is implicated in cancer, autoimmune diseases and neurological diseases, such as Alzheimer’s disease.^8, 18, 19^ The B-DNA to Z-DNA transition seems to suggest that the induction of structural instability mediated by Z-conformation permeating large-scale deletions in genes.^20^ Chemical base modifications, such as 5-methylcytosine adducts, which are products of epigenetic modification, shift the equilibrium from B-DNA conformer to Z-DNA conformer, as demonstrated in chromatin compaction and transcription regulation.^21–23^ Furthermore, Z-DNA binding proteins, the class of proteins that recognize the left-handed Z-form of DNA and RNA, include important enzymes. For example, Adenosine Deaminase Acting on RNA 1 (ADAR1), is involved in RNA editing by converting adenosine to inosine, and it plays a significant role in cancer and autoimmune disorders.^24–27^ Intriguingly, ADAR has a characteristic affinity towards Z-DNA.^28^ Despite repeated observations of the presence of Z-DNA structures and their implications in altering cellular processes, the investigation of Z-DNA in *vitro* and *in vivo* systems has been challenging. This can be attributed to the instability of Z-DNA structures and the transient appearance of Z-DNA in cellular events, thus limiting the fundamental understanding of Z-DNA structure and function.

To generate Z-DNA conformers, Gannett and others have incorporated C-8 aryl guanine in short oligos and successfully demonstrated the stable formation of sequence dependent Z-DNA at physiological conditions.^29^ Another way of inducing stability of Z-conformation is by introducing high concentrations of cations or spermine, spermidine, hexamine cobalt, and ruthenium complexes to neutralize the negative charge of the zigzag phosphate backbone.^30, 31^ Sugiyama and coworkers demonstrated that *in vitro* synthetic sequence-specific methylated guanosine adducts still require high salt concentrations to maintain a stable sequence dependent Z-DNA conformer.^32^ Recently, Damha and his colleagues showed C2′-fluorinated nucleic acids can adapt Z-conformations of nucleobases in a sequence dependent manner at physiological conditions.^33^ Herein, We aimed to investigate whether the formation of Z-DNA or BZ-DNA could be induced independently of the sequence context by introducing complementary strands modified with C-5-propyl/propargyl indole-substituted pyrimidines (U, C) and C-8-propyl/propargyl indole-substituted purines (A, G) into a 15-nucleotide-long DNA duplex. Also, we explored if the linker rigidity plays a role in inducing a conformational switch. To the best of our knowledge, no reports have explored the impact of linker rigidity in modified nucleic acids on their promiscuity toward restriction enzymes, as facilitated by conformational adaptation in a sequence independent manner.

Inspiration to use the indole was drawn from the privileged molecular interactions mediated by indole moiety in amino acids and large number of currently available pharmaceuticals.^34^ Indole’s unique chemical characteristics arise from its ability to form amphipathic and dipole interactions via imino hydrogen and hydrophobic interactions via the aryl group owing to its bicyclic nature. Additionally, indole has been readily used in understanding NCS, such as triplex structures that arise by the ability of their base pairs to act as both hydrogen bond donors and acceptors with nitro or formyl indole-modified nucleosides.^35, 36^ Therefore, we envisioned that modifying the carbon atom at the 5^th^ position on a pyrimidine ring, and carbon atom at the 8^th^ position on a purine ring with indole through a linker, would generate Z-inducing DNA chimeras (ZImera) with higher propensity at physiological conditions independent of sequence composition and other parameters, such as application of positively charged neutralizers, providing a platform to explore the role of Z conformers in cellular fate.

## Materials & Methods

All chemicals and solvents were purchased from Sigma-Aldrich, Fisher Scientific, Oakwood chemicals and Ambeed chemicals and used without further purification. Reactions were monitored by thin layer chromatography (TLC) using Merck pre-coated silica plates (Silica Gel 60 F254, 0.25 mm). TLC plates were visualized under ultraviolet light at 254 nm and by charring using ceric ammonium molybdate, p-anisaldehyde solutions. Chromatography was performed on a Teledyne ISCO CombiFlash Rf 200i using disposable silica cartridges. ^1^H, ^13^C and ^31^P NMR spectra were acquired on an OXFORD NMR AS500 (^1^H at 500.0 MHz, ^13^C at 126.0 MHz, ^31^P at 202.0 MHz). The residual solvent peaks were used as internal references and expressed in parts per million (ppm). Spin multiplicities were represented as s (singlet), d (doublet), t (triplet), quartet (q), dd (doublet of doublet), ddd (doublet of doublet of doublet), dt (doublet of triplet), m (multiplet) and broad singlet (br. s). Coupling constants (J) are given in Hertz. Mass spectra were recorded by either the Proteomics Facility at the Advanced Science Research Centre, CUNY or Novatia, LLC.

### Synthesis of indole coupled base modified nucleoside phosphoramidites

All phospnoramidites were synthesized using established synthetic methods, please see supplementary file for detailed synthetic methodologies, and characterization of all the intermediates and final phosphoramidites.

### Solid-phase synthesis of DNA sequences and purification

All DNA reagents required for the DNA synthesis were purchased from Glen Research. Wild-type DNA sequences were purchased from Integrated DNA Technologies. All DNA sequences were synthesized by following a standard solid-phase DNA synthesis protocols on an ABI394 DNA synthesizer (Biolytics) using a 0.2 µmol scale. The synthesis were performed in the standard mode using 2-cyanoethyl-N,N-diisopropylphosphoramidites. 0.1 M solution of each phosphoramidite and 0.25 M 5-Ethylthio-1-H-Tetrazole as an activator were used for the DNA synthesis. Iodine solution (0.02 M Iodine in THF/Py/Water) was used for oxidizing phosphoramidites. The reaction volume and duration for coupling natural phosphoramidites were 220 µL and 90 seconds, respectively, while for modified phosphoramidites, the reaction volume remained 220 µL, but the duration was extended to 600 seconds. The synthesized DNA sequences were cleaved from the solid-phase by treating with 30% aq. NH_4_OH (37 °C, 24 h, in dark). Deprotection was done according to base modification. DNA sequences were purified using high-performance liquid chromatography (Agilent) equipped with a C-18 reverse-phase column (Phenomenex) and UV detector with 0.1 M Triethylammonium Acetate (TEAA) and acetonitrile as a mobile phase. A UV-vis spectrophotometer was employed to quantify the purified DNA to obtain the stock concentration.

### CD spectral analysis

Circular dichroism (CD) measurements were conducted using a Jasco-1500 Circular Dichroism spectrometer. The data were collected using a quartz cuvette with a 10 mm pathlength, in the wavelength range of 220 nm to 330 nm, with a data pitch of 2 nm, an integration time of 8 seconds, a spectral bandwidth of 1 nm, and a scanning speed of 20 nm/min. Three accumulations were averaged for each spectrum. A blank measurement was first taken with 400 µL of PBS buffer. 200 pmol of DNA samples in 400 µL of PBS buffer were analyzed. Blank spectra were subtracted from the sample spectra, which were subsequently smoothed and zeroed at 320 nm.

### Tm Measurements

To analyze the thermal stability of the duplexes, a 50 nM of each duplexes (in 200 µL, Gibco™ 1X PBS, pH=7.4) were prepared incubating 10 µmol of fluorescent oligonucleotide sequence with 25 µmol of dabsyl modified complementary oligonucleotide sequence. Each duplex sample was placed into a reduced volume quartz cuvette (Horiba, path length=3mm), fluorescence intensity was measured using the FluoroMax Plus spectrofluorometer (Horiba) with a TC1 temperature controller (Quantum Northwest) and an EXT-440 liquid cooling system (Koolance). The emission wavelength utilized in the thermal stability assays correlated with fluoresceine (λ_em=514 nm), with the excitation wavelength at λex=488 nm, and slit width of 2.5 nm. Measurements were taken at 1 °C intervals (tolerance=±0.5°C), across a temperature range of 5 °C to 70 °C.

### Enzymatic Experiments

To investigate the impact of indole-modification in oligonucleotide sequences on their susceptibility to digestion by endonucleases, endonuclease digestion assays were carried out on the duplexes. For each assay, the 10 µmol duplex samples were prepared by incubating 10 µmol of fluorophore labelled oligonucleotide sequence with 25 µmol of quencher labelled complementary oligonucleotide sequence (sample volume: 10 µL). Prior to treatment with endonucleases, 2.5 µL of 6X reaction buffer, supplied by manufacturer respective to each enzyme was added (buffer Cf= 1X). After addition of respective endonuclease, the reaction samples were incubated for the specified amount of time in a TC 9639 Thermal Cycler (Benchmark). For DNase I digestion assays, the duplex samples were treated with 1.4 units of DNase I (Thermoscientific, REF# EN0521, 0.56 u/µL). The reactions were incubated for 15 minutes at 37 °C. For EcoRI digestion assays, the duplex samples were treated with 70 units of EcoRI-HF (NEB, R3101M, 28 u/µL). The reactions were incubated overnight in a thermocycler at 37 °C. For XmaI digestion assays, the duplex samples were treated with 7 units of XmaI (NEB, R0180S, 2.8 u/µL). The reactions were incubated overnight at 37 °C. For SmaI digestion assays, the duplex samples were treated with 7 units of SmaI (Thermoscientific, REF# ER0665, 2.8 u/µL). The reactions were incubated overnight in a thermocycler at 25 °C. To assess enzymatic digestions in all endonuclease digestion assays, the treated duplex samples were diluted with 185 µL of 1X PBS (Duplex Cf= 50 nM) post-incubation, and fluorescence measurement was performed at 20 °C. The emission wavelength range utilized for measuring treated duplex samples was, λ_em=500 to 600 nm, and the excitation wavelength was λex=488 nm, with a 2.5nm slit width. An unmodified, wild-type duplex acted as the control, and was measured alongside each modified duplex, to ensure consistency in conditions between each experiment. Heat maps were constructed using fluorescence intensities recorded for each duplexes normalizing against that of duplex 1; WT:cWT.

## Results

### Synthesis of ZImera duplexes

Using propyl- and propargyl-linked indole, we synthesized corresponding nucleoside phosphoramidites through a classical Sonogashira cross-coupling, followed by hydrogenation, DMT protection and phosphoramidite synthesis to generate compounds **1a**-**4a**. Compounds **1b**-**4b** were synthesized using the same approach without hydrogenation (**Scheme 1**).

**Scheme 1:**
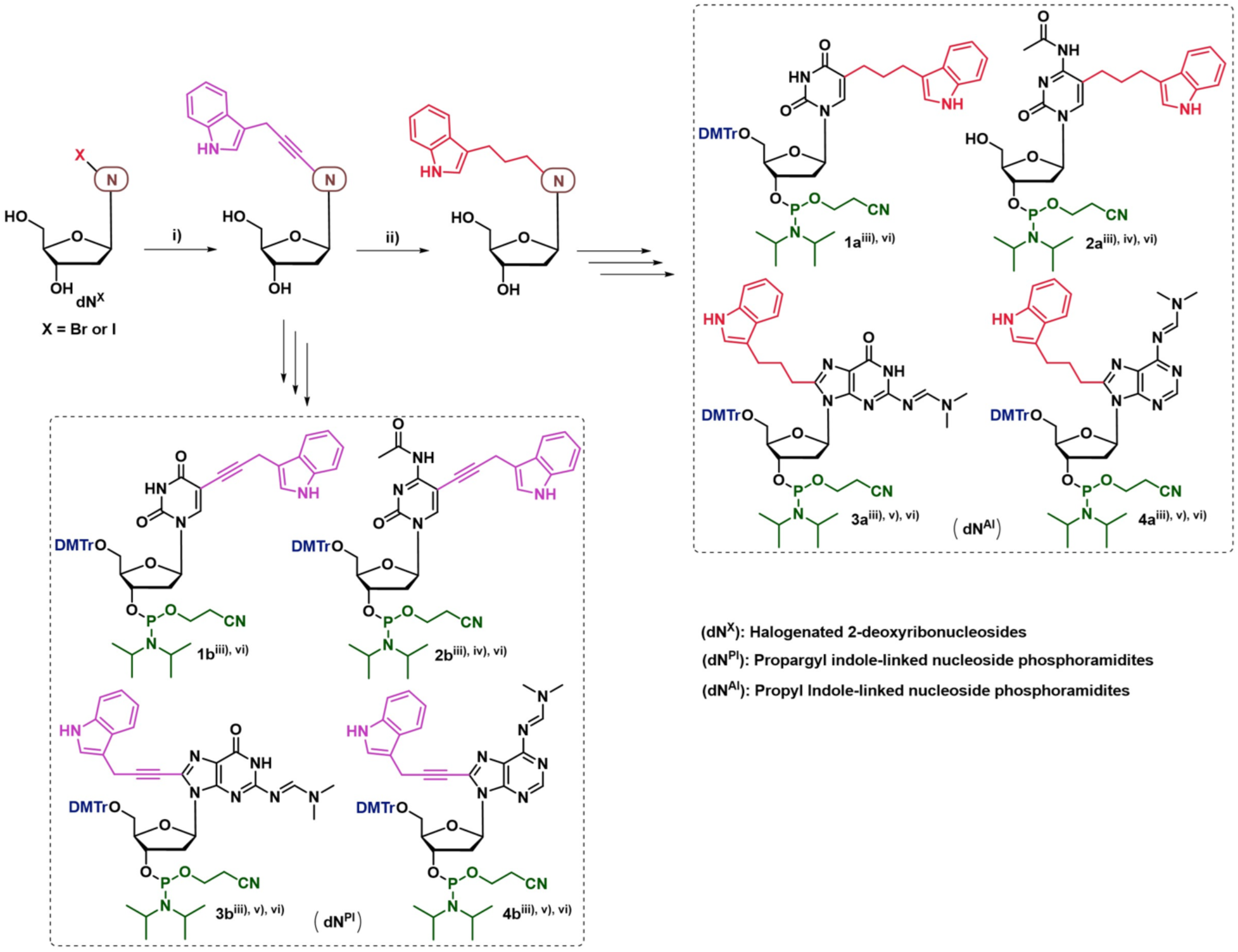
Design and synthesis of propyl and propargyl-linked indole modified nucleoside phosphoramidites Reagents and conditions: (i) 3-Prop-2-ynyl-indole, Pd(PPh_3_)_4_, CuI, Et_3_N, DMF, 60 °C, 2-3 h; ii) H_2_, 10% Pd/C, MeOH, 50 °C, overnight; (iii) DMTrCl, DMAP, pyridine, rt, overnight; (iv) Ac_2_O, DMF, 0 °C to rt, 8-10 h; (v) DMF-DMA, DMF, rt, 3-4 h; (vi) 2-cyanoethyl-*N,N*-diisopropylchlorophosphoramidite, DIPEA, DCM, 0 °C to rt, 1.0-1.5 h.

We used standard solid-state chemistry on an ABI394 DNA synthesizer to synthesize the short oligos listed in **Table 1**, using coupling times extended to 600 seconds to couple **1a/b**, **2a/b**, **3a/b**, and **4a/b**. We did not observe any reduction in coupling efficiency for propyl-linked indole-modified nucleoside phosphoramidites **1a**, **2a**, **3a**, and **4a**. Based on DMT detection, the overall yield of Z1-Z6 and cZ1-cZ5 oligos was approximately 80%-85%, the same yield as that of comparable WT sequences, suggesting that modification at C-5 and C-8 using flexible propyl linkage does not impact coupling efficiency. However, we observed a drastic reduction in the coupling efficiency of propargyl-linked indole nucleoside phosphoramidites (**1b**, **2b**, **3b**, and **4b**), reducing the overall yield to 15%-20% for Z11-Z55 oligonucleotides, respectively. The analysis of each oligonucleotides using mass spectrometry (**Table S1**) confirmed successful synthesis and purification. Hocek and others observed similar coupling efficiencies for nucleoside phosphoramidite derivatives in the synthesis of hypermodified DNA oligos, suggesting that rigidity of the linker might play a role in orienting the activated 3’ amidite moiety during the DNA synthesis.^37^

**Table 1:**
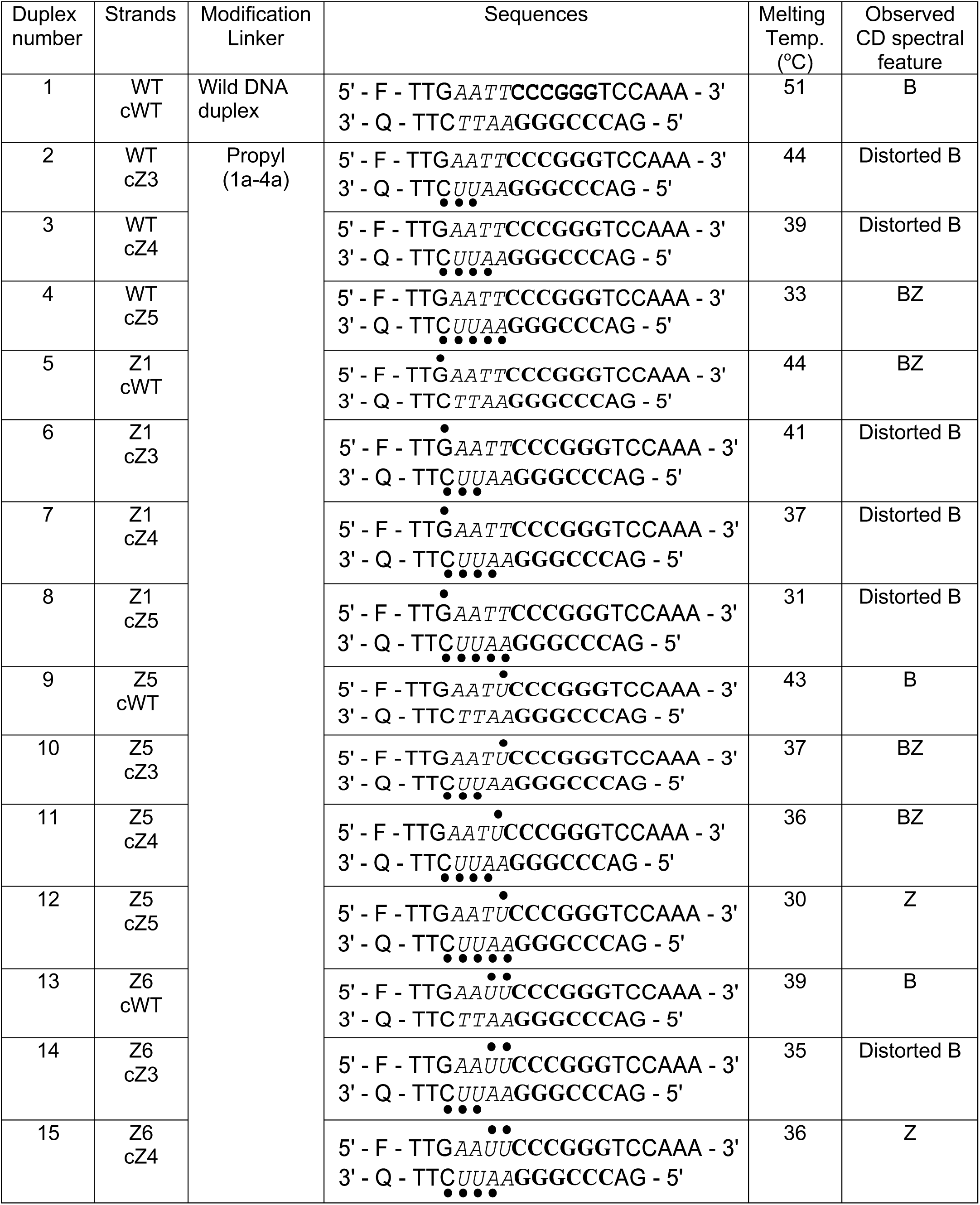
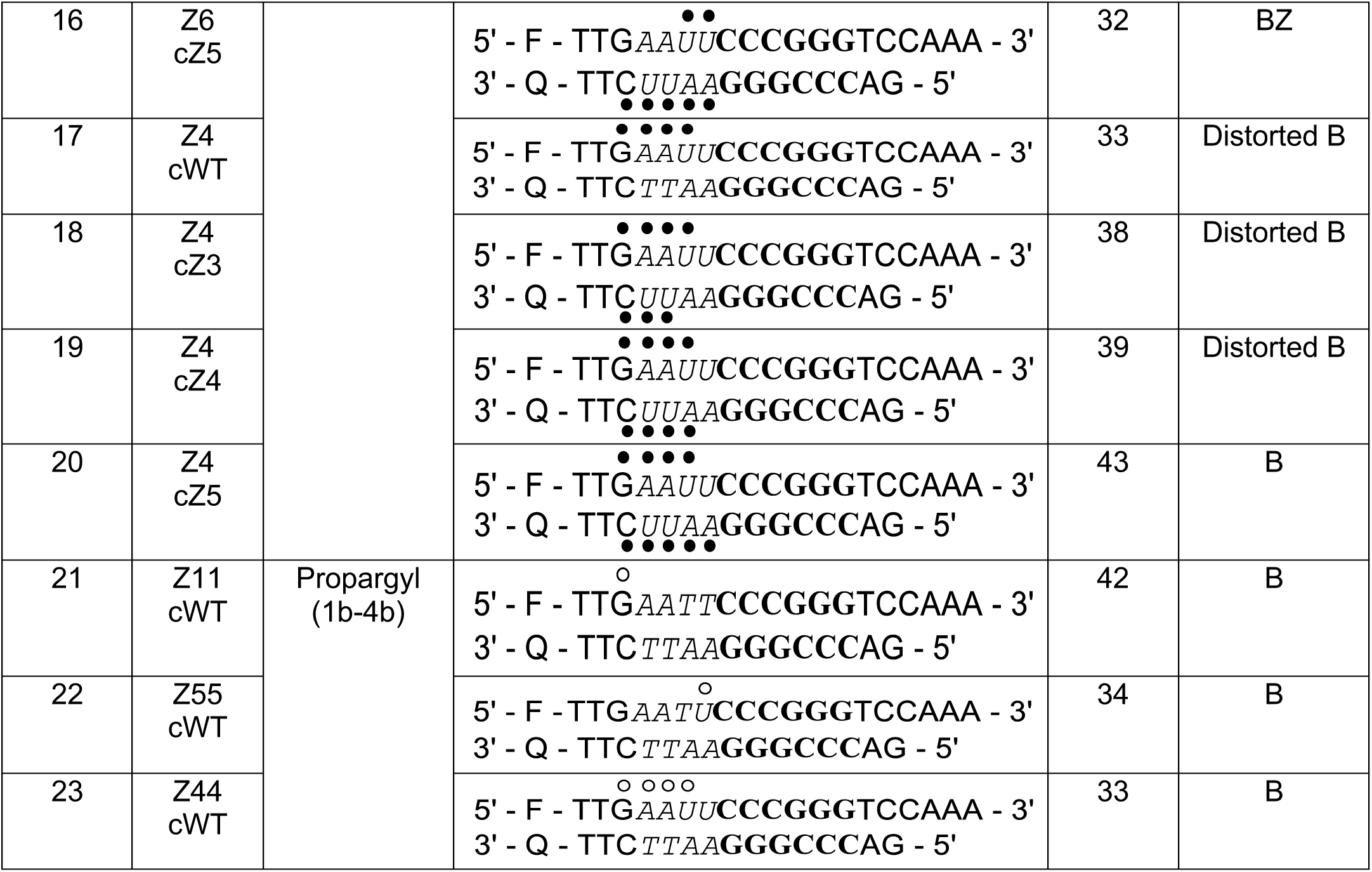
List of oligonucleotides sequences: solid circles (•): 3-propyl-indole linked nucleotide; open circles (○): 3-(prop-2-yn-1-yl)-indole linked nucleotide; F: 5’-(6-FAM)-labeled; Q: 3’-(Dabcyl)-labeled. *`Italics`* EcoR1 restriction site; **bold** Xmal, Smal restriction site.

### Thermal stability and CD spectral features of ZImera

Since modifications at the C-5 and C-8 positions of nucleobases are known to destabilize duplexes, we first compared the annealing behavior at physiological temperature and the corresponding melting temperatures of modified duplexes to those of the wild-type duplex (**Table 1, Figure S1-S2**). The wild-type duplex exhibited a melting temperature of 51 °C, while duplexes containing indole modifications showed reductions in melting temperatures. All duplexes with one to three cytidine and/or uridine modifications show a drop in melting by 6 °C. For instance, duplex 2 demonstrated a decrease in melting temperature by 7 °C compared to the wild-type duplex, with one cytidine and two uridine modifications. The melting temperature further decreased by approximately 5 °C for each additional modified adenine introduced in duplexes 3 and 4. Interestingly, the conformation of the duplexes (2-4) also transitions from B to BZ as the number of base modifications increases (**Table 1**, **Figure 1 B-D**). This trend persisted across duplexes 5-8 and 9-12, indicating that the incorporation of uridine and adenine modifications significantly affects the melting temperature and conformation (**Table 1**, **Figure 1 E-L, Figure S3**). Notably, duplexes 8, 12, 16 and 17 exhibited the lowest melting temperatures, further supporting the idea that propyl-modified uridine acts as a destabilizer. When two modified uridines are adjacent in duplexes (duplexes 13–16), those duplexes exhibit the lowest melting temperatures and adopt more distorted conformations (**Table 1**). Additional adenine and guanine modifications did not lead to substantial reductions in melting temperature. Furthermore, in duplexes 5, 9, 13, and 17, which contain only one modified strand, the decrease in melting temperature correlated more strongly with the incorporation of modified uridine than with modified guanine. These findings may indicate that propyl-modified uridine contributes more significantly to duplex destabilization than propyl modified guanine. Overall, the melting temperature is most strongly impacted when the CUUAA sequence is modified with propyl-linked indole. This trend is evident in duplexes 6 to 8, 10 to 12, 14 to 16, and 18 to 19. Interestingly, duplex 20, which has the highest number of modifications, exhibited an unexpectedly higher melting temperature. This suggests that stability of the duplex 20 is primarily governed by Watson-Crick base pairing and the flexibility of propyl linker likely enhances base stacking on both sides of the duplex, promoting the adoption of a B-conformation. In propargyl-linked duplexes, the melting temperature consistently decreased by 8-7 °C as the number of modified uridines and adenines increased suggesting that the linker rigidity destabilizes the duplex. (**Table 1, Figures S1-S2**). We investigated the structural features of ZImera using circular dichroism (CD) studies (Figure 1). As anticipated, the wild-type duplex displayed a B-form conformation. However, with an increasing number of base modifications in duplexes (2– 4), the conformation transitioned from a distorted B-form to a BZ-form. A similar trend was observed in duplexes 6–8, 9–12, and 13–16 (**Figure 1**), where the conformation shifted from B-form to Z-form. Interestingly, duplexes with the highest number of base modifications (17–20) maintained a regular B-form (**Figure 1T**). Notably, similar to melting point deviations, the transition to the B-conformation observed in CD spectra correlates with A:U modifications, which may play a key role in governing conformational adaptation.

**Figure 1.**
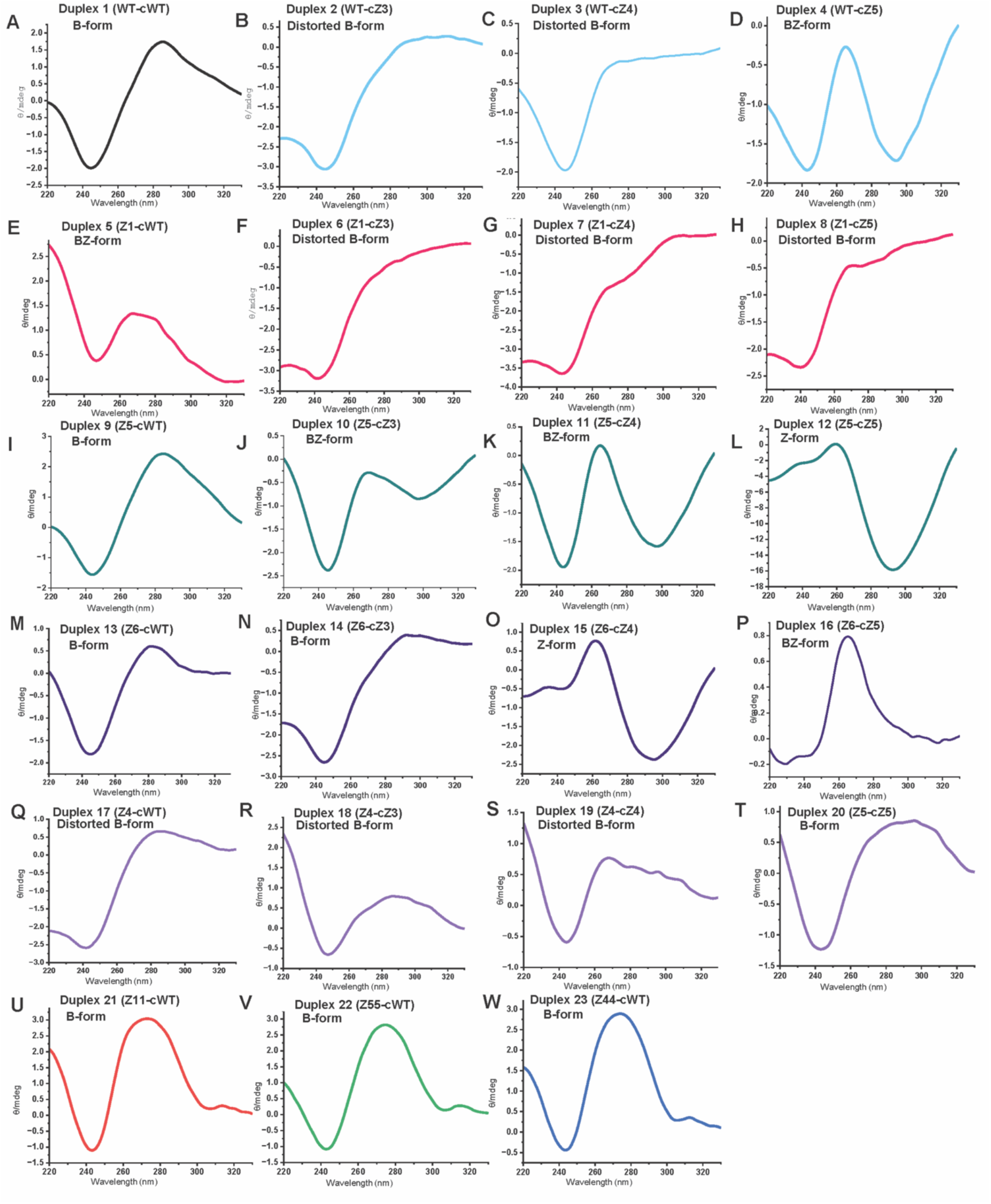
CD spectra of ZImera: CD spectra of ZImera were recorded at 25 °C using a 10 mm pathlength quartz cuvette in the range of 220 nm to 330 nm, with a scanning speed of 20 nm/min, 1 nm bandwidth, 8s response time, and three accumulations.

### Restriction digestion of ZImera

Restriction-modification (R-M) enzymes show both nuclease and methylation activities that resemble primitive immune systems that destroy foreign DNA in bacteria.^38^ R-M enzymes are programmed to protect host chromosomes by recognizing and cutting foreign DNA from invading viruses. Given their important role in biology, we designed the sequences listed in **Table 1** with EcoRI, Xmal, and Smal restriction enzymatic cleavage sites and strategically incorporated modified **1a-d** and **2a-d** directly on EcoRI restriction site and juxtaposed to XmaI/SmaI restriction site to explore how conformational adaptation of a duplex, owing to these modifications, could lead to NCS and how the resulting NCS could impact the specificity and promiscuity of the duplex towards restriction enzymes and DNase 1.

First, we assessed the sensitivity of the duplexes towards DNase 1 cleavage (**Figure 2A-B, Figure S3**). Remarkably, despite the known high promiscuity of DNA duplexes towards nuclease activity mediated by DNase1, we observed that sensitivity of the modified duplexes towards DNase1 varies based on the position of the modification. For example, DNase1 cleaves duplex 2 in a manner similar to that of wild-type duplex sequence, suggesting that the modifications do not interfere with sequence promiscuity towards DNase1. Circular dichroism (CD) analysis revealed that the duplex 2 forms a distorted B-DNA structure (**Table 1**, **Figure 1B**). Therefore, despite the modifications introduced in cZ3, these results suggest that a distorted B-DNA structure could potentially make the backbone of duplex 2 more accessible to DNase1 cleavage, in turn resulting in a more promiscuous duplex. However, the sensitivity towards DNase1 gradually increases when modified A introduced for duplex 3 and duplex 4 (**Figure 2A, first column, Figure S3**).

**Figure 2.**
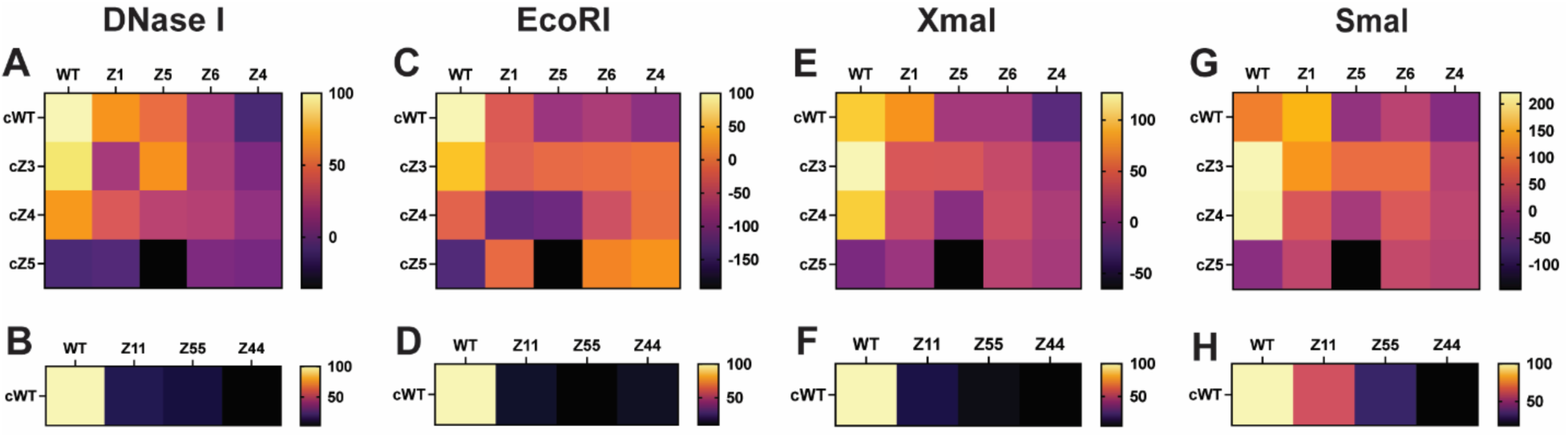
Heatmap showing a comparative analysis of sequence promiscuity towards DNase 1 (A-B), EcoRI (C-D), XmaI (E-F), and SmaI (G-H). Enzymatic assays were performed using FRET assays with a Fluorophore-labeled sense strand and a quencher-labeled anti-sense strand. Each duplex was incubated overnight at 37 °C with corresponding enzyme, and enzymatic activity was analyzed using FRET at 20 °C using a fluorescence spectrometer. Fluorescence intensity measured was normalized against the control WT:cWT

The nuclease sensitivity was observed for duplexes 5, 9 13, 17 formed with Z1, Z5, Z6 and Z4, shows that nuclease resistance increases with higher modifications and that this sensitivity is independent of the observed conformational adaptation (**Figure 2A, first row; Figure S3**). DNase I is an endonuclease that nonspecifically, but preferentially, cleaves double-stranded B-DNA at 3’ hydroxyl and 5’ phosphoryl nucleotides using a single-strand nicking mechanism. The significant nuclease resistance observed in duplex 4 and duplex 17, with only 4 to 5 modifications introduced into a 15-mer duplex, may be attributed to the loss of minor grooves resulting from a BZ transition or a distorted B-conformation (**Figures 1D and 1Q**, **Figure 2A**). Furthermore, duplexes 17, 18, 19, 20, 12, 8, and 4 containing Z4 and cZ5 (**Figure 2A-4^th^ row and Figure 2A-5^th^ column**) exhibited the highest resistance to DNase I. In Z4, the GAAU segment was modified with propyl-linked indole, seems to contribute to DNase I resistance. Similarly, in cZ5, modifications to the CUUAA segment also resulted in increased resistance to DNase I. The resistance to DNase1 was profound when oligos were modified with **1b**-**4b** suggesting that propargyl linker induces more rigid structures, thus, despite the observed B-conformation (**Figure 1 U-W**), and rigid modification introduced to the strand compromises duplex specificity towards DNase1. However, the duplex 21 (Z11:cWT) shows a less resistance compared to that of duplex 22 (Z55:cWT) and duplex 23 (Z44:cWT), which mimic a pattern similar to that of the propyl modified series (**Figure 2B**). The duplexes containing Z6 (**Figure 2A-4^th^ column**) significantly deviate in melting points and exhibit greater deviations from the B-DNA structure. These duplexes also show higher DNase I resistance, suggesting that the strategic incorporation of propyl-linked indole purines and pyrimidines can lead to increased DNase I sensitivity.

Second, we assessed endonuclease EcoRI activity towards the duplexes (**Figure 2 C-D, Figure S4**). EcoRI is known to first scan DNA duplexes in one dimension to find its cleavage site and then cleave between G and A in a GAATTC palindrome.^38^ A 2D NMR analysis of dodecamer structure with an EcoRI site showed that structural anomalies, such as kinks or increased flexibility of a duplex, facilitate scission. This finding suggests that modification at the EcoRI site, which leads to a distorted B conformation, may have played a role in increasing the promiscuity of duplex 2 (WT:cZ3) towards EcoRI, whereas, distorted B-forming duplex 3 (WT:cZ4) and BZ forming duplex 4 (WT:cZ5) shows higher resistivity towards EcoRI.^38^ Duplexes 5, 9, 13, and 17 containing modified restriction site Z1, Z5, Z6 and Z4 show a gradual increase in resistance to EcoRI (**Figure 2C-1^st^ row**). This resistance is expected, as the modifications are introduced directly at the restriction site. However, unexpectedly, the duplexes with fully modified EcoRI site on one or both strands (**Figure 2C-4^th^ row and 5^th^ column**) showed variable resistance to ECoRI. For example, duplex 20 forms a B conformation (**Figure 1T**) and showed promiscuity towards ECoRI despite the presence of modified segments directly on cutting site 5’GAAU3’ or 3’CUUAA5’ suggesting that conformational change is what governs the ECoRI promiscuity not the modification when indole linked with propyl linker. Similar to DNase1, resistivity of the duplex is increased when the sense strand was modified with **1b**-**4b**, since all sequences resisted EcoRI activity likely resulting from steric hindrance introduced by the indole moiety (**Figure 2D**). This line of evidence suggests that the modification induced steric effects by the indole moiety leading to abolished interaction between the duplex and EcoRI.

Third, we explored how a modification introduced to a juxtaposition to a restriction site would impact scission (**Figure 2E-F, 2G-H, Figure S5, Figure S6**). The promiscuity of the duplex towards Smal and Xmal also depends on modification position, even though the modifications are not directly on the Smal and XmaI cleavage site. For example, both Smal and XmaI cleave duplex 2 (WT:cZ3) and duplex 3 (WT:cZ4) to a greater extent than the WT duplex 1, while duplex 4 (WT: cZ5) show no promiscuity towards Xmal or SmaI. Smal and Xmal are isoschizomers, i.e., restriction enzymes that recognize the same DNA sequence, such as CCCGGG, but may not cut at the same location.^39^ Xmal cleaves between the external cytosines, while Smal cleaves between CG in the CCCGGG to introduce blunt end scission.^39^ It has been shown that both endonucleases form a stable, specific complex with DNA. However, the landing of endonuclease on the duplex leads to the bending of DNA such that Smal bends the DNA towards the major groove, similar to EcoRI, while Xmal bends the DNA towards the minor groove.^39^ Our findings strongly suggest that the induction of NCS in response to hybridization with complementary strand containing C-5 and C-8 modified nucleobases leads to more promiscuity of duplex towards Xmal and Smal than that towards WT duplex. With the exception of duplexes 2 and 3, all other duplexes exhibit modest to high resistance to XmaI and SmaI. This resistance occurs despite the modifications being positioned adjacent to the restriction site, suggesting that downstream modifications influence upstream restriction sites through conformational adaptation of the duplex in response to the modified bases. Additionally, the formation of BZ conformation in duplexe 5 (**Figure 1E**) appears to increase their susceptibility to XmaI and SmaI. This promiscuity is notably higher than that observed in the wild-type duplex 1, which adopts a standard B-form, potentially because of the more exposed DNA backbone facilitates efficient cleavage. Despite forming a B-DNA conformation, a profound loss of sensitivity towards nucleases was observed for duplexes modified with **1b**, **2b**, **3b**, and **4b**, suggesting that rigidity of the indole moiety impacted the duplex structure more strongly when a propargyl linker was used. All duplexes 21, 22 and 23, containing propargyl linked indole, Z11:cWT, Z55:cWT, and Z44:cWT, were resistant to DNase1, EcoRI, and Xmal, despite the extent or site of modification. However, the promiscuity of these duplexes towards SmaI slightly deviates from this trend. For example, Z11:cWT3 shows lower resistance to SmaI, and the resistivity gradually increased with the increasing number of modifications.

## Discussion

Here, we demonstrate that strategically positioning nucleotides linked to indole via a flexible propyl linker can fine-tune sequence promiscuity toward nucleases and restriction enzymes. Specifically, we investigated two modifications: one at the C-5 position of pyrimidine and another at the C-8 position of purine in native DNA. Our findings reveal that the rigidity of the linker tethering the bulky substituent at these positions plays a critical role in determining sequence promiscuity toward nucleases. For flexible propyl linkers, promiscuity toward restriction enzymes was particularly variable. Notably, we found that introducing modifications directly at or near a restriction site impacts nuclease cleavage, with promiscuity influenced by downstream conformational changes resulting from upstream promiscuity. For example, all duplexes resisted cleavage by DNase I, EcoRI, SmaI, and XmaI when the sense strand was modified with C-5 propargyl indole-linked pyrimidines (U, C) and C-8 propargyl indole-linked purines (A, G). In contrast, sequence promiscuity varied when the complementary strand was modified with C-5 propyl indole-linked pyrimidines and C-8 propyl indole-linked purines, depending on the position and extent of the modification.

As the understanding of noncanonical DNA structures (NCS) grows, there is a pressing need for molecular tools to expand this knowledge and its applications. Confirmation of DNA structure by CD spectroscopy is well established. For example, left-handed Z-DNA, an unusual form of DNA structure, was first identified using CD spectroscopy, demonstrating the accuracy of this technique in clearly depicting structural deviations in DNA. While more detailed structural insights into conformational adaptation are currently underway, our study provides a preliminary molecular framework for inducing non-canonical structures (NCS), such as Z-DNA and BZ-DNA, in native DNA strands. The chimera (ZImera) comprising C-5 indole-linked pyrimidines and C-8 indole-linked purines exhibits unique biochemical properties centered on conformational adaptation, enabling the induction of NCS in native DNA. The resulting NCS exposes specific sites, increasing sequence promiscuity toward site-specific restriction nucleases and nonspecific nucleases such as DNase I. Future approaches could focus on inducing excluded sites for precise cleavage by base-editing enzymes and DNA repair enzymes. Since many nucleases, DNA/RNA-binding proteins, and cellular machinery preferentially bind B- and A-conformations or single-stranded DNA/RNA, the demonstrated conformational adaptation in ZImera may provide a platform for designing novel antisense therapeutic approaches.

We envision a new era of NCS-based antisense therapeutic strategies, where noncoding therapeutics capable of inducing NCS can be used as a therapeutic modality. The ZImera described here could also serve as a tool for screening small molecules, such as Z-disruptors, Z-DNA binders, and BZ inhibitors, under physiological conditions (e.g., PBS) without requiring specific sequences. Additionally, Z-conformation-binding proteins like ADAR could be selectively targeted using ZImera tethered to PROTAC for protein degradation strategies as previously shown.^40^ Modified duplexes could also be employed to design DNA materials with expanded chemical functionality and nuclease resistance. In conclusion, this study demonstrates, for the first time, that complementary strands modified with indole-linked pyrimidines and purines via flexible or rigid linkers can be used to target native nucleic acid sequences to induce distorted B, Z, or BZ conformations. This conformational adaptation controls sequence promiscuity toward digestion enzymes, offering a novel strategy for developing molecular tools to study NCS and create innovative nucleic acid-based therapeutics.

## Acknowledgments

We gratefully acknowledge the support of the NIGMS grant R35GM139336, which made the research described in this manuscript possible. We also extend our heartfelt thanks to Steve Benner and Pete Gannett for their critical reading of the manuscript.

## References

(1) Wang, G.; Vasquez, K. M. Dynamic alternative DNA structures in biology and disease. Nat Rev Genet 2023, 24 (4), 211–234. DOI: 10.1038/s41576-022-00539-9 From NLM Medline.

(2) Rich, A.; Zhang, S. Timeline: Z-DNA: the long road to biological function. Nat Rev Genet 2003, 4 (7), 566–572. DOI: 10.1038/nrg1115 From NLM Medline.

(3) Choi, J.; Majima, T. Conformational changes of non-B DNA. Chem Soc Rev 2011, 40 (12), 5893–5909. DOI: 10.1039/c1cs15153c From NLM Medline.

(4) Quyen, D. V.; Ha, S. C.; Lowenhaupt, K.; Rich, A.; Kim, K. K.; Kim, Y. G. Characterization of DNA-binding activity of Z alpha domains from poxviruses and the importance of the beta-wing regions in converting B-DNA to Z-DNA. Nucleic Acids Res 2007, 35 (22), 7714–7720. DOI: 10.1093/nar/gkm748 From NLM Medline.

(5) Wieg, B.; Wolfl, S.; Dorbic, T.; Vahrson, W.; Rich, A. Transcription of human c-myc in permeabilized nuclei is associated with formation of Z-DNA in three discrete regions of the gene. EMBO J 1992, 11 (12), 4653–4663. DOI: 10.1002/j.1460-2075.1992.tb05567.x From NLM Medline.

(6) Wang, G.; Vasquez, K. M. Impact of alternative DNA structures on DNA damage, DNA repair, and genetic instability. DNA Repair (Amst) 2014, 19, 143–151. DOI: 10.1016/j.dnarep.2014.03.017 From NLM Medline.

(7) Fogg, J. M.; Randall, G. L.; Peeh, B. M.; Sumners, W. L.; Harris, S. A.; Zechiedrich, L. Bullied no more: when and how DNA shoves proteins around. Q Rev Biophys 2012, 45 (3), 257–299. DOI: 10.1017/S0033583512000054 From NLM Medline.

(8) Ravichandran, S.; Subramani, V. K.; Kim, K. K. Z-DNA in the genome: from structure to disease. Biophys Rev 2019, 11 (3), 383–387. DOI: 10.1007/s12551-019-00534-1 From NLM PubMed-not-MEDLINE.

(9) Smirnov, E.; Molinova, P.; Chmurciakova, N.; Vacik, T.; Cmarko, D. Non-canonical DNA structures in the human ribosomal DNA. Histochem Cell Biol 2023, 160 (6), 499–515. DOI: 10.1007/s00418-023-02233-1 From NLM Medline.

(10) Htun, H.; Dahlberg, J. E. Topology and formation of triple-stranded H-DNA. Science 1989, 243 (4898), 1571–1576. DOI: 10.1126/science.2648571 From NLM Medline.

(11) Zhang, F.; Huang, Q.; Yan, J.; Chen, Z. Histone Acetylation Induced Transformation of B-DNA to Z-DNA in Cells Probed through FT-IR Spectroscopy. Anal Chem 2016, 88 (8), 4179–4182. DOI: 10.1021/acs.analchem.6b00400 From NLM Medline.

(12) Duval-Valentin, G.; de Bizemont, T.; Takasugi, M.; Mergny, J. L.; Bisagni, E.; Helene, C. Triple-helix specific ligands stabilize H-DNA conformation. J Mol Biol 1995, 247 (5), 847–858. DOI: 10.1006/jmbi.1995.0185 From NLM Medline.

(13) Rich, A.; Nordheim, A.; Wang, A. H. The chemistry and biology of lek-handed Z-DNA. Annu Rev Biochem 1984, 53, 791–846. DOI: 10.1146/annurev.bi.53.070184.004043 From NLM Medline.

(14) Buzzo, J. R.; Devaraj, A.; Gloag, E. S.; Jurcisek, J. A.; Robledo-Avila, F.; Kesler, T.; Wilbanks, K.; Mashburn-Warren, L.; Balu, S.; Wickham, J.;, et al. Z-form extracellular DNA is a structural component of the bacterial biofilm matrix. Cell 2021, 184 (23), 5740–5758 e5717. DOI: 10.1016/j.cell.2021.10.010 From NLM Medline.

(15) Kim, Y. G.; Muralinath, M.; Brandt, T.; Pearcy, M.; Hauns, K.; Lowenhaupt, K.; Jacobs, B. L.; Rich, A. A role for Z-DNA binding in vaccinia virus pathogenesis. Proc Natl Acad Sci U S A 2003, 100 (12), 6974–6979. DOI: 10.1073/pnas.0431131100 From NLM Medline.

(16) Wong, B.; Chen, S.; Kwon, J. A.; Rich, A. Characterization of Z-DNA as a nucleosome-boundary element in yeast Saccharomyces cerevisiae. Proc Natl Acad Sci U S A 2007, 104 (7), 2229–2234. DOI: 10.1073/pnas.0611447104 From NLM Medline.

(17) Arndt-Jovin, D. J.; Udvardy, A.; Garner, M. M.; Riher, S.; Jovin, T. M. Z-DNA binding and inhibition by GTP of Drosophila topoisomerase II. Biochemistry 1993, 32 (18), 4862–4872. DOI: 10.1021/bi00069a023 From NLM Medline.

(18) Vasudevaraju, P.; Bharathi; Garruto, R. M.; Sambamurti, K.; Rao, K. S. Role of DNA dynamics in Alzheimer’s disease. Brain Res Rev 2008, 58 (1), 136–148. DOI: 10.1016/j.brainresrev.2008.01.001 From NLM Medline.

(19) Wells, R. D. Non-B DNA conformations, mutagenesis and disease. Trends Biochem Sci 2007, 32 (6), 271–278. DOI: 10.1016/j.tibs.2007.04.003 From NLM Medline.

(20) Wang, G.; Christensen, L. A.; Vasquez, K. M. Z-DNA-forming sequences generate large-scale deletions in mammalian cells. Proc Natl Acad Sci U S A 2006, 103 (8), 2677–2682. DOI: 10.1073/pnas.0511084103 From NLM Medline.

(21) Portela, A.; Esteller, M. Epigenetic modifications and human disease. Nat Biotechnol 2010, 28 (10), 1057–1068. DOI: 10.1038/nbt.1685 From NLM Medline.

(22) Behe, M.; Felsenfeld, G. Effects of methylation on a synthetic polynucleotide: the B--Z transition in poly(dG-m5dC).poly(dG-m5dC). Proc Natl Acad Sci U S A 1981, 78 (3), 1619–1623. DOI: 10.1073/pnas.78.3.1619 From NLM Medline.

(23) Zacharias, W.; Jaworski, A.; Wells, R. D. Cytosine methylation enhances Z-DNA formation in vivo. J Bacteriol 1990, 172 (6), 3278–3283. DOI: 10.1128/jb.172.6.3278-3283.1990 From NLM Medline.

(24) Koeris, M.; Funke, L.; Shrestha, J.; Rich, A.; Maas, S. Modulation of ADAR1 editing activity by Z-RNA in vitro. Nucleic Acids Res 2005, 33 (16), 5362–5370. DOI: 10.1093/nar/gki849 From NLM Medline.

(25) Liu, J.; Wang, F.; Zhang, Y.; Liu, J.; Zhao, B. ADAR1-Mediated RNA Editing and Its Role in Cancer. Front Cell Dev Biol 2022, 10, 956649. DOI: 10.3389/fcell.2022.956649 From NLM PubMed-not-MEDLINE.

(26) Jiao, Y.; Xu, Y.; Liu, C.; Miao, R.; Liu, C.; Wang, Y.; Liu, J. The role of ADAR1 through and beyond its editing activity in cancer. Cell Commun Signal 2024, 22 (1), 42. DOI: 10.1186/s12964-023-01465-x From NLM Medline.

(27) Hubbard, N. W.; Ames, J. M.; Maurano, M.; Chu, L. H.; Somfleth, K. Y.; Gokhale, N. S.; Werner, M.; Snyder, J. M.; Lichauco, K.; Savan, R.;, et al. ADAR1 mutation causes ZBP1-dependent immunopathology. Nature 2022, 607 (7920), 769–775. DOI: 10.1038/s41586-022-04896-7 From NLM Medline.

(28) Barraud, P.; Allain, F. H. ADAR proteins: double-stranded RNA and Z-DNA binding domains. Curr Top Microbiol Immunol 2012, 353, 35–60. DOI: 10.1007/82_2011_145 From NLM Medline.

(29) Thomsen, N. M.; Vongsutilers, V.; Ganneh, P. M. The synthesis of C8-aryl purines, nucleosides and phosphoramidites. Crit Rev Eukaryot Gene Expr 2011, 21 (2), 155–176. DOI: 10.1615/critreveukargeneexpr.v21.i2.50 From NLM Medline.

(30) Egli, M.; Williams, L. D.; Gao, Q.; Rich, A. Structure of the pure-spermine form of Z-DNA (magnesium free) at 1-A resolution. Biochemistry 1991, 30 (48), 11388–11402. DOI: 10.1021/bi00112a005 From NLM Medline.

(31) Thomas, T. J.; Thomas, T. Polyamine-induced Z-DNA conformation in plasmids containing (dA-dC)n.(dG-dT)n inserts and increased binding of lupus autoantibodies to the Z-DNA form of plasmids. Biochem J 1994, 298 *( Pt* *2*) (Pt 2), 485–491. DOI: 10.1042/bj2980485 From NLM Medline.

(32) Xu, Y.; Ikeda, R.; Sugiyama, H. 8-Methylguanosine: a powerful Z-DNA stabilizer. J Am Chem Soc 2003, 125 (44), 13519–13524. DOI: 10.1021/ja036233i From NLM Medline.

(33) El-Khoury, R.; Cabrero, C.; Movilla, S.; Kaur, H.; Friedland, D.; Dominguez, A.; Thorpe, J. D.; Roman, M.; Orozco, M.; Gonzalez, C.;, et al. Formation of lek-handed helices by C2’-fluorinated nucleic acids under physiological salt conditions. Nucleic Acids Res 2024, 52 (13), 7414–7428. DOI: 10.1093/nar/gkae508 From NLM Medline.

(34) Stempel, E.; Gaich, T. Cyclohepta[b]indoles: A Privileged Structure Motif in Natural Products and Drug Design. Acc Chem Res 2016, 49 (11), 2390–2402. DOI: 10.1021/acs.accounts.6b00265 From NLM Medline.

(35) Lai, J. S.; Kool, E. T. Selective pairing of polyfluorinated DNA bases. J Am Chem Soc 2004, 126 (10), 3040–3041. DOI: 10.1021/ja039571s From NLM Medline.

(36) Escude, C.; Nguyen, C. H.; Mergny, J.-L.; Sun, J.-S.; Bisagni, E.; Garestier, T.; Helene, C. Selective Stabilization of DNA Triple Helixes by Benzopyridoindole Derivatives. Journal of the American Chemical Society 1995, 117 (41), 10212–10219. DOI: 10.1021/ja00146a006.

(37) Jestrabova, I.; Postova Slavetinska, L.; Hocek, M. Arylethynyl- or Alkynyl-Linked Pyrimidine and 7-Deazapurine 2’-Deoxyribonucleoside 3’-Phosphoramidites for Chemical Synthesis of Hypermodified Hydrophobic Oligonucleotides. ACS Omega 2023, 8 (42), 39447–39453. DOI: 10.1021/acsomega.3c05202 From NLM PubMed-not-MEDLINE.

(38) Heitman, J. How the EcoRI endonuclease recognizes and cleaves DNA. Bioessays 1992, 14 (7), 445–454. DOI: 10.1002/bies.950140704 From NLM Medline.

(39) Withers, B. E.; Dunbar, J. C. The endonuclease isoschizomers, SmaI and XmaI, bend DNA in opposite orientations. Nucleic Acids Res 1993, 21 (11), 2571–2577. DOI: 10.1093/nar/21.11.2571 From NLM Medline.

(40) Wang, Z.; Zhang, D.; Qiu, X.; Inuzuka, H.; Xiong, Y.; Liu, J.; Chen, L.; Chen, H.; Xie, L.; Kaniskan, H. U.;, et al. Structurally Specific Z-DNA Proteolysis Targeting Chimera Enables Targeted Degradation of Adenosine Deaminase Acting on RNA 1. J Am Chem Soc 2024, 146 (11), 7584–7593. DOI: 10.1021/jacs.3c13646 From NLM Medline.

