## Supporting information for "The modification of DNA with indole-linked nucleotides alters its sensitivity to enzymatic cleavage"

| S. No | Table of contents | Page No. |
|---|---|---|
| 1 | Synthesis of indole coupled base modified nucleoside phosphoramidites | S3 |
| 2 | Table 1. List of base modified DNA sequences, their calculated and measured masses | S24 |
| 3 | Figure S1-S2 Thermal analysis of ZImera | S25 |
| 4 | FRET assays:<br>Figure S3 Analysis of DNase 1 activity of ZImera<br>Figure S4 Analysis of EcoRI activity of ZImera<br>Figure S5 Analysis of XmaI activity of ZImera<br>Figure S6 Analysis of SmaI activity of ZImera | S27 |
| 5 | Copies NMR Spectra | S29 |
| 6 | Mass Spectra of DNA sequences | S63 |
| 7 | Reference | S67 |


### **1.** Synthesis of indole coupled base modified nucleoside phosphoramidites:

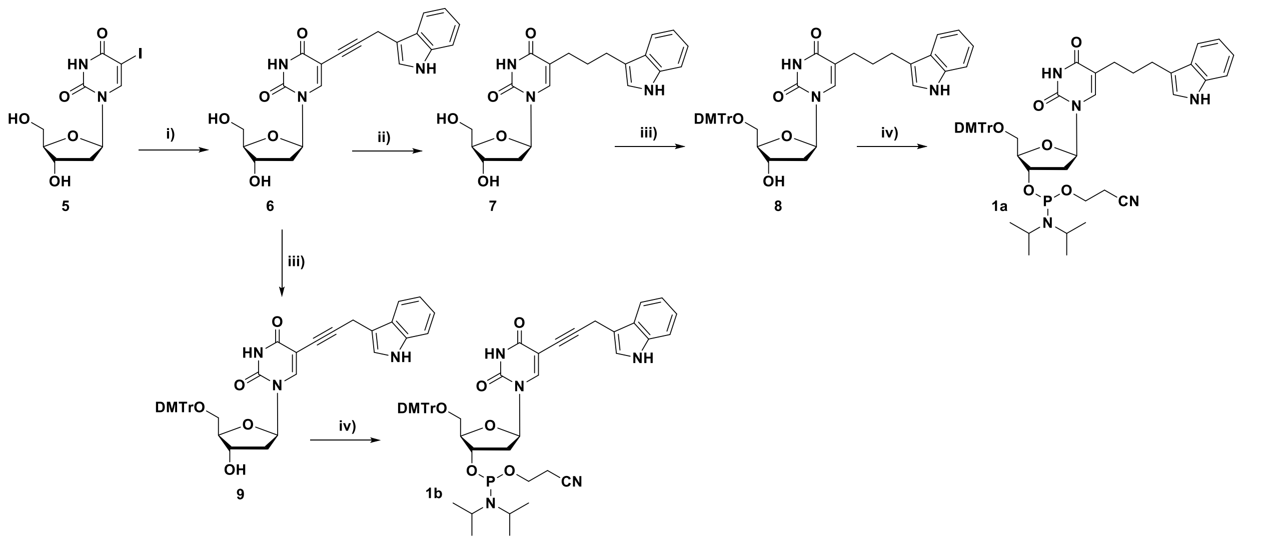

**Scheme 1**: **Reagents and conditions:** (i) $Pd(PPh_3)_4$, CuI, $Et_3N$, 1*H*-Indole,3-(2-propynyl), DMF, 60 °C 2 h (ii) $H_2$, 10% Pd/C, MeOH, 50 °C, overnight (iii) DMTrCl, DMAP, pyridine, rt, overnight (iv) 2-cyanoethyl-*N*,*N*-diisopropylchlorophosphoramidite, DIPEA, DCM, 0 °C to rt, 1.0 h.

**Synthesis of 5-(3-(1*H*-indol-3-yl)prop-1-yn-1-yl)-1-((2R,4S,5R)-4-hydroxy-5 (hydroxymethyl)tetrahydrofuran-2-yl)pyrimidine-2,4(1*H*,3*H*)-dione (6):**

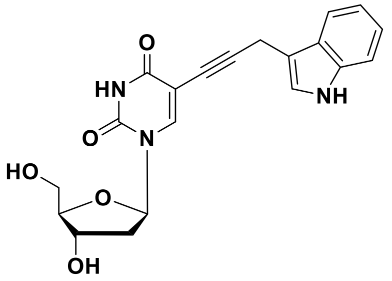

To a solution of 5-iododeoxyuridine (**5**) (3.0 g, 8.50 mmol) in anhydrous DMF (30 mL), CuI (0.48 g, 2.50 mmol) and TEA (5.90 mL, 42.0 mmol) were added under stirring. After 5 minutes of stirring, 3-(prop-2-yn-1-yl)-1*H*-indole[1] (1.7 g, 11.0 mmol) and $Pd(PPh_3)_4$ (0.98 g, 0.85 mmol) were added to the reaction mixture. The reaction mixture was then heated to 60 °C and stirred for 2 h. The progress of the reaction was followed using TLC. After completion of the reaction, the reaction mixture was diluted with ethyl acetate (50 mL) and filtered through a celite. The filtrate was evaporated under reduced pressure and the crude product was purified by flash chromatography (0–10% MeOH in $CH_2Cl_2$). The desired compound **6** (2.3 g, 6.0 mmol, 71%) was obtained as a white solid.

**R$_f$** (DCM/Methanol 9:1) 0.45;

**$^1$H NMR** (500 MHz, DMSO-*d$_6$*) $\delta_H$ 11.58 (s, 1H), 10.90 (s, 1H), 8.16 (s, 1H), 7.60 (d, *J* = 7.8 Hz, 1H), 7.36 (d, *J* = 8.1 Hz, 1H), 7.29 (d, *J* = 1.1 Hz, 1H), 7.09 (dd, *J* = 8.0, 7.1 Hz, 1H), 7.00 (t, *J* = 7.4 Hz, 1H), 6.12 (t, *J* = 6.7 Hz, 1H), 5.25 (d, *J* = 4.3 Hz, 1H), 5.11 (t, *J* = 5.0 Hz, 1H), 4.26 – 4.21 (m, 1H), 3.85 (s, 2H), 3.80 (d, *J* = 3.0 Hz, 1H), 3.66 – 3.53 (m, 2H), 2.18 – 2.06 (m, 2H);

**$^{13}$C NMR** (126 MHz, DMSO-*d$_6$*) $\delta_C$ 162.3, 149.9, 143.7, 136.8, 126.9, 123.5, 121.6, 118.9, 118.9 111.9, 109.8, 99.5, 92.1, 88.1, 85.1, 73.6, 70.7, 61.5, 16.2;

**HRMS** ESI/Q-TOF $[M+Na]^+$ calcd. for $C_{20}H_{19}N_3NaO_5$ 404.1217, found 404.1220.

**Synthesis of 5-(3-(1*H*-indol-3-yl)propyl)-1-((2R,4S,5R)-4-hydroxy-5-(hydroxymethyl)tetrahydrofuran-2-yl)pyrimidine-2,4(1*H*,3*H*)-dione (7):**

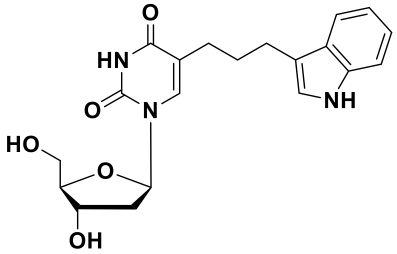

To a solution of compound **6** (2.0 g, 5.2 mmol) in methanol (40 mL), 10% Pd/C (0.56 g, 0.52 mmol) was added under stirring. The reaction mixture was then stirred under hydrogen atmosphere at 50 °C overnight. After completion of the reaction, the reaction mixture was filtered through celite to remove the catalyst. The filtrate was evaporated under reduced pressure to yield the pure compound **7** (1.5 g, 3.9 mmol, 74 %) as a white solid.

**R$_f$** (DCM/Methanol 9:1) 0.4;

**$^1$H NMR** (500 MHz, DMSO-*d$_6$*) $\delta_H$ 11.26 (s, 1H), 10.73 (s, 1H), 7.72 (s, 1H), 7.48 (d, *J* = 7.8 Hz, 1H), 7.32 (d, *J* = 8.1 Hz, 1H), 7.12 (s, 1H), 7.05 (t, *J* = 7.4 Hz, 1H), 6.96 (t, *J* = 7.3 Hz, 1H), 6.18 (t, *J* = 6.8 Hz, 1H), 5.23 (d, *J* = 3.8 Hz, 1H), 5.04 (t, *J* = 4.8 Hz, 1H), 4.25 (s, 1H), 3.78 (d, *J* = 2.6 Hz, 1H), 3.63 – 3.51 (m, 2H), 2.68 (t, *J* = 7.2 Hz, 2H), 2.34 – 2.24 (m, 2H), 2.16 – 2.03 (m, 2H), 1.88 – 1.76 (m, 2H);

**$^{13}$C NMR** (126 MHz, DMSO-*d$_6$*) $\delta_C$ 163.9, 150.8, 136.7, 136.6, 127.6, 122.6, 121.2, 118.7, 118.5, 114.6, 114.0, 111.8, 87.8, 84.4, 70.9, 61.8, 46.1, 28.9, 26.8, 24.8;

**HRMS** (ESI/Q-TOF) $[M+Na]^+$ calcd. for $C_{20}H_{23}N_3NaO_5$ 408.1530, found 408.1544.

**Synthesis of 5-(3-(1*H*-indol-3-yl)propyl)-1-((2R,4S,5R)-5-((bis(4-methoxyphenyl)(phenyl)methoxy)methyl)-4-hydroxytetrahydrofuran-2-yl)pyrimidine-2,4(1*H*,3*H*)-dione (8):**

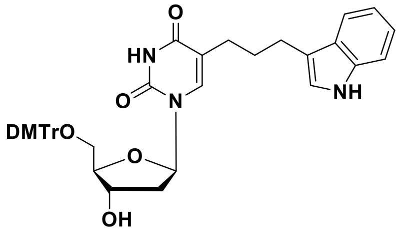

To a solution of compound **7** (1.0 g, 2.6 mmol) in anhydrous pyridine (15 mL), DMAP (32 mg, 0.26 mmol) was added. To this mixture, a solution of 4,4′-dimethoxytrityl chloride (1.1 g, 3.1 mmol) in 5 mL of anhydrous pyridine was added in four equal portions over the time of 1 h. The reaction mixture was stirred at room temperature for 12 h. After completion of the reaction, the solvent was evaporated under reduced pressure. The resulting crude mixture was purified by flash chromatography (0–2% MeOH in $CH_2Cl_2$ with 0.2% of $Et_3N$) yielding the desired compound **8** (0.90 g, 1.31 mmol, 50%) as a yellow foam.

**R$_f$** (DCM/Methanol 9:1) 0.5;

**$^1$H NMR** (500 MHz, DMSO-*d$_6$*) $\delta_H$ 11.32 (s, 1H), 10.71 (s, 1H), 7.44 (s, 1H), 7.39 (d, *J* = 7.9 Hz, 2H), 7.34 (d, *J* = 7.9 Hz, 1H), 7.30 (m 3H), 7.25 (m 4H), 7.21 (t, *J* = 7.3 Hz, 1H), 7.04 (t, *J* = 7.6 Hz, 1H), 6.98 (s, 1H), 6.91 (dd, *J* = 17.6, 9.9 Hz, 1H), 6.85 (m, 4H), 6.21 (t, *J* = 6.8 Hz, 1H), 5.34 (d, *J* = 4.5 Hz, 1H), 4.32 (dd, *J* = 7.0, 3.1 Hz, 1H), 3.88 (t, *J* = 6.8 Hz, 1H), 3.68 (d, *J* = 3.0 Hz, 7H), 3.18 (qd, *J* = 10.4, 3.6 Hz, 2H), 2.44 (t, *J* = 7.6 Hz, 2H), 2.26 (dt, *J* = 13.5, 6.9 Hz, 1H), 2.21 – 2.12 (m, 1H), 2.07 – 1.97 (m, 1H), 1.90 (dt, *J* = 14.1, 7.5 Hz, 1H), 1.68 – 1.53 (m, 2H);

**$^{13}$C NMR** (126 MHz, DMSO-$d_6$) $\delta_C$ 163.7, 158.6, 150.7, 150.0, 145.1, 136.7, 136.1, 135.9, 135.7, 130.2, 130.1, 128.3, 128.6, 127.6, 127.3, 122.4, 121.2, 118.6, 118.5, 114.6, 114.5, 113.7, 111.7, 86.3, 85.9, 84.3, 71.1, 64.2, 55.4, 49.1, 29.5, 27.1, 24.9;

**HRMS** (ESI/Q-TOF) $[M+Na]^+$ calcd. for $C_{41}H_{41}N_3NaO_7$ 710.2837, found 710.2840.

**Synthesis of (2R,3S,5R)-5-(5-(3-(1*H*-indol-3-yl)propyl)-2,4-dioxo-3,4-dihydropyrimidin-1(2*H*)-yl)-2-((bis(4-methoxyphenyl)(phenyl)methoxy)methyl)tetrahydrofuran-3-yl (2-cyanoethyl) diisopropylphosphoramidite (1a):**

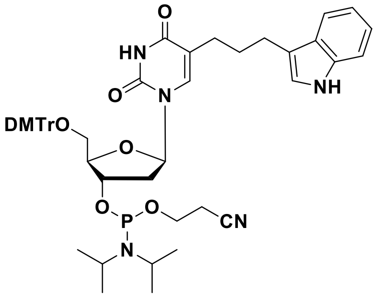

To a solution of compound **8** (0.6 g, 0.87 mmol) in anhydrous $CH_2Cl_2$, *N*,*N*-diisopropylethylamine (0.75 mL, 4.36 mmol) was added under stirring. The reaction mixture was cooled to 0 °C, and 2-cyanoethyl-*N*,*N*-diisopropylchlorophosphoramidite (0.3 mL, 1.40 mmol) was added under an argon atmosphere. The mixture was allowed to warm to room temperature and stirred for 1.0 h. The reaction mixture was then diluted with anhydrous $CH_2Cl_2$ (30 mL) and washed successively with 5% $NaHCO_3$ (30 mL) and brine solution (30 mL). The organic phase was dried over anhydrous $Na_2SO_4$ and concentrated under reduced pressure. The crude product was purified by column chromatography using (60% EtOAc in hexane containing 0.5% $Et_3N$) as the eluent. The desired compound **1a** (0.52 g, 0.6 mmol, 68%) was obtained as a white foam.

**$R_f$** (EtOAc/Hexane 7:3) 0.6;

**$^1$H NMR** (500 MHz, $CD_3CN$) $\delta_H$ 8.53 (s, 1H), 7.85 (s, 1H), 7.52 (m, 1H), 7.44 (m, 3H), 7.31 (m, 7H), 7.24 (m, 1H), 7.15 (t, *J* = 7.6 Hz, 1H), 7.05 (t, *J* = 7.4 Hz, 1H), 6.82 (dd, *J* = 8.0, 5.8 Hz, 4H), 6.75 (d, *J* = 3.0 Hz, 1H), 6.40 (q, *J* = 7.4 Hz, 1H), 4.70 – 4.59 (m, 1H), 4.23 – 4.11 (m, 3H), 3.90 (dd, *J* = 12.0, 5.8 Hz, 1H), 3.72 (d, *J* = 6.1 Hz, 6H), 3.67 – 3.46 (m, 4H), 3.31 (td, *J* = 10.1, 2.5 Hz, 1H), 2.62 (q, *J* = 6.4 Hz, 1H), 2.57 – 2.47 (m, 2H), 2.43 (dd, *J* = 15.1, 8.8 Hz, 1H), 2.28 (dt, *J* = 19.2, 7.0 Hz, 1H), 2.18 – 2.10 (m, 1H), 1.98 – 1.86 (m, 1H), 1.76 (ddd, *J* = 9.6, 8.9, 4.0 Hz, 1H), 1.2-1.7 (m, 9H), 1.07 (d, *J* = 10 Hz, 3H);

**$^{13}$C NMR** (126 MHz, $CD_3CN$) $\delta_C$ 163.4, 158.8, 150.4, 144.9, 136.5, 135.8, 135.8, 135.7, 135.6, 135.6, 130.2, 130.2, 130.1, 130.1, 128.2, 128.1, 128.0, 127.5, 127.1, 121.8, 121.3, 118.6, 118.5, 115.2, 114.9, 113.2, 111.2, 86.4, 84.8, 84.4, 73.4, 63.3, 60.0, 58.6, 58.4, 57.2, 54.9, 43.1, 43.0, 39.1, 29.1, 26.8, 24.5, 24.0, 23.9, 21.0, 20.2, 20.1, 20.0, 13.6;

**$^{31}$P NMR** (202 MHz, $CD_3CN$) $\delta_P$ 149.3, 148.9;

**HRMS** (ESI/Q-TOF) $[M+Na]^+$ calcd. for $C_{50}H_{58}N_5NaO_8P$ 910.3915, found 910.3795.

**Synthesis of 5-(3-(1*H*-indol-3-yl)prop-1-yn-1-yl)-1-((2R,4S,5R)-5-((bis(4-methoxyphenyl)(phenyl)methoxy)methyl)-4-hydroxytetrahydrofuran-2-yl)pyrimidine-2,4(1*H*,3*H*)-dione (9):**

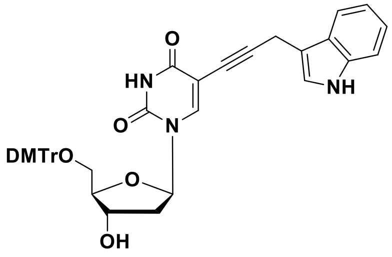


To a solution of compound **6** (1.0 g, 2.6 mmol) in anhydrous pyridine (15 mL), DMAP (32 mg, 0.26 mmol) was added under stirring. To this mixture, a solution of 4,4'-dimethoxytrityl chloride (1.1 g, 3.1 mmol) in 5 mL of anhydrous pyridine was added in four equal portions over the time of 1 h. The reaction mixture was stirred at room temperature for 12 h. After completion of the reaction, the solvent was evaporated under reduced pressure. The resulting crude mixture was purified by flash chromatography (0–2% MeOH in $CH_2Cl_2$, with 0.2% of $Et_3N$) yielding the desired compound **9** (0.95 g, 1.39 mmol, 53%) as a yellowish foam.

**$R_f$** (DCM/Methanol 9:1) 0.6;

**$^1$H NMR** (500 MHz, DMSO-*$d_6$*) $\delta_H$ 11.64 (s, 1H), 10.88 (s, 1H), 7.90 (s, 1H), 7.53 (d, *J* = 7.9 Hz, 1H), 7.42 (d, *J* = 7.7 Hz, 2H), 7.36 (d, *J* = 8.1 Hz, 1H), 7.32 – 7.28 (m, 7H), 7.20 (d, *J* = 4.7 Hz, 2H), 7.09 (t, *J* = 7.5 Hz, 1H), 6.96 (t, *J* = 7.4 Hz, 1H), 6.86 (dd, *J* = 8.7, 4.0 Hz, 5H), 6.13 (t, *J* = 6.6 Hz, 1H), 5.35 (d, *J* = 4.4 Hz, 1H), 4.29 (s, 1H), 3.93 (s, 6H), 3.26 (dd, *J* = 10.4, 5.3 Hz, 1H), 3.13 – 3.06 (m, 1H), 2.28 (dd, *J* = 13.4, 6.7 Hz, 1H), 2.23 – 2.13 (m, 1H);

**$^{13}$C NMR** (126 MHz, DMSO-*$d_6$*) $\delta_C$ 162.3, 158.5, 149.8, 145.3, 142.6, 136.9,136.1, 135.8, 130.2, 130.1, 128.4, 127.1, 123.4, 121.6, 118.9, 118.8, 113.7, 113.6, 112.0, 109.6, 99.7, 92.2, 86.3, 86.2, 85.4, 73.1, 71.0, 64.2, 55.5, 16.1;

**HRMS** (ESI/Q-TOF) $[M+Na]^+$ calcd. for $C_{41}H_{37}N_3NaO_7$ 706.2529, found 707.2556.

**Synthesis of (2R,3S,5R)-5-(5-(3-(1*H*-indol-3-yl)prop-1-yn-1-yl)-2,4-dioxo-3,4-dihydropyrimidin-1(2*H*)-yl)-2-((bis(4-methoxyphenyl)(phenyl)methoxy)methyl)tetrahydrofuran-3-yl (2-cyanoethyl) diisopropylphosphoramidite (1b):**

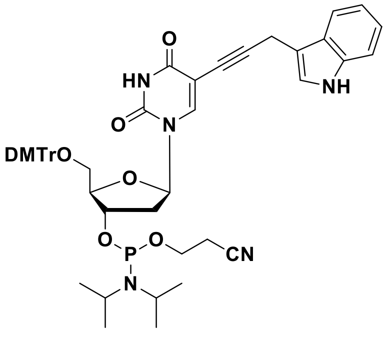


To a solution of compound **9** (0.5 g, 0.73 mmol) in anhydrous $CH_2Cl_2$, *N*,*N*-diisopropylethylamine (0.63 mL, 3.66 mmol) was added under stirring. The reaction mixture was cooled to 0 °C, and 2-cyanoethyl-*N*,*N*-diisopropylchlorophosphoramidite (0.25 mL, 1.2 mmol) was added under an argon atmosphere. The mixture was allowed to warm to room temperature and stirred for 1.0 h. The reaction mixture was then diluted with anhydrous $CH_2Cl_2$ (30 mL) and washed successively with 5% $NaHCO_3$ (30 mL) and brine solution (30 mL). The organic phase was dried over anhydrous $Na_2SO_4$ and concentrated under reduced pressure. The crude product was purified by column chromatography using (50% EtOAc in hexane containing 0.5% $Et_3N$) as the eluent. The desired compound **1b** (0.37 g, 0.43 mmol, 58%) was obtained as a white foam.

**$R_f$** (EtOAc/Hexane 7:3) 0.6;

$^{1}$**H NMR** (500 MHz, $CD_3CN$) $\delta_H$ 9.16 (s, 1H), 9.09 (s, 1H), 7.95 (d, *J* = 13.5 Hz, 1H), 7.56 (d, *J* = 7.9 Hz, 1H), 7.54 – 7.50 (m, 1H), 7.46 – 7.38 (m, 5H), 7.34 (td, *J* = 7.8, 2.5 Hz, 2H), 7.28 – 7.22 (m, 1H), 7.21 – 7.15 (m, 1H), 7.13 – 7.09 (m, 1H), 7.09 – 7.04 (m, 1H), 6.87 (m, 4H), 6.20 (m, 1H), 4.68 (m, 1H), 4.24 – 4.13 (m, 1H), 3.82 (m, 1H), 3.75 (s, 3H), 3.74 (s, 3H), 3.72 (dd, *J* = 7.9, 5.7 Hz, 1H), 3.68 – 3.58 (m, 4H), 3.40 – 3.27 (m, 2H), 2.69 (t, *J* = 6.0 Hz, 1H), 2.58 (td, *J* = 6.0, 2.3 Hz, 3H), 2.48 – 2.37 (m, 1H), 1.23-1.20 (m, 9H), 1.12 (d, J = 5 Hz, 3H);

$^{13}$**C NMR** (126 MHz, $CD_3CN$) $\delta_C$ 161.9, 158.8, 149.4, 145.0, 142.0, 136.7, 135.9, 135.5, 130.2, 130.1, 128.0, 128.0, 126.9, 126.7, 122.8, 121.7, 119.0, 118.4, 113.2, 113.2, 111.4, 110.1, 100.0, 91.9, 86.7, 85.4, 85.4, 73.3, 73.0, 72.4, 63.0, 58.6, 58.5, 58.2, 55.0, 45.1, 45.0, 43.1, 43.0, 39.8, 39.7, 24.0, 23.9, 23.9, 22.2, 22.2, 20.1, 20.0, 19.7, 15.6;

$^{31}$**P NMR** (202 MHz, $CD_3CN$) $\delta_P$ 148.0;

**HRMS** (ESI/Q-TOF) $[M+H]^+$ calcd. for $C_{50}H_{55}N_5O_8P$ 884.3783, found 884.3750.

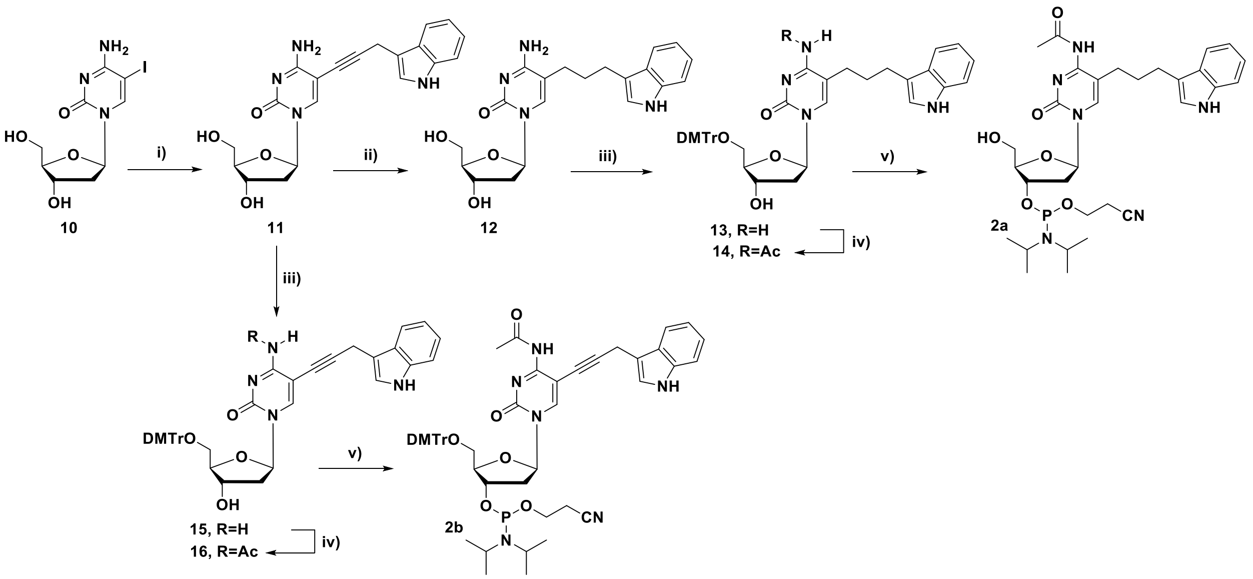


**Scheme 2**: **Reagents and conditions:** (i) $Pd(PPh_3)_4$, CuI, $Et_3N$, 1*H*-Indole,3-(2-propynyl), DMF, 60 °C 1.5 h (ii) $H_2$, 10% Pd/C, MeOH, 50 °C, overnight (iii) DMTrCl, DMAP, pyridine, rt, overnight (iv) $Ac_2O$, DMF, rt, 8-10 h (v) 2-cyanoethyl-*N*,*N*-diisopropylchlorophosphoramidite, DIPEA, DCM, 0 °C to rt, 1.5 h.

**Synthesis of 5-(3-(1*H*-indol-3-yl)prop-1-yn-1-yl)-4-amino-1-((2R,4S,5R)-4-hydroxy-5-(hydroxymethyl)tetrahydrofuran-2-yl)pyrimidin-2(1*H*)-one (11):**

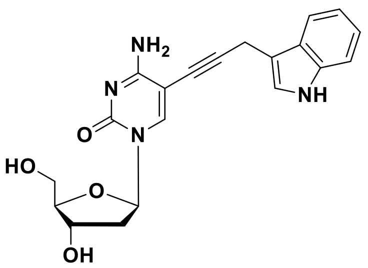


To a solution of 5-iodo-2'-deoxycytidine (**10**) (2.0 g, 5.66 mmol) in anhydrous DMF (30 mL), CuI (0.32 g, 1.70 mmol) and TEA (3.95 mL, 28.3 mmol) were added under stirring. After 5 minutes of stirring, 3-(prop-2-yn-1-yl)-1*H*-indole (1.14 g, 7.36 mmol) and $Pd(PPh_3)_4$ (0.66 g, 0.57 mmol) were added to the reaction mixture. The reaction mixture was heated to 60 °C and stirred for 1.5 h. After completion of the reaction, the reaction mixture was diluted with ethyl acetate (50 mL) and filtered through celite. The filtrate was evaporated under reduced pressure and the crude product was purified by flash chromatography using dichloromethane and methanol (8:2, v/v) as eluent. The desired compound **11** (1.5 g, 3.9 mmol, 70%) was obtained as a white solid.

**$R_f$** (DCM/Methanol 9:1) 0.4;

**$^1$H NMR** (500 MHz, DMSO-$d_6$) $\delta_H$ 10.09 (s, 1H), 7.29 (s, 1H), 6.90 (br.s, 2H), 6.83 (d, *J* = 7.8 Hz, 1H), 6.55 (d, *J* = 8.1 Hz, 1H), 6.46 (s, 1H), 6.28 (t, *J* = 7.4 Hz, 1H), 6.19 (t, *J* = 7.4 Hz, 1H), 5.97 (s, 1H), 5.31 (t, *J* = 6.5 Hz, 1H), 4.38 (d, *J* = 4.0 Hz, 1H), 4.23 (t, *J* = 4.9 Hz, 1H), 3.09 (s, 2H), 2.97 (d, *J* = 2.9 Hz, 1H), 2.84 – 2.69 (m, 2H), 1.69 (s, 1H), 1.36 – 1.27 (m, 1H), 1.17 (dt, *J* = 13.1, 6.5 Hz, 1H);

**$^{13}$C NMR** (126 MHz, DMSO-$d_6$) $\delta_C$ 167.1, 156.2, 146.3, 139.0, 129.1, 125.7, 123.9, 121.1, 121.1 114.2, 112.0, 97.0, 93.0, 90.1, 87.9, 74.8, 72.8, 63.7, 43.4, 18.5;

**HRMS** (ESI/Q-TOF) $[M+H]^+$ calcd. for $C_{20}H_{21}N_4O_4$ 381.1563, found 381.1542.

**Synthesis of 5-(3-(1*H*-indol-3-yl)propyl)-4-amino-1-((2R,4S,5R)-4-hydroxy-5 (hydroxymethyl)tetrahydrofuran-2-yl)pyrimidin-2(1*H*)-one (12):**

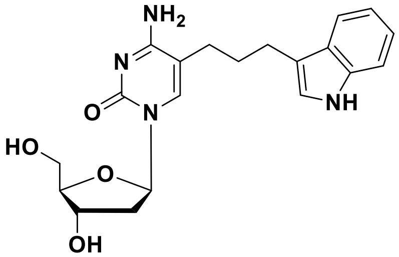


To a solution of compound **11** (1.3 g, 3.4 mmol) in methanol (40 mL), 10% Pd/C (0.36 g, 0.34 mmol) was added under stirring. The reaction mixture was then stirred under a hydrogen atmosphere at 50 °C overnight. The progress of the reaction was followed using TLC. After completion of the reaction, the reaction mixture was filtered through celite to remove the catalyst. The filtrate was evaporated under reduced pressure to yield the pure compound **12** (1.1 g, 2.9 mmol, 84 %) as a white solid

**$R_f$** (DCM/Methanol 8:2) 0.5;

**$^1$H NMR** (500 MHz, DMSO-$d_6$) $\delta_H$ 10.74 (s, 1H), 7.68 (s, 1H), 7.50 (d, *J* = 7.8 Hz, 1H), 7.32 (d, *J* = 8.1 Hz, 1H), 7.29 (s, 1H), 7.11 (d, *J* = 1.7 Hz, 1H), 7.05 (t, *J* = 7.5 Hz, 1H), 6.96 (t, *J* = 7.4 Hz, 1H), 6.93 (s, 1H), 6.18 (t, *J* = 6.7 Hz, 1H), 5.18 (d, *J* = 4.2 Hz, 1H), 5.01 (t, *J* = 5.2 Hz, 1H), 4.22 (td, *J* = 6.7, 3.4 Hz, 1H), 3.77 (q, *J* = 3.5 Hz, 1H), 3.64 – 3.50 (m, 2H), 2.72 (t, *J* = 5.0 Hz, 2H), 2.41 – 2.30 (m, 2H), 2.09 (ddd, *J* = 13.0, 5.9, 3.3 Hz, 1H), 1.99 (dt, *J* = 13.2, 6.6 Hz, 1H), 1.85 – 1.76 (m, 2H);

**$^{13}$C NMR** (126 MHz, DMSO-*d$_6$*) $\delta_C$ 156.0, 152.5, 151.7, 150.2, 136.8, 127.6, 122.9, 121.3, 118.8, 118.8, 114.3, 111.8, 88.7, 85.0, 71.8, 62.8, 46.1, 38.4, 31.2, 28.8, 27.7, 24.8;

**HRMS** (ESI/Q-TOF) $[M+H]^+$ calcd. for $C_{20}H_{25}N_4O_4$ 385.1876, found 385.1864.

**Synthesis of 5-(3-(1*H*-indol-3-yl)propyl)-4-amino-1-((2R,4S,5R)-5-((bis(4 methoxyphenyl)(phenyl)methoxy)methyl)-4-hydroxytetrahydrofuran-2-yl)pyrimidin-2(1*H*)-one (13):**

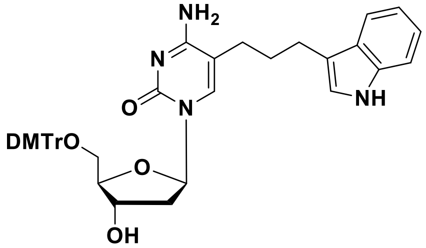


To a solution of compound **12** (1.0 g, 2.6 mmol) in anhydrous pyridine (15 mL), DMAP (32 mg, 0.26 mmol) was added. To this mixture, a solution of 4,4'-dimethoxytrityl chloride (1.1 g, 3.1 mmol) in 5 mL of anhydrous pyridine was added in four equal portions over the time of 1 h. The reaction mixture was stirred at room temperature for 12 h. After completion of the reaction, the solvent was evaporated under reduced pressure. The resulting crude mixture was purified by flash chromatography using (0–2% MeOH in $CH_2Cl_2$, with 0.2% of $Et_3N$) yielding the desired compound **13** (0.97 g, 1.41 mmol, 54%) as a yellowish foam.

**R$_f$** (DCM/Methanol 9:1) 0.5;

**$^1$H NMR** (500 MHz, DMSO-*d$_6$*) $\delta_H$ 10.70 (s, 1H), 7.42 (s, 1H), 7.39 (d, *J* = 7.5 Hz, 3H), 7.33 – 7.22 (m, 9H), 7.19 (t, *J* = 7.1 Hz, 1H), 7.04 (t, *J* = 7.4 Hz, 1H), 6.95 (s, 1H), 6.92 (t, *J* = 7.5 Hz, 1H), 6.83 (d, *J* = 7.0 Hz, 4H), 6.24 (t, *J* = 6.6 Hz, 1H), 5.27 (d, *J* = 4.3 Hz, 1H), 4.28 (s, 1H), 3.88 (d, *J* = 3.4 Hz, 1H), 3.67 (s, 3H), 3.66 (s 3H), 3.17 (s, 2H), 2.47 – 2.39 (m, 2H), 2.20 – 2.05 (m, 3H), 2.02 – 1.92 (m, 1H), 1.70 – 1.49 (m, 2H);

**$^{13}$C NMR** (126 MHz, DMSO-*d$_6$*) $\delta_C$ 165.2, 158.6, 158.6, 155.3, 155.3, 145.2, 138.0, 136.7, 135.97, 135.7, 130.2, 130.2, 128.3, 128.1, 127.5, 127.2, 122.3, 121.2, 118.7, 118.5, 114.8, 113.7, 111.7, 106.4, 86.2, 85.7, 84.9, 71.1, 64.2, 55.4, 40.9, 29.7, 27.5, 24.8;

**HRMS** (ESI/Q-TOF) $[M+H]^+$ calcd. for $C_{41}H_{43}N_4O_6$ 687.3183, found 687.3166.

**Synthesis of N-(5-(3-(1*H*-indol-3-yl)propyl)-1-((2R,4S,5R)-5-((bis(4-methoxyphenyl)(phenyl)methoxy)methyl)-4-hydroxytetrahydrofuran-2-yl)-2-oxo-1,2-dihydropyrimidin-4-yl)acetamide (14):**

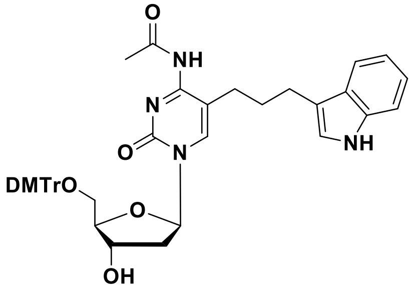


To a solution of compound **13** (0.85g, 1.24 mmol) in dry DMF (12 mL), acetic anhydride (0.13 mL, 1.36 mmol) was added dropwise over 15 minutes at 0 °C under an argon atmosphere. The reaction mixture was then stirred at room temperature for 8 h. Progress of the reaction was monitored by TLC. After completion of the reaction, the reaction mixture was concentrated under reduced pressure. The crude product was purified by flash chromatography using (0–2% MeOH in $CH_2Cl_2$, with 0.2% of $Et_3N$) yielding the desired compound **14** (0.55 g, 0.76 mmol, 61%) as a pale-yellow solid.

**$R_f$** (DCM/Methanol 9:1) 0.55;

**$^1$H NMR** (500 MHz, DMSO-*d$_6$*) $\delta_H$ 10.72 (s, 1H), 9.84 (s, 1H), 7.84 (s, 1H), 7.37 (t, *J* = 9.2 Hz, 3H), 7.30 (dd, *J* = 14.2, 7.9 Hz, 3H), 7.22 (m, 4H), 7.20 (d, *J* = 7.3 Hz, 1H), 7.04 (t, *J* = 7.4 Hz, 1H), 6.96 – 6.93 (m, 1H), 6.91 (d, *J* = 7.3 Hz, 1H), 6.84 (d, *J* = 8.5 Hz, 4H), 6.15 (t, *J* = 6.2 Hz, 1H), 5.34 (d, *J* = 4.6 Hz, 1H), 4.30 – 4.24 (m, 1H), 3.97 (dd, *J* = 7.6, 3.9 Hz, 1H), 3.68 (s, 3H), 3.66 (s, 3H) 3.26 – 3.17 (m, 2H), 2.45 – 2.41 (m, 2H), 2.36 – 2.30 (m, 2H), 2.18 (s, 3H), 2.16 – 2.08 (m, 2H), 1.64 – 1.51 (m, 2H);

**$^{13}$C NMR** (126 MHz, DMSO-*d$_6$*) $\delta_C$ 171.1, 162.2, 158.6, 154.3, 145.1, 142.1, 136.7, 135.9, 135.7, 130.2, 130.2, 128.4, 128.1, 127.5, 127.3, 122.4, 121.3, 118.6, 118.5, 118.4, 114.5, 113.7, 113.4, 111.7, 110.5, 86.4, 86.3, 70.6, 55.4, 41.2, 29.8, 27.6, 25.2, 24.8;

**HRMS** (ESI/Q-TOF) $[M+H]^+$ calcd. for $C_{43}H_{45}N_4O_7$ 729.3288, found 729.3376.

**Synthesis of (2R,3S,5R)-5-(5-(3-(1*H*-indol-3-yl)propyl)-4-acetamido-2-oxopyrimidin-1(2*H*)-yl)-2-(hydroxymethyl)tetrahydrofuran-3-yl (2-cyanoethyl) diisopropylphosphoramidite (2a):**

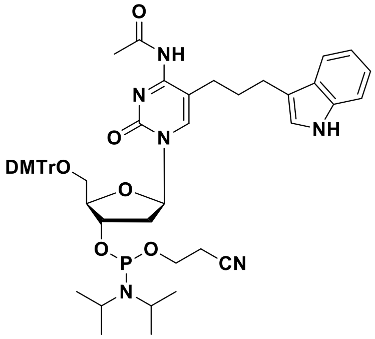


To a solution of compound **14** (0.4 g, 0.55 mmol) in anhydrous $CH_2Cl_2$, *N*,*N*-diisopropylethylamine (0.47 mL, 2.74 mmol) was added under stirring. The reaction mixture was cooled to 0 °C, and 2-cyanoethyl-*N*,*N*-diisopropylchlorophosphoramidite (0.19 mL, 0.88 mmol) was added under an argon atmosphere. The mixture was allowed to warm to room temperature and stirred for 1.5 h. The reaction mixture was then diluted with anhydrous $CH_2Cl_2$ (20 mL) and washed successively with 5% $NaHCO_3$ (20 mL) and brine solution (20 mL). The organic phase was dried over anhydrous $Na_2SO_4$ and concentrated under reduced pressure. The crude product was purified by column chromatography using (70% EtOAc in hexane containing 0.5% $Et_3N$) as the eluent. The desired compound **2a** (0.28 g, 0.31 mmol, 56%) was obtained as a white foam.

**$R_f$** (EtOAc/Hexane 8:2) 0.5;

**$^1$H NMR** (500 MHz, $CD_3CN$) $\delta$ 9.02 (s, 1H), 7.49 (d, *J* = 7.6 Hz, 3H), 7.46 (br.s, 1H), 7.42 – 7.31 (m, 8H), 7.27 (m, 1H), 7.14 (t, *J* = 7.5 Hz, 1H), 7.02 (t, *J* = 7.4 Hz, 1H), 6.87 (m, 5H), 6.22 (br. s, 1H), 4.65 (br.s, 1H), 4.20 (br. s, 1H), 3.76 (s, 3H), 3.75 (s, 3H), 3.72 – 3.58 (m, 4H), 3.44 (m, 1H), 3.35 (dd, *J* = 10.8, 3.9 Hz, 1H), 2.56 (m, 5H), 1.21 (s, 3H), 1.20 (s, 3H), 1.19 (s, 3H), 1.18 (s, 3H);

**$^{13}$C NMR** (126 MHz, $CD_3CN$) $\delta$ 173.4, 161.5, 147.6, 139.3, 138.5, 138.3, 132.9, 132.9, 130.9, 130.7, 129.8, 124.6, 124.1, 121.3, 121.2, 121.1, 115.9, 114.0, 89.2, 88.1, 75.2, 75.2, 65.6, 62.7, 61.3, 61.2, 59.9, 57.7, 45.8, 45.7, 29.8, 27.0, 26.7, 26.6, 23.7, 22.9, 22.7,22.7, 16.3;

**$^{31}$P NMR** (202 MHz, $CD_3CN$) $\delta_P$ 148.1, 148.1;

**HRMS** (ESI/Q-TOF) $[M+H]^+$ calcd. for $C_{52}H_{62}N_6O_8P$ 929.4361, found 929.4310.

**Synthesis of 5-(3-(1*H*-indol-3-yl)prop-1-yn-1-yl)-4-amino-1-((2R,4S,5R)-5-((bis(4-methoxyphenyl)(phenyl)methoxy)methyl)-4-hydroxytetrahydrofuran-2-yl)pyrimidin-2(1*H*)-one (15):**

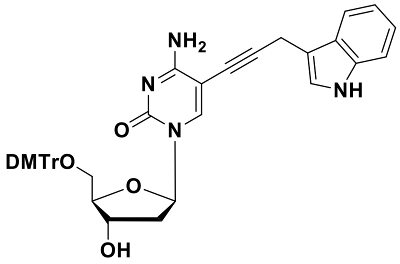


To a solution of compound **11** (1.1 g, 2.9 mmol) in anhydrous pyridine (10 mL), DMAP (35 mg, 0.29 mmol) was added under stirring. To this mixture, a solution of 4,4'-dimethoxytrityl chloride (1.2 g, 3.5 mmol) in 5 mL of anhydrous pyridine was added in four equal portions over the time of 1 h. The reaction mixture was stirred at room temperature for 12 h. After completion of the reaction, the solvent was evaporated under reduced pressure. The resulting crude mixture was purified by flash chromatography using (0–2% MeOH in $CH_2Cl_2$, with 0.2% of $Et_3N$) as the eluent, yielding the desired compound **15** (1.1 g, 1.6 mmol, 56%) as a white solid.

**$R_f$** DCM/Methanol 9:1) 0.5;

**$^1$H NMR** (500 MHz, DMSO-$d_6$) $\delta_H$ 10.87 (d, *J* = 62.5 Hz, 1H), 7.86 (s, 1H), 7.75 (s, 1H), 7.57 (d, *J* = 7.9 Hz, 1H), 7.40 (d, *J* = 8.2 Hz, 1H), 7.36 (d, *J* = 8.1 Hz, 1H), 7.27 (m 6H), 7.20 – 7.12 (m, 2H), 7.08 (t, *J* = 7.6 Hz, 1H), 6.96 (t, *J* = 7.4 Hz, 1H), 6.83 (dd, *J* = 8.6, 6.0 Hz, 4H), 6.77 (br.s, 1H), 6.12 (t, *J* = 6.6 Hz, 1H), 5.29 (d, *J* = 4.4 Hz, 1H), 4.21 (m, 1H), 3.93 (d, *J* = 2.0 Hz, 1H), 3.75 (d, *J* = 9.5 Hz, 2H), 3.68 (s, 3H), 3.67 (s, 3H), 3.23 (dd, *J* = 10.4, 5.2 Hz, 1H), 3.08 (dd, *J* = 10.4, 2.4 Hz, 1H), 2.22 (ddd, *J* = 12.8, 5.8, 3.3 Hz, 1H), 2.11 (dt, *J* = 13.4, 6.7 Hz, 1H);

**$^{13}$C NMR** (126 MHz, DMSO-$d_6$) $\delta_C$ 165.0, 158.5, 158.5, 153.9, 145.2, 143.4, 136.8, 136.1, 135.7, 130.3, 130.1, 128.3, 128.0, 127.1, 126.9, 123.4, 121.6, 118.9, 118.7, 113.7, 113.6, 111.9, 109.7, 94.7, 91.1, 86.3, 86.2, 85.9, 72.2, 71.1, 64.2, 55.4, 41.2, 16.3;

**HRMS** (ESI/Q-TOF) $[M+H]^+$ calcd. for $C_{41}H_{39}N_4O_6$ 682.2791, found 682.2862.

**Synthesis of N-(5-(3-(1*H*-indol-3-yl)prop-1-yn-1-yl)-1-((2R,4S,5R)-5-((bis(4-methoxyphenyl)(phenyl)methoxy)methyl)-4-hydroxytetrahydrofuran-2-yl)-2-oxo-1,2-dihydropyrimidin-4-yl)acetamide (16):**

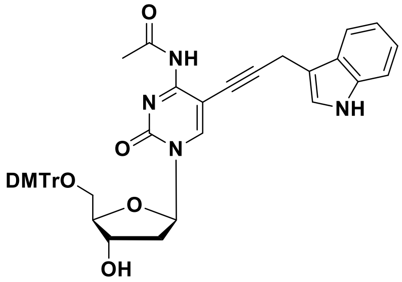


To a solution of compound **15** (1.0 g, 1.5 mmol) in dry DMF (15 mL), acetic anhydride (0.15 mL, 1.6 mmol) was added dropwise over 15 minutes at 0 °C under an argon atmosphere. The reaction mixture was then stirred at room temperature for 10 h. Progress of the reaction was monitored by TLC. After completion of the reaction, the reaction mixture was concentrated under reduced pressure. The crude product was purified by flash chromatography (0–2% MeOH in $CH_2Cl_2$, with 0.2% of $Et_3N$) as the eluent, yielding the desired compound **16** (0.6 g, 0.83 mmol, 57%) as a pale-yellow solid

**$R_f$** (DCM/Methanol 9:1) 0.6;

**$^1$H NMR** (500 MHz, DMSO-$d_6$) $\delta_H$ 10.91 (s, 1H), 9.18 (s, 1H), 8.22 (s, 1H), 7.55 (d, *J* = 7.8 Hz, 1H), 7.40 (d, *J* = 7.7 Hz, 2H), 7.36 (d, *J* = 8.1 Hz, 1H), 7.29 (m, 6H), 7.22 – 7.17 (m, 2H), 7.09 (t, *J* = 7.5 Hz, 1H), 6.97 (t, *J* = 7.4 Hz, 1H), 6.85 (m, 4H), 6.06 (t, *J* = 6.2 Hz, 1H), 5.36 (d, *J* = 4.4 Hz, 1H), 4.28 (s, 1H), 4.03 (s, 1H), 3.72 (s, 2H), 3.68 (d, *J* = 3.4 Hz, 8H), 3.24 (dd, *J* = 10.5, 5.3 Hz, 1H), 3.14 (d, *J* = 8.5 Hz, 1H), 2.42 – 2.35 (m, 1H), 2.28 (s, 3H), 2.24 – 2.17 (m, 1H);

**$^{13}$C NMR** (126 MHz, DMSO-$d_6$) $\delta_C$ 170.5, 161.3, 158.5, 158.5, 152.9, 146.3, 145.2, 136.8, 136.0, 135.7, 130.2, 130.1, 128.4, 128.0, 127.1, 126.8, 123.4, 121.7, 119.0, 118.6, 113.7, 113.7, 112.0, 109.3, 96.1, 94.0, 87.4, 86.8, 86.4, 71.5, 70.7, 63.9, 55.4, 41.4, 25.6, 16.2;

**HRMS** (ESI/Q-TOF) $[M+H]^+$ calcd. for $C_{43}H_{41}N_4O_7$ 725.2975, found 725.2970.

**Synthesis of (2R,3S,5R)-5-(5-(3-(1*H*-indol-3-yl)prop-1-yn-1-yl)-4-acetamido-2-oxopyrimidin-1(2*H*)-yl)-2-((bis(4-methoxyphenyl)(phenyl)methoxy)methyl)tetrahydrofuran-3-yl (2-cyanoethyl) diisopropylphosphoramidite (2b):**

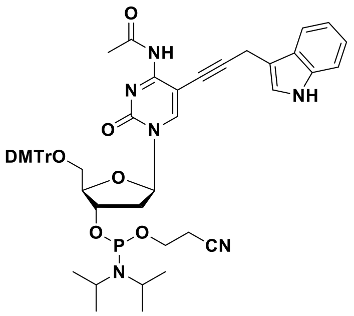


To a solution of compound **16** (0.3 g, 0.41 mmol) in anhydrous $CH_2Cl_2$, *N,N*-diisopropylethylamine (0.36 mL, 2.1 mmol) was added under stirring. The reaction mixture was cooled to 0 °C, and 2-cyanoethyl-*N,N*-diisopropylchlorophosphoramidite (0.14 mL, 0.66 mmol) was added under an argon atmosphere. The mixture was allowed to warm to room temperature and stirred for 1.5 h. The reaction mixture was then diluted with anhydrous $CH_2Cl_2$ (20 mL) and washed successively with 5% $NaHCO_3$ (20 mL) and brine solution (20 mL). The organic phase was dried over anhydrous $Na_2SO_4$ and concentrated under reduced pressure. The crude product was purified by column chromatography using (60% EtOAc in hexane containing 0.5% $Et_3N$) as the eluent. The desired compound **2b** (0.24 g, 0.26 mmol, 64%) was obtained as a white foam.

**$R_f$** (EtOAc/Hexane 7:3) 0.5;

**$^1$H NMR** (500 MHz, $CD_3CN$) $\delta_H$ 9.16 (s, 1H), 8.31 (s, 1H), 7.60 (d, *J* = 7.9 Hz, 1H), 7.51 (d, *J* = 7.4 Hz, 2H), 7.45 (d, *J* = 8.2 Hz, 1H), 7.42 – 7.37 (m, 5H), 7.33 (t, *J* = 7.6 Hz, 2H), 7.25 (t, *J* = 6.8 Hz, 1H), 7.19 (t, *J* = 7.3 Hz, 1H), 7.12 – 7.07 (m, 2H), 6.86 (dd, *J* = 8.8, 5.3 Hz, 4H), 6.10 (t, *J* = 6.1 Hz, 1H), 4.71 (m, 1H), 4.26 (qt, *J* = 4 Hz, 7 Hz, 1H), 3.75 (s, 3H), 3.74 (s, 3H), 3.70 – 3.61 (m, 3H), 3.38 (ddd, *J* = 13.6, 11.0, 3.2 Hz, 2H), 3.29 (br. s, 1H), 2.73 – 2.62 (m, 1H), 2.58 (q, *J* = 6.0 Hz, 4H), 2.49 – 2.42 (m, 1H), 2.40 (s, 3H), 1.23 (s, 3H), 1.22 (s, 6H), 1.21 (s, 3H);

$^{13}$**C NMR** (126 MHz, $CD_3CN$) $\delta_C$ 160.2, 158.7, 158.7, 145.3, 144.9, 136.6, 135.9. 135.5, 130.2, 130.0, 128.0, 128.0, 127.0, 126.6, 122.7, 121.8, 119.2, 118.4, 118.2, 113.2, 113.2, 111.5, 109.6, 96.8, 87.4, 86.7, 86.0, 85.9, 72.3, 72.3, 70.5, 62.5, 60.0, 58.6, 57.2, 54.5, 43.1, 43.0, 40.5, 25.2, 24.0, 44.0, 23.9, 23.9, 21.0, 20.1, 20.0, 15.7, 13.5;

$^{31}$**P NMR** (202 MHz, $CD_3CN$) $\delta_P$ 148.1. 148.1;

**HRMS** (ESI/Q-TOF) $[M+H]^+$ calcd. for $C_{52}H_{57}N_6O_8P$ 925.4048, found 925.4040.

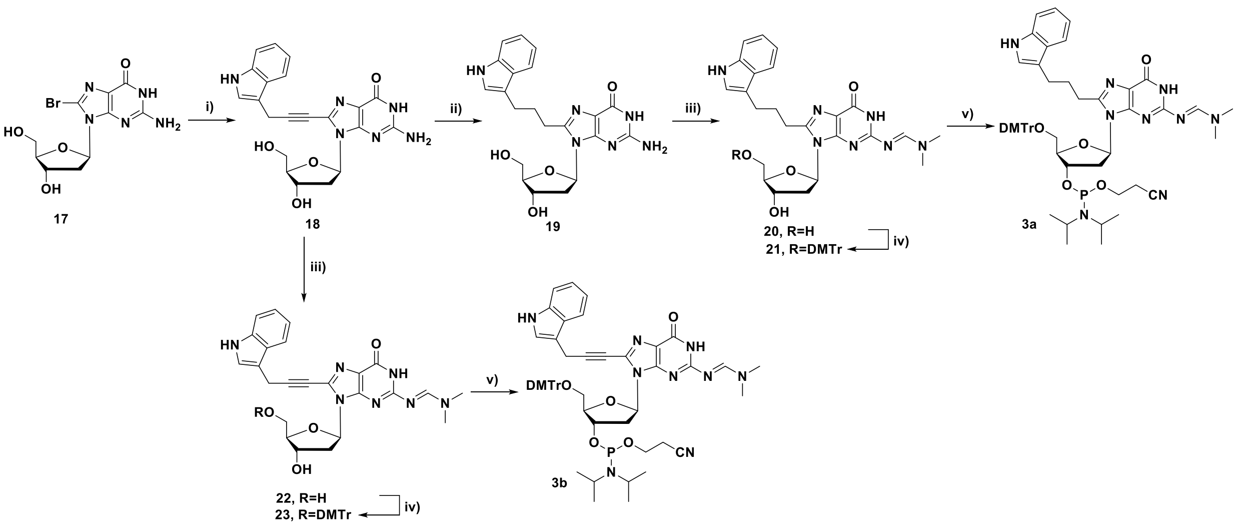

**Scheme 3**: **Reagents and conditions:** (i) $Pd(PPh_3)_4$, CuI, $Et_3N$, 1*H*-Indole,3-(2-propynyl), DMF, 60 °C 3 h (ii) $H_2$, 10% Pd/C, MeOH, 50 °C, overnight (iii) DMF-DMA, DMF, rt, 3h (iv) DMTrCl, DMAP, pyridine, rt, overnight (v) 2-cyanoethyl-*N*,*N*-diisopropylchlorophosphoramidite, DIPEA, DCM, 0 °C to rt, 1.5 h.

**Synthesis of 8-(3-(1H-indol-3-yl)prop-1-yn-1-yl)-2-amino-9-((2R,4S,5R)-4-hydroxy-5-hydroxymethyl)tetrahydrofuran-2-yl)-1,9-dihydro-6H-purin-6-one (18):**

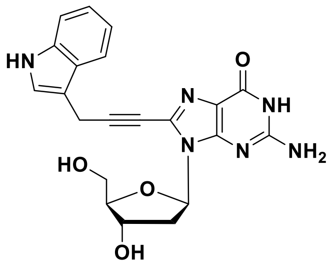

To a solution of 8-Bromo-2'-deoxyguanosine (**17**) (2.0 g, 5.78 mmol) in anhydrous DMF (40 mL), CuI (0.33 g, 1.73 mmol) and TEA (4.03 mL, 28.9 mmol) were added under stirring. After 5 minutes of stirring, 3-(prop-2-yn-1-yl)-1*H*-indole (1.17 g, 7.51 mmol) and $Pd(PPh_3)_4$ (0.67 g, 0.58 mmol) were added to the reaction mixture. The reaction mixture was then heated to 60 °C and stirred for 3 h. The progress of the reaction was followed using TLC. After completion of the reaction, the reaction mixture was diluted with ethyl acetate (50 mL) and filtered through a celite. The filtrate was evaporated under reduced pressure and the crude product was purified by flash chromatography (0–15% MeOH in $CH_2Cl_2$). The desired compound **18** (1.4 g, 3.3 mmol, 58%) was obtained as brown solid.

**$R_f$** (DCM/Methanol 8:2) 0.5;

**$^1$H NMR** (500 MHz, DMSO-$d_6$) $\delta_H$ 10.99 (s, 1H), 7.63 (d, *J* = 7.8 Hz, 1H), 7.39 (d, *J* = 8.1 Hz, 1H), 7.31 (d, *J* = 1.9 Hz, 1H), 7.12 (t, *J* = 7.5 Hz, 1H), 7.04 (t, *J* = 7.4 Hz, 1H), 6.51 (s, 2H), 6.29 (dd, *J* = 7.6, 7.1 Hz, 1H), 5.23 (d, *J* = 4.0 Hz, 1H), 4.87 (t, *J* = 5.8 Hz, 1H), 4.37 – 4.31 (m, 1H), 3.83 – 3.76 (m, 1H), 3.62 – 3.58 (m, 1H), 3.51 – 3.45 (m, 1H), 3.10 – 3.03 (m, 1H), 2.10 – 2.05 (m, 1H).

**$^{13}$C NMR** (126 MHz, DMSO-$d_6$) $\delta_C$ 156.5, 154.2, 151.0, 136.8, 130.3, 126.8, 123.5, 121.8, 119.1, 118.8, 117.4, 112.0, 108.5, 94.3, 88.2, 84.3, 71.6, 71.5, 62.6, 37.4, 16.0.

**HRMS** (ESI/Q-TOF) $[M+H]^+$ calcd. for $C_{21}H_{21}N_6O_4$ 421.1624, found 421.1617.

**Synthesis of 8-(3-(1*H*-indol-3-yl)propyl)-2-amino-9-((2R,4S,5R)-4-hydroxy-5-(hydroxymethyl)tetrahydrofuran-2-yl)-1,9-dihydro-6*H*-purin-6-one (19):**

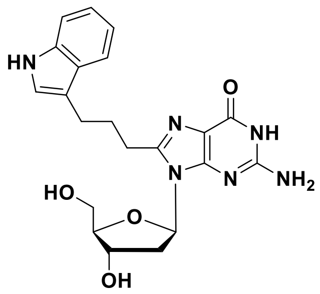


To a solution of compound **18** (1.3 g, 3.1 mmol) in methanol (60 mL), 10% Pd/C (0.33 g, 0.31 mmol) was added under stirring. The reaction mixture was then stirred under a hydrogen atmosphere at 50 °C overnight. The progress of the reaction was followed using TLC. After completion of the reaction, the reaction mixture was filtered through celite to remove the catalyst. The filtrate was evaporated under reduced pressure to yield the pure compound **19** (1.0 g, 2.4 mmol, 76 %) as a yellow solid.

**$R_f$** (DCM:Methanol 8:2) 0.5;

**$^1$H NMR** (500 MHz, DMSO-$d_6$) $\delta_H$ 10.77 (s, 1H), 10.61 (s, 1H), 7.53 (d, *J* = 7.9 Hz, 1H), 7.34 (d, *J* = 8.0 Hz, 1H), 7.14 (s, 1H), 7.06 (t, *J* = 7.5 Hz, 1H), 6.96 (t, *J* = 7.4 Hz, 1H), 6.32 (s, 2H), 6.13 (t, *J* = 10.0 Hz, 1H), 5.23 (s, 1H), 4.99 (s, 1H), 4.35 (s, 1H), 3.79 (d, *J* = 2.9 Hz, 1H), 3.60 (d, *J* = 11.4 Hz, 1H), 3.49 (dd, *J* = 10.7, 4.9 Hz, 1H), 3.17 (s, 1H), 2.85 – 2.75 (m, 4H), 2.11 – 1.99 (m, 3H);

**$^{13}$C NMR** (126 MHz, DMSO-$d_6$) $\delta_C$ 156.8, 153.2, 152.0, 148.6, 136.8, 127.6, 122.7, 121.3, 118.8, 118.6, 116.0, 114.6, 111.8, 88.0, 83.7, 71.5, 62.5, 49.1, 28.2, 27.7, 24.7;

**HRMS** (ESI/Q-TOF) $[M+H]^+$ calcd. for $C_{21}H_{25}N_6O_4$ 425.1932, found 425. 1932.

**Synthesis of (E)-N'-(8-(3-(1*H*-indol-3-yl)propyl)-9-((2R,4S,5R)-4-hydroxy-5-(hydroxymethyl)tetrahydrofuran-2-yl)-6-oxo-6,9-dihydro-1*H*-purin-2-yl)-N,N-dimethylformimidamide (20):**

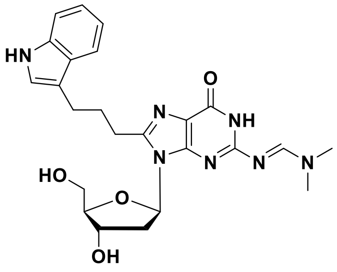


To a solution of compound **19** (0.80 g, 1.88 mmol) in anhydrous DMF (10 mL) under argon atmosphere, dimethylformamide dimethylacetal (5.0 mL, 37.7 mmol) was added under stirring. The reaction mixture was continued to stir at room temperature for 3 h. The progress of the reaction was followed using TLC. After the reaction was complete, the solvent was removed under reduced pressure. The crude product was purified by flash chromatography using (0–10% MeOH in DCM) as eluent. The pure product **20** (0.66 g, 1.4 mmol, 73%) was obtained as a white solid.

**R$_f$** (DCM/Methanol 9:1) 0.4;

**$^1$H NMR** (500 MHz, DMSO-*d$_6$*) $\delta_H$ 11.30 (s, 1H), 10.77 (s, 1H), 8.46 (s, 1H), 7.53 (d, *J* = 7.8 Hz, 1H), 7.34 (d, *J* = 8.1 Hz, 1H), 7.15 (d, *J* = 2.0 Hz, 1H), 7.06 (dd, *J* = 11.1, 4.0 Hz, 1H), 6.97 (t, *J* = 7.4 Hz, 1H), 6.18 (t, *J* = 7.3 Hz, 1H), 5.29 (d, *J* = 4.4 Hz, 1H), 4.90 (dd, *J* = 6.7, 5.0 Hz, 1H), 4.42 (dt, *J* = 7.2, 3.6 Hz, 1H), 3.80 (dd, *J* = 7.9, 4.4 Hz, 1H), 3.63 (dt, *J* = 11.5, 4.6 Hz, 1H), 3.51 (ddd, *J* = 11.7, 6.7, 5.0 Hz, 1H), 3.13 (s, 3H), 3.02 (s, 3H), 2.97 (dt, *J* = 14.2, 7.0 Hz, 1H), 2.82 (m, 4H), 2.09 (m, 3H);

**$^{13}$C NMR** (126 MHz, DMSO-*d$_6$*) $\delta_C$ 158.4, 157.7, 156.9, 150.7, 149.7, 136.8, 127.8, 122.8, 121.3, 119.2, 118.8, 118.6, 114.5, 111.8, 87.9, 83.8, 71.3, 62.4, 41.2, 38.4, 35.0, 28.2, 27.7, 24.7;

**HRMS** (ESI/Q-TOF) [M+H]$^+$ calcd. for $C_{24}H_{30}N_7O_4$ 480.2353, found 480.2348.

**Synthesis of (E)-N'-(8-(3-(1*H*-indol-3-yl)propyl)-9-((2R,4S,5R)-5-((bis(4-methoxyphenyl)(phenyl)methoxy)methyl)-4-hydroxytetrahydrofuran-2-yl)-6-oxo-6,9-dihydro-1*H*-purin-2-yl)-N,N-dimethylformimidamide (21):**

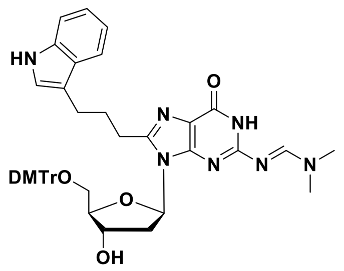


To a solution of compound **20** (0.5 g, 1.04 mmol) in anhydrous pyridine (10 mL), DMAP (13 mg, 0.10 mmol) was added. To this mixture, a solution of 4,4'-dimethoxytrityl chloride (0.42 g, 1.25 mmol) in 3 mL of anhydrous pyridine was added in four equal portions over the time of 1 h. The reaction mixture was stirred at room temperature for 12 h. After completion of the reaction, the solvent was evaporated under reduced pressure. The resulting crude mixture was purified by flash chromatography (0–2% MeOH in $CH_2Cl_2$, with 0.2% of $Et_3N$) as the eluent, yielding the desired compound **21** (0.43 g, 0.55 mmol, 53%) as a pale-yellow solid.

**R$_f$** (DCM/Methanol 9:1) 0.4;

**$^1$H NMR** (500 MHz, DMSO-*d$_6$*) $\delta_H$ 11.28 (s, 1H), 10.76 (s, 1H), 8.32 (s, 1H), 7.50 (d, *J* = 7.9 Hz, 1H), 7.33 (d, *J* = 8.1 Hz, 1H), 7.29 (d, *J* = 7.0 Hz, 2H), 7.17 (m 7H), 7.10 (d, *J* = 1.7 Hz, 1H), 7.05 (t, *J* = 7.6 Hz, 1H), 6.94 (t, *J* = 7.3 Hz, 1H), 6.77 (d, *J* = 8.8 Hz, 2H), 6.73 (d, *J* = 8.8 Hz, 2H), 6.28 – 6.22 (m, 1H), 5.32 (d, *J* = 4.9 Hz, 1H), 4.55 (dt, *J* = 11.7, 5.7 Hz, 1H), 3.87 (td, *J* = 6.4, 3.7 Hz, 1H), 3.70 (s, 3H), 3.69 (s, 3H), 3.19 (dt, *J* = 11.9, 6.0 Hz, 1H), 3.12 (dd, *J* = 10.1, 2.7 Hz, 1H), 3.07 (dd, *J* = 12.7, 6.7 Hz, 1H), 3.01 (s, 6H), 2.88 (t, *J* = 7.4 Hz, 2H), 2.77 (t, *J* = 7.5 Hz, 2H), 2.23 – 2.07 (m, 3H);

**$^{13}$C NMR** (126 MHz, DMSO-*d$_6$*) $\delta_C$ 158.4, 158.3, 157.9, 157.7, 156.5, 150.5, 149.9, 145.4, 136.8, 136.2, 136.0, 130.1, 129.9, 128.1, 127.6, 127.0, 122.6, 121.3, 119.2, 118.8, 118.6, 114.6, 113.4, 113.4, 111.8, 85.7, 83.2, 70.9, 64.3, 55.4, 55.4, 41.2, 38.3, 35.0, 28.1, 27.7, 24.8;

**HRMS** (ESI/Q-TOF) [M+H]$^+$ calcd. for $C_{45}H_{48}N_7O_6$ 782.3661, found 782.3659.

**Synthesis of (2R,3S,5R)-5-(8-(3-(1*H*-indol-3-yl)propyl)-2-(((E)-(dimethylamino)methylene)amino)-6-oxo-1,6-dihydro-9*H*-purin-9-yl)-2-((bis(4-methoxyphenyl)(phenyl)methoxy)methyl)tetrahydrofuran-3-yl (2-cyanoethyl) diisopropylphosphoramidite (3a):**

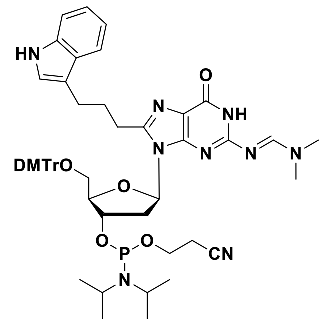


To a solution of compound **21** (0.35 g, 0.45 mmol) in anhydrous $CH_2Cl_2$, *N,N*-diisopropylethylamine (0.39 mL, 2.24 mmol) was added under stirring. The reaction mixture was cooled to 0 °C, and 2-cyanoethyl-*N,N*-diisopropylchlorophosphoramidite (0.15 mL, 0.72 mmol) was added under an argon atmosphere. The mixture was allowed to warm to room temperature and stirred for 1.5 h. The reaction mixture was then diluted with anhydrous $CH_2Cl_2$ (20 mL) and washed successively with 5% $NaHCO_3$ (20 mL) and brine solution (20 mL). The organic phase was dried over anhydrous $Na_2SO_4$ and concentrated under reduced pressure. The resulting crude mixture was purified by flash chromatography (30–40% acetone in hexane containing 0.5% $Et_3N$) as the eluent. The desired compound **3a** (0.24 g, 0.22 mmol, 55%) was obtained as a pale-yellow solid.

**$R_f$** (EtOAc/Hexane, 9:1) 0.4;

**$^1$H NMR** (500 MHz, $CD_3CN$) $\delta_H$ 9.52 (br. s, 1H), 9.13 (br. s, 1H), 8.42 (d, *J* = 3.4 Hz, 1H), 7.60 (dd, *J* = 7.7, 4.6 Hz, 1H), 7.42 (d, *J* = 8.1 Hz, 1H), 7.36 (m, 2H), 7.27 – 7.17 (m, 7H), 7.17 – 7.12 (m, 1H), 7.11 (d, *J* = 2.0 Hz, 1H), 7.07 – 7.01 (m, 1H), 6.76 (m, 4H), 6.21 (m, 1H), 5.08 – 4.94 (m, 1H), 4.14 – 4.04 (m, 1H), 3.75 (m, 6H), 3.67 – 3.53 (m, 3H), 3.36 – 3.20 (m, 3H), 3.09 – 3.04 (m, 6H), 2.97 – 2.86 (m, 4H), 2.65 – 2.60 (m, 1H), 2.48 (t, *J* = 6.0 Hz, 1H), 2.43 – 2.31 (m, 1H), 2.31 – 2.25 (m, 3H), 1.23 (br. s, 3H), 1.20 (d, *J* = 6.8 Hz, 3H), 1.16 (m, 6H), 1.04 (d, *J* = 6.8 Hz, 3H);

**$^{13}$C NMR** (126 MHz, $CD_3CN$) $\delta_C$ 158.6, 158.5, 158.5, 157.6, 157.4, 156.1, 150.6, 150.3, 145.2, 145.1, 136.6, 135.9, 135.8, 129.9, 129.7, 127.9, 127.7, 127.5, 126.7, 122.2, 121.3, 119.3, 118.6, 118.6, 114.9, 112.9, 111.3, 85.8, 84.6, 83.4, 83.3, 63.8, 63.5, 58.7, 58.2, 54.9, 54.8, 43.2, 43.1, 40.7, 34.3, 27.8, 27.8, 27.4, 24.3, 23.9, 23.9, 23.9, 23.9, 23.9, 23.8, 23.8, 20.1, 19.9;

**$^{31}$P NMR** (202 MHz, $CD_3CN$) $\delta_P$ 149.8, 149.6;

**HRMS** (ESI/Q-TOF) $[M+H]^+$ calcd. for $C_{54}H_{65}N_9O_7P$ 982.4739, found 982.4736.

**Synthesis of (E)-N'-(8-(3-(1*H*-indol-3-yl)prop-1-yn-1-yl)-9-((2R,4S,5R)-4-hydroxy-5-(hydroxymethyl)tetrahydrofuran-2-yl)-6-oxo-6,9-dihydro-1*H*-purin-2-yl)-N,N-dimethylformimidamide (22):**

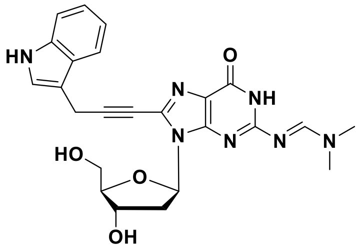


To a solution of compound **18** (0.7 g, 1.66 mmol) in anhydrous DMF (8 mL) under argon atmosphere, dimethylformamide dimethylacetal (4.4 mL, 33.3 mmol) was added under stirring. The reaction mixture was stirred at room temperature for 3 h. The progress of the reaction was followed using TLC. After the reaction was complete, the solvent was removed under reduced pressure. The crude product was purified by flash chromatography using (0–10% MeOH in DCM as the eluent. The pure compound **22** (0.52 g, 1.1 mmol, 66%) was obtained as a white solid.

**R$_f$** (DCM/Methanol, 9:1) 0.5;

**$^1$H NMR** (500 MHz, DMSO-*d$_6$*) $\delta_H$ 11.49 (s, 1H), 10.99 (s, 1H), 8.50 (s, 1H), 7.64 (d, *J* = 7.8 Hz, 1H), 7.39 (d, *J* = 8.1 Hz, 1H), 7.32 (s, 1H), 7.12 (t, *J* = 7.5 Hz, 1H), 7.05 (t, *J* = 7.4 Hz, 1H), 6.38 (t, *J* = 7.2 Hz, 1H), 5.31 (d, *J* = 4.1 Hz, 1H), 4.84 (t, *J* = 5.9 Hz, 1H), 4.43 (td, *J* = 6.6, 3.5 Hz, 1H), 3.82 (dd, *J* = 8.2, 4.8 Hz, 1H), 3.68 – 3.59 (m, 1H), 3.57 – 3.47 (m, 1H), 3.15 (s, 3H), 3.10 – 3.06 (m, 1H), 3.04 (s, 3H), 2.13 (ddd, *J* = 12.8, 6.5, 3.0 Hz, 1H).

**$^{13}$C NMR** (126 MHz, DMSO-*d$_6$*) $\delta_C$ 158.7, 158.0, 157.4, 149.7, 136.8, 131.5, 126.8, 123.6, 121.8, 120.5, 119.12, 118.8, 112.1, 108.4, 95.0, 88.2, 84.6, 71.3, 71.2, 62.5, 41.3, 38.0, 35.1, 16.0;

**HRMS** (ESI/Q-TOF) $[M+H]^+$ calcd. for $C_{24}H_{26}N_7O_4$ 476.2046, found 476.2037.

**Synthesis of (E)-N'-(8-(3-(1*H*-indol-3-yl)prop-1-yn-1-yl)-9-((2R,4S,5R)-5-((bis(4-methoxyphenyl)(phenyl)methoxy)methyl)-4-hydroxytetrahydrofuran-2-yl)-6-oxo-6,9-dihydro-1*H*-purin-2-yl)-N,N-dimethylformimidamide (23):**

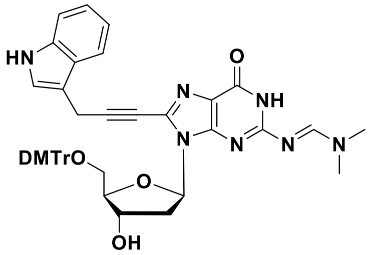


To a solution of compound **22** (0.4 g, 0.84 mmol) in anhydrous pyridine (8 mL), DMAP (10 mg, 0.1 mmol) was added under stirring. To this mixture, a solution of 4,4'-dimethoxytrityl chloride (0.34 g, 1.01 mmol) in 4 mL of anhydrous pyridine was added in four equal portions over the time of 1 h. The reaction mixture was stirred at room temperature for 12 h. After completion of the reaction, the solvent was evaporated under reduced pressure. The resulting crude residue was purified by flash chromatography (0–2% MeOH in $CH_2Cl_2$, with 0.2% of $Et_3N$) as a eluent, yielding the desired compound **23** (0.38 g, 0.49 mmol, 58%) as a pale-yellow solid.

**R$_f$** (DCM/Methanol, 9:1) 0.5;

**$^1$H NMR** (500 MHz, DMSO-*d$_6$*) $\delta_H$ 11.46 (s, 1H), 10.98 (s, 1H), 8.33 (s, 1H), 7.61 (d, *J* = 7.9 Hz, 1H), 7.40 (d, *J* = 8.1 Hz, 1H), 7.30 (d, *J* = 6.8 Hz, 2H), 7.25 (d, *J* = 1.9 Hz, 1H), 7.22 – 7.10 (m, 9H), 7.06 – 7.01 (m, 1H), 6.76 (d, *J* = 8.9 Hz, 2H), 6.72 (d, *J* = 8.9 Hz, 2H), 6.42 (dd, *J* = 7.5, 5.7 Hz, 1H), 5.36 (d, *J* = 4.8 Hz, 1H), 3.25 (dd, *J* = 9.8, 7.4 Hz, 1H), 3.15 – 3.07 (m, 2H), 3.02 (s, 6H), 2.22 (ddd, *J* = 13.2, 7.6, 5.6 Hz, 1H);

**$^{13}$C NMR** (126 MHz, DMSO-*d$_6$*) $\delta_C$ 158.4, 158.4, 158.2, 157.7, 157.5, 149.6, 145.4, 136.9, 136.1, 131.5, 130.1, 130.0, 128.1, 127.0, 126.8, 123.5, 121.8, 120.4, 119.1, 118.8, 113.5, 113.4, 112.1, 108.5, 94.7, 86.3, 85.7, 84.0, 71.7, 71.3, 64.7, 55.4, 55.4, 35.1, 16.0;

**HRMS** (ESI/Q-TOF) $[M+H]^+$ calcd. for $C_{45}H_{44}N_7O_6$ 778.3353, found 778.3357.

**Synthesis of (2R,3S,5R)-5-(8-(3-(1*H*-indol-3-yl)prop-1-yn-1-yl)-2-(((E)-(dimethylamino)methylene)amino)-6-oxo-1,6-dihydro-9*H*-purin-9-yl)-2-((bis(4-methoxyphenyl)(phenyl)methoxy)methyl)tetrahydrofuran-3-yl (2-cyanoethyl) diisopropylphosphoramidite (3b):**

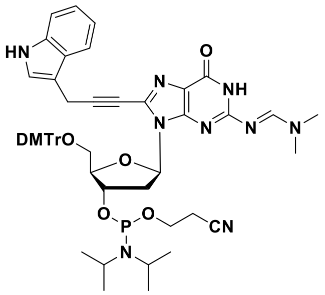

To a solution of compound **23** (0.3 g, 0.39 mmol) in anhydrous $CH_2Cl_2$, *N,N*-diisopropylethylamine (0.34 mL, 1.93 mmol) was added under stirring. The reaction mixture was cooled to 0 °C, and 2-cyanoethyl-*N,N*-diisopropylchlorophosphoramidite (0.13 mL, 0.62 mmol) was added under an argon atmosphere. The mixture was allowed to warm to room temperature and stirred for 1.5 h. The reaction mixture was then diluted with anhydrous $CH_2Cl_2$ (20 mL) and washed successively with 5% $NaHCO_3$ (20 mL) and brine solution (20 mL). The organic phase was dried over anhydrous $Na_2SO_4$ and concentrated under reduced pressure. The crude residue was purified by column chromatography using (30–40% acetone in hexane containing 0.5% $Et_3N$) as the eluent. The desired compound **3b** (0.26 g, 0.27 mmol, 69%) was obtained as a pale-yellow solid.

**$R_f$** (EtOAc/Hexane, 9:1) 0.4;

**$^1$H NMR** (500 MHz, $CD_3CN$) $\delta_H$ 9.42 (s, 1H), 9.29 (s, 1H), 8.42 (d, *J* = 6.4 Hz, 1H), 7.71 (d, *J* = 7.9 Hz, 1H), 7.49 (d, *J* = 8.2 Hz, 1H), 7.38 (m, 2H), 7.29 (t, *J* = 2.9 Hz, 1H), 7.27 (d, *J* = 1.5 Hz, 1H), 7.26 – 7.19 (m, 7H), 7.17 – 7.12 (m, 1H), 6.82 – 6.72 (m, 4H), 6.52 (td, *J* = 8.1, 4.9 Hz, 1H), 4.98-4.88 (m, 1H), 4.16 – 4.07 (m, 2H), 4.00 (s, 2H), 3.77 (d, *J* = 3.6 Hz, 3H), 3.76 (d, *J* = 2.7 Hz, 3H), 3.72 (ddd, *J* = 10.4, 7.5, 6.0 Hz, 1H), 3.65 – 3.53 (m, 3H), 3.36 – 3.25 (m, 2H), 3.22 (dd, *J* = 10.1, 3.0 Hz, 1H), 3.08 (m, 6H), 2.62 (t, *J* = 6.0 Hz, 1H), 2.50 – 2.42 (m, 1H), 1.19-1.13 (m, 9H), 1.03 (d, *J* = 6.8 Hz, 3H);

**$^{13}$C NMR** (126 MHz, $CD_3CN$) $\delta_C$ 158.6, 158.5, 158.0, 157.3, 157.1, 149.6, 145.2, 136.7, 135.9, 135.8, 132.1, 129.9, 129.8, 127.9, 127.8, 127.7, 126.7, 126.7, 126.6, 122.9, 121.9, 120.4, 119.9, 118.6, 118.4, 112.9, 111.6, 108.9, 94.3, 85.8, 84.9, 84.2, 74.1, 71.1, 64.1, 59.9, 58.4, 58.3, 54.8, 54.8, 43.2, 43.1, 40.9, 40.9, 37.0, 34.4, 23.9, 23.9, 23.8, 23.8, 20.1, 20.0, 15.8, 13.5;

**$^{31}$P NMR** (202 MHz, $CD_3CN$) $\delta_P$ 149.9, 149.8;

**HRMS** (ESI/Q-TOF) $[M+H]^+$ calcd. for $C_{54}H_{61}N_9O_7P$ 978.4426, found 978.4431.

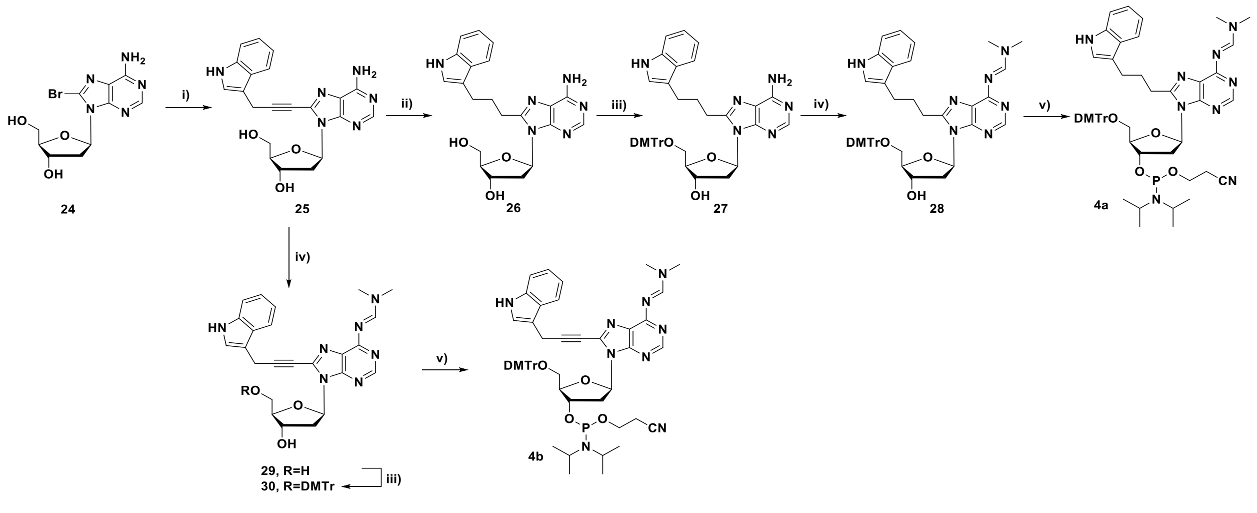

**Scheme 4**: **Reagents and conditions:** (i) $Pd(PPh_3)_4$, CuI, $Et_3N$, 1*H*-Indole,3-(2-propynyl), DMF, 60 °C 2 h (ii) $H_2$, 10% Pd/C, MeOH, 50 °C, overnight (iii) DMTrCl, DMAP, pyridine, rt, overnight (iv) DMF-DMA, DMF, rt, 4h (v) 2-cyanoethyl-*N*,*N*-diisopropylchlorophosphoramidite, DIPEA, DCM, 0 °C to rt, 1.5 h.

**Synthesis of (2R,3S,5R)-5-(8-(3-(1*H*-indol-3-yl)prop-1-yn-1-yl)-6-amino-9*H*-purin-9-yl)-2-(hydroxymethyl)tetrahydrofuran-3-ol (25):**

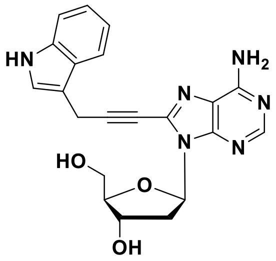

To a solution of 8-Bromo-2'-deoxyadenosine (**24**) (2.0 g, 6.1 mmol) in anhydrous DMF (30 mL), CuI (0.35 g, 1.8 mmol) and TEA (3.4 mL, 24.0 mmol) were added under stirring. After 5 minutes of stirring, 3-(prop-2-yn-1-yl)-1*H*-indole (1.2 g, 7.9 mmol) and $Pd(PPh_3)_4$ (0.7 g, 0.61 mmol) were added to the reaction mixture. The reaction mixture was heated to 60 °C and stirred for 2 h. After completion of the reaction, the reaction mixture was diluted with ethyl acetate (50 mL) and filtered through a celite. The filtrate was evaporated under reduced pressure and the crude product was purified by flash chromatography (0–10% MeOH in $CH_2Cl_2$). The desired compound **25** (1.4 g, 3.5 mmol, 57%) was obtained as a yellow solid.

**$R_f$** (DCM/Methanol 9:1) 0.4;

**$^1$H NMR** (500 MHz, DMSO-*d₆*) $\delta_H$ 11.01 (s, 1H), 8.13 (s, 1H), 7.64 (d, *J* = 7.9 Hz, 1H), 7.50 (s, 2H), 7.40 (d, *J* = 8.1 Hz, 1H), 7.33 (d, *J* = 2.0 Hz, 1H), 7.13 (t, *J* = 7.4 Hz, 1H), 7.05 (t, *J* = 7.4 Hz, 1H), 6.48 (dd, *J* = 7.9, 6.7 Hz, 1H), 5.37 (dd, *J* = 7.9, 4.3 Hz, 1H), 5.33 (d, *J* = 4.0 Hz, 1H), 4.50 – 4.46 (m, 1H), 4.10 (s, 2H), 3.91 (dd, *J* = 6.7, 4.3 Hz, 1H), 3.67 (dt, *J* = 11.8, 4.3 Hz, 1H), 3.53 – 3.47 (m, 1H), 3.15 (m, 1H), 2.18 (ddd, *J* = 12.8, 6.2, 2.2 Hz, 1H);

**$^{13}$C NMR** (126 MHz, DMSO-$d_6$) $\delta_C$ 156.37, 153.48, 148.83, 136.81, 133.96, 126.80, 123.64, 121.84, 119.64, 119.16, 118.78, 112.02, 108.26, 96.52, 88.80, 85.66, 71.79, 70.61, 62.69, 38.03, 16.06;

**HRMS** (ESI/Q-TOF) $[M+H]^+$ calcd. for $C_{21}H_{21}N_6O_3$ 405.1675, found 405.1672.

**Synthesis of (2R,3S,5R)-5-(8-(3-(1*H*-indol-3-yl)propyl)-6-amino-9*H*-purin-9-yl)-2-(hydroxymethyl)tetrahydrofuran-3-ol (26):**

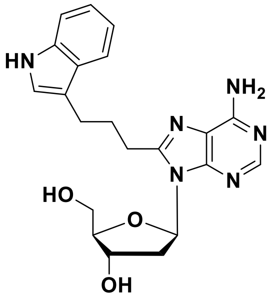

To a solution of compound **25** (1.1 g, 2.7 mmol) in methanol (40 mL), 10% Pd/C (0.29 g, 0.27 mmol) was added under stirring. The reaction mixture was then stirred under a hydrogen atmosphere at 50 °C overnight. After completion of the reaction, the reaction mixture was filtered through celite to remove the catalyst. The filtrate was evaporated under reduced pressure to yield the pure compound **26** (0.85 g, 2.1 mmol, 77 %) as a white solid.

**$R_f$** (DCM/Methanol 9:1) 0.4;

**$^1$H NMR** (500 MHz, DMSO-$d_6$) $\delta_H$ 10.79 (s, 1H), 8.05 (s, 1H), 7.52 (d, *J* = 7.7 Hz, 1H), 7.34 (d, *J* = 8.1 Hz, 1H), 7.17 (m, 3H), 7.06 (d, *J* = 7.9 Hz, 1H), 6.97 (t, *J* = 7.4 Hz, 1H), 6.28 (t, *J* = 7.2 Hz, 1H), 5.58 (dd, *J* = 8.1, 3.6

Hz, 1H), 5.29 (d, $J$ = 4.1 Hz, 1H), 4.46 (s, 1H), 3.90 (s, 1H), 3.66 (dt, $J$ = 11.0, 3.2 Hz, 1H), 3.55 – 3.47 (m, 1H), 3.17 (d, $J$ = 5.1 Hz, 1H), 3.13 – 3.05 (m, 1H), 2.96 (t, $J$ = 7.5 Hz, 2H), 2.82 (t, $J$ = 7.4 Hz, 2H), 2.14 (m, 2H);

**$^{13}$C NMR** (126 MHz, DMSO-$d_6$) $\delta_C$ 155.9, 152.5, 151.7, 150.2, 136.8, 127.6, 122.9, 121.3, 118.8, 118.7, 118.6, 114.3, 111.8, 88.7, 84.9, 71.8, 62.7,38.4, 28.8, 27.7, 24.8;

**HRMS** (ESI/Q-TOF) $[M+H]^+$ calcd. for $C_{21}H_{25}N_6O_3$ 409.1983, found 409.1990.

**Synthesis of (2R,3S,5R)-5-(8-(3-(1*H*-indol-3-yl)propyl)-6-amino-9*H*-purin-9-yl)-2-((bis(4-methoxyphenyl)(phenyl)methoxy)methyl)tetrahydrofuran-3-ol (27):**

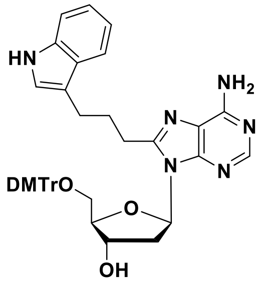


To a solution of compound **26** (0.75 g, 1.83 mmol) in anhydrous pyridine (12 mL), DMAP (22 mg, 0.183 mmol) was added. To this mixture, a solution of 4,4'-dimethoxytrityl chloride (0.74 g, 2.19 mmol) in 4 mL of anhydrous pyridine was added in four equal portions over the time of 1 h. The reaction mixture was stirred at room temperature for 12 h. After completion of the reaction, the solvent was evaporated under reduced pressure. The resulting crude mixture was purified by flash chromatography (0–2% MeOH in $CH_2Cl_2$ with 0.2% of $Et_3N$) yielding the desired compound **27** (0.67 g, 0.94 mmol, 52%) as a yellow solid.

**$R_f$** (DCM/methanol 9:1) 0.5;

**$^1$H NMR** (500 MHz, DMSO-$d_6$) $\delta_H$ 10.77 (s, 1H), 8.58 (d, $J$ = 4.0 Hz, 2H), 7.79 (t, $J$ = 7.6 Hz, 1H), 7.70 (s, 1H), 7.55 (d, $J$ = 8.0 Hz, 1H), 7.42 – 7.36 (m, 2H), 7.33 (d, $J$ = 8.0 Hz, 1H), 7.26 (dd, $J$ = 7.2, 5.7 Hz, 2H), 7.19 – 7.09 (m, 7H), 7.05 (t, $J$ = 7.4 Hz, 1H), 6.94 – 6.86 (m, 1H), 6.82 (d, $J$ = 8.7 Hz, 1H), 6.77 (dd, $J$ = 8.7, 4.3 Hz, 1H), 6.30 (t, $J$ = 6.5 Hz, 1H), 5.31 (d, $J$ = 4.8 Hz, 1H), 4.61 – 4.53 (m, 1H), 3.96 (dd, $J$ = 8.9, 4.4 Hz, 1H), 3.71 (s, 3H), 3.67 (s, 3H), 3.29 (t, $J$ = 7.2 Hz, 1H), 3.10 (ddd, $J$ = 15.9, 10.1, 5.0 Hz, 2H), 3.03 (t, $J$ = 7.9 Hz, 2H), 2.83 (t, $J$ = 7.6 Hz, 2H), 2.24 – 2.11 (m, 3H);

**$^{13}$C NMR** (126 MHz, DMSO-$d_6$) $\delta_C$ 1584, 158.1, 153.9, 153.2, 150.6, 150.0, 149.5, 145.9, 145.5, 137.9, 136.8, 136.6, 136.3, 136.0, 130.2, 130.2, 130.1, 130.1, 128.9, 128.1, 128.0, 127.6, 126.9, 124.4, 122.8, 121.3, 120.3, 118.8, 118.6, 114.4, 113.5, 111.8, 86.1, 85.7, 84.1, 71.3, 70.0, 55.4, 37.2, 28.7, 27.8, 24.9;

**HRMS** (ESI/Q-TOF) $[M+H]^+$ calcd. for $C_{42}H_{43}N_6O_5$ 711.3289, found 711.3291.

**Synthesis of (E)-N'-(8-(3-(1*H*-indol-3-yl)propyl)-9-((2R,4S,5R)-5-((bis(4-methoxyphenyl)(phenyl)methoxy)methyl)-4-hydroxytetrahydrofuran-2-yl)-9*H*-purin-6-yl)-N,N-dimethylformimidamide (28):**

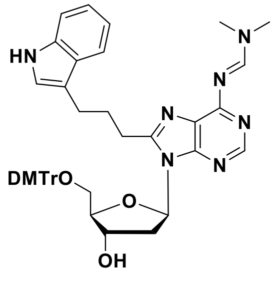


To a solution of compound **27** (0.6 g, 0.88 mmol) in anhydrous DMF (8 mL) under argon atmosphere, dimethylformamide dimethylacetal (2.24 mL, 16.9 mmol) was added. The reaction mixture was stirred at room temperature for 4 h. After the reaction was complete, the solvent was removed under reduced pressure. The crude product was purified by flash chromatography (0–5% MeOH in $CH_2Cl_2$ with 0.2% of $Et_3N$). The pure product **28** (0.44 g, 0.57 mmol, 68%) was obtained as a pale-yellow solid.

**$R_f$** (DCM/Methanol 9;1) 0.55;

**$^1$H NMR** (500 MHz, $CDCl_3$) $\delta_H$ 8.85 (s, 1H), 8.27 (s, 1H), 7.96 (s, 1H), 7.62 (d, *J* = 7.8 Hz, 1H), 7.41 (d, *J* = 7.3 Hz, 2H), 7.36 (d, *J* = 8.1 Hz, 1H), 7.30 (m, 5H), 7.25 (t, *J* = 7.3 Hz, 2H), 7.20 (t, *J* = 7.1 Hz, 2H), 7.11 (t, *J* = 7.4 Hz, 1H), 7.00 (d, *J* = 1.5 Hz, 1H), 6.78 (dd, *J* = 8.7, 6.6 Hz, 4H), 6.12 (t, *J* = 6.9 Hz, 1H), 4.77 – 4.70 (m, 1H), 3.97 (dd, *J* = 10.2, 5.8 Hz, 1H), 3.78 (s, 6H), 3.46 – 3.36 (m, 3H), 3.25 (s, 3H), 3.18 (s, 3H), 3.01 (t, *J* = 8.0 Hz, 2H), 2.92 (t, *J* = 7.2 Hz, 2H), 2.43 – 2.22 (m, 3H), 2.18 – 2.09 (m, 1H);

**$^{13}$C NMR** (126 MHz, $CDCl_3$) $\delta_C$ 160.1, 158.4, 158.4, 157.8, 154.8, 152.6, 151.3, 144.8, 136.4, 136.0, 136.1, 130.1, 130.0, 128.2, 127.8, 127.4, 126.8, 125.5, 121.9, 119.2, 119.0, 115.4, 113.1, 111.2, 86.3, 95.4, 83.7, 73.1, 64.0, 55.2, 41.2, 37.1, 35.2, 28.1, 27.9, 24.8;

**HRMS** (ESI/Q-TOF) $[M+H]^+$ calcd. for $C_{45}H_{48}N_7O_5$ 766.3711, found 766.3688.

**Synthesis of (2R,3S,5R)-5-(8-(3-(1*H*-indol-3-yl)propyl)-6-(((E)-(dimethylamino)methylene)amino)-9*H*-purin-9-yl)-2-((bis(4-methoxyphenyl)(phenyl)methoxy)methyl)tetrahydrofuran-3-yl (2-cyanoethyl) diisopropylphosphoramidite (4a):**

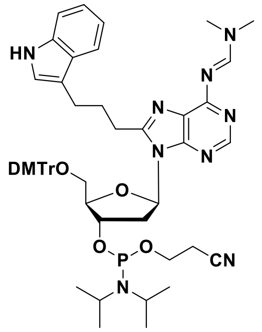


To a solution of compound **28** (0.35 g, 0.46 mmol) in anhydrous $CH_2Cl_2$, *N,N*-diisopropylethylamine (0.4 mL, 2.28 mmol) was added under stirring. The reaction mixture was cooled to 0 °C, and 2-cyanoethyl-*N,N*-diisopropylchlorophosphoramidite (0.16 mL, 0.73 mmol) was added under an argon atmosphere. The mixture was allowed to warm to room temperature and stirred for 1.5 h. The reaction mixture was then diluted with anhydrous $CH_2Cl_2$ (20 mL) and washed successively with 5% $NaHCO_3$ (20 mL) and brine solution (20 mL). The organic phase was dried over anhydrous $Na_2SO_4$ and concentrated under reduced pressure. The crude product was purified by column chromatography using (20–30% acetone in hexane containing 0.5% $Et_3N$) as the eluent. The desired compound **4a** (0.28 g, 0.29 mmol, 63%) was obtained as a white foam.

**$R_f$** (EtOAc/Hexane 9:1) 0.35;

**$^1$H NMR** (500 MHz, $CD_3CN$) $\delta_H$ 9.09 (br. s, 1H), 8.87 (s, 1H), 8.26 (m, 2H), 7.64 (d, *J* = 7.9 Hz, 1H), 7.43 (d, *J* = 8.1 Hz, 1H), 7.34 (m, 3H), 7.25 – 7.17 (m, 4H), 7.16 – 7.11 (m, 2H), 7.05 (t, *J* = 7.5 Hz, 1H), 6.81 – 6.67 (m, 5H), 6.29 (td, *J* = 7.9, 4.9 Hz, 1H), 5.23 – 5.03 (m, 1H), 3.92 – 3.80 (m, 1H), 3.76 (s, 3H), 3.75 (s, 3H), 3.71 (m, 1H), 3.60 (m, 4H), 3.38 – 3.28 (m, 1H), 3.23 (m, 1H), 3.19 (s, 3H), 3.18 (s, 3H), 3.15 – 3.11 (m, 1H), 3.08 (td, *J* = 7.6, 1.9 Hz, 3H), 2.97 – 2.88 (m, 3H), 2.68 (m, 1H), 2.57 (t, *J* = 6.0 Hz, 1H), 2.48 – 2.39 (m, 1H), 2.38 – 2.25 (m, 2H), 1.27-.20 (m, 9H), 1.12 (d, *J* = 6.8 Hz, 3H);

**$^{13}$C NMR** (126 MHz, $CD_3CN$) $\delta_C$ 170.7, 158.6, 157.7, 154.9, 152.5, 151.0, 145.2, 136.7, 136.1, 135.9, 130.0, 129.8, 128.0, 127.9, 127.7, 126.7, 122.2, 121.4, 118.7, 114.9, 112.9, 111.3, 85.7, 83.8, 60.0, 58.8, 58.7, 54.9, 43.1, 42.9, 40.5, 34.2, 28.8, 28.0, 27.6, 27.4, 26.6, 24.4, 24.0, 23.9, 23.94, 23.92, 23.9, 20.2, 20.1;

**$^{31}$P NMR** (202 MHz, $CD_3CN$) $\delta_P$ 148.4, 148.3;

**HRMS** (ESI/Q-TOF) $[M+H]^+$ calcd. for $C_{54}H_{65}N_9O_6P$ 966.4790, found 966.4910.

**Synthesis of (E)-N'-(8-(3-(1*H*-indol-3-yl)prop-1-yn-1-yl)-9-((2R,4S,5R)-4-hydroxy-5-(hydroxymethyl)tetrahydrofuran-2-yl)-9*H*-purin-6-yl)-N,N-dimethylformimidamide (29):**

To a solution of compound **25** (0.8 g, 1.98 mmol) in anhydrous DMF (10 mL) under argon atmosphere, dimethylformamide dimethylacetal (5.26 mL, 39.6 mmol) was added. The reaction mixture was stirred at room temperature for 4 h. After the reaction was complete, the solvent was removed under reduced pressure. The crude product was purified by flash chromatography using (0–10% MeOH in DCM) as the eluent. The desired product **29** (0.56 g, 1.22 mmol, 62%) was obtained as a yellow solid.

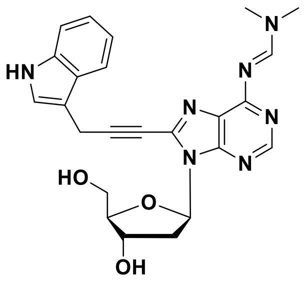


$\mathbf{R_f}$ (DCM/Methanol 9:1) 0.50;

**$^1$H NMR** (500 MHz, DMSO-$d_6$) $\delta_H$ 11.01 (s, 1H), 8.13 (s, 1H), 7.64 (d, *J* = 7.9 Hz, 1H), 7.50 (s, 2H), 7.40 (d, *J* = 8.1 Hz, 1H), 7.33 (d, *J* = 2.0 Hz, 1H), 7.13 (t, *J* = 7.4 Hz, 1H), 7.05 (t, *J* = 7.4 Hz, 1H), 6.48 (dd, *J* = 7.9, 6.7 Hz, 1H), 5.37 (dd, *J* = 7.9, 4.3 Hz, 1H), 5.33 (d, *J* = 4.0 Hz, 1H), 4.50 – 4.46 (m, 1H), 4.10 (s, 2H), 3.91 (dd, *J* = 6.7, 4.3 Hz, 1H), 3.67 (dt, *J* = 11.8, 4.3 Hz, 1H), 3.53 – 3.47 (m, 1H), 3.15 (ddd, *J* = 13.6, 8.8, 5.6 Hz, 2H), 2.18 (ddd, *J* = 12.8, 6.2, 2.2 Hz, 1H);

**$^{13}$C NMR** (126 MHz, DMSO-$d_6$) $\delta_C$ 159.6, 158.4, 152.7, 151.0, 136.9, 135.8, 126.80, 126.1, 123.7, 121.9, 119.2, 118.8, 112.1, 108.4, 97.3, 88.7, 85.5, 71.7, 70.8, 62.7, 41.2, 37.9, 35.1;

**HRMS** (ESI/Q-TOF) $[M+H]^+$ calcd. for $C_{24}H_{26}N_7O_3$ 460.2092, found 460.2088.

**Synthesis of (E)-N'-(8-(3-(1*H*-indol-3-yl)prop-1-yn-1-yl)-9-((2R,4S,5R)-5-((bis(4-methoxyphenyl)(phenyl)methoxy)methyl)-4-hydroxytetrahydrofuran-2-yl)-9*H*-purin-6-yl)-N,N-dimethylformimidamide (30):**

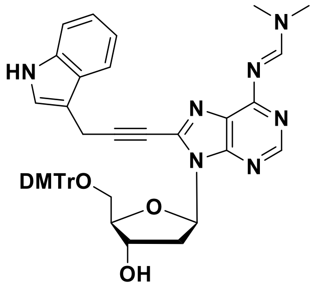


To a solution of compound **29** (0.45 g, 0.98 mmol) in anhydrous pyridine (12 mL), DMAP (12 mg, 0.1 mmol) was added. To this mixture, a solution of 4,4'-dimethoxytrityl chloride (0.4 g, 1.17 mmol) in 8 mL of anhydrous pyridine was added in four equal portions over the course of 1 h. The reaction mixture was stirred at room temperature for 12 h. After completion of the reaction, the solvent was evaporated under reduced pressure. The resulting crude mixture was purified by flash chromatography (0–2% MeOH in $CH_2Cl_2$ with 0.2% $Et_3N$) yielding the desired compound **30** (0.4 g, 0.53 mmol, 54%) as a pale-yellow solid.

$\mathbf{R_f}$ (DCM/Methanol 9:1) 0.6;

**$^1$H NMR** (500 MHz, DMSO-$d_6$) $\delta_H$ 11.02 (s, 1H), 8.89 (s, 1H), 8.26 (s, 1H), 7.65 (d, *J* = 7.9 Hz, 1H), 7.40 (d, *J* = 8.1 Hz, 1H), 7.34 (s, 1H), 7.29 (d, *J* = 7.2 Hz, 2H), 7.23 – 7.09 (m, 8H), 7.04 (t, *J* = 7.4 Hz, 1H), 6.77 (d, *J* = 8.7 Hz, 2H), 6.71 (d, *J* = 8.8 Hz, 2H), 6.53 (t, *J* = 6.5 Hz, 1H), 5.39 (br.s, 1H), 4.63 (br.s, 1H), 4.06 (s, 2H), 4.01 (dd, *J* = 9.3, 4.7 Hz, 1H), 3.70 (s, 3H), 3.68 (s, 3H), 3.21 (s, 3H), 3.17 (m, 3H), 3.13 (s, 3H), 2.30 – 2.22 (m, 2H);

**$^{13}$C NMR** (126 MHz, DMSO-$d_6$) $\delta_C$ 159.4, 158.4, 158.3, 158.2, 152.8, 151.0, 145.5, 136.9, 136.2, 136.1, 136.1, 130.2, 130.0, 128.1, 128.1, 127.0, 126.8, 126.0, 123.7, 121.9, 119.2, 118.8, 113.5, 113.4, 112.1, 108.2, 96.9, 86.3, 85.7, 84.9, 71.3, 71.2, 55.4, 55.4, 41.2, 37.2, 35.1, 16.1;

**HRMS** (ESI/Q-TOF) $[M+H]^+$ calcd. for $C_{45}H_{44}N_7O_5$ 762.3404, found 762.3409.

**Synthesis of (2R,3S,5R)-5-(8-(3-(1*H*-indol-3-yl)prop-1-yn-1-yl)-6-(((E)-(dimethylamino)methylene)amino)-9*H*-purin-9-yl)-2-((bis(4-methoxyphenyl)(phenyl)methoxy)methyl)tetrahydrofuran-3-yl (2-cyanoethyl) diisopropylphosphoramidite (4b):**

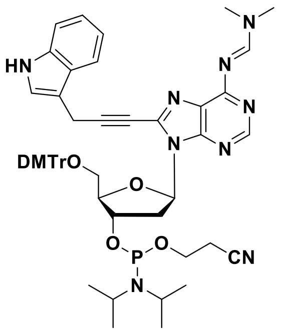


To a solution of compound **30** (0.3 g, 0.39 mmol) in anhydrous $CH_2Cl_2$, *N,N*-diisopropylethylamine (0.34 mL, 1.97 mmol) was added under stirring. The reaction mixture was cooled to 0 °C, and 2-cyanoethyl-*N,N*-diisopropylchlorophosphoramidite (0.13 mL, 0.63 mmol) was added under an argon atmosphere. The mixture was allowed to warm to room temperature and stirred for 1.5 h. The reaction mixture was then diluted with anhydrous $CH_2Cl_2$ (20 mL) and washed successively with 5% $NaHCO_3$ (20 mL) and brine solution (20 mL). The organic phase was dried over anhydrous $Na_2SO_4$ and concentrated under reduced pressure. The crude product was purified by column chromatography using (20–30% acetone in hexane containing 0.5% $Et_3N$). The desired compound **4b** (0.23 g, 0.24 mmol, 61%) was obtained as a white foam.

TLC (AcOEt:Hexane 9:1) $R_f$ =0.35;

**$^1$H NMR** (500 MHz, $CD_3CN$) $\delta_H$ 9.29 (br. s, 1H), 8.90 (s, 1H), 8.31 (s, 1H), 7.73 (d, *J* = 7.8 Hz, 1H), 7.49 (d, *J* = 8.8 Hz, 1H), 7.40 – 7.33 (m, 3H), 7.27 – 7.18 (m, 7H), 7.15 (t, *J* = 7.9 Hz, 1H), 6.81 – 6.70 (m, 4H), 6.63 – 6.56 (m, 1H), 5.17 – 5.08 (m, 1H), 4.19 (m, 1H), 4.07 (m, 2H), 3.77 (s, 3H), 3.76 (s, 3H), 3.73 – 3.67 (m, 2H), 3.67 – 3.59 (m, 2H), 3.57 – 3.47 (m, 2H), 3.36 (dd, *J* = 10.3, 3.7 Hz, 1H), 3.26 (m, 1H), 3.21 (s, 3H), 3.19 (s, 3H), 2.80 (t, *J* = 5.9 Hz, 1H), 2.55 (t, *J* = 6.0 Hz, 1H), 2.53 – 2.42 (m, 1H), 1.28 (d, *J* = 6.7 Hz, 3H), 1.24 (d, *J* = 7.1 Hz, 3H), 1.21 (m, 6H);

**$^{13}$C NMR** (126 MHz, DMSO-*d$_6$*) $\delta_C$ 159.4, 158.55, 158.5, 158.0, 152.6, 150.9, 145.2, 136.7, 136.3, 136.1, 135.9, 130.0, 128.0, 127.7, 126.7, 126.0, 123.0, 121.9, 119.2, 118.4, 112.9, 111.6, 108.7, 96.1, 85.8, 85.2, 84.6, 73.0, 72.9, 71.0, 63.6, 60.0, 58.8, 58.6, 54.9, 54.9 45.1, 43.1, 43.1, 43.0, 40.6, 36.2, 34.2, 24.0, 23.9, 23.9, 23.9, 22.2, 22.1, 20.2, 15.9, 13.6;

**$^{31}$P NMR** (202 MHz, $CD_3CN$) $\delta_P$ 148.0, 147.8 ;

**HRMS** (ESI/Q-TOF) $[M+H]^+$ calcd. for $C_{54}H_{61}N_9O_6P$ 982.4739, found 982.4736.

### 2. List of base-modified DNA sequences, along with their calculated and measured masses.

| Strands | Sequences | Mass calc. [Da] | Mass found [Da] |
|---|---|---|---|
| Z1<br>Z11 | 5' - F - TTG*AATT***CCCGGG**TCCAAA - 3'<br>5' - F - TTG*AATT***CCCGGG**TCCAAA - 3' | 6512.5<br>6508.5 | 6512.5<br>6508.4 |
| Z4<br>Z44 | 5' - F - TTG*AAUU***CCCGGG**TCCAAA - 3'<br>5' - F - TTG*AAUU***CCCGGG**TCCAAA - 3' | 7113.3<br>7093.2 | 7113.5<br>7093.3 |
| Z5<br>Z55 | 5' - F - TTG*AATU***CCCGGG**TCCAAA - 3'<br>5' - F - TTG*AATU***CCCGGG**TCCAAA - 3' | 6498.5<br>6494.5 | 6498.4<br>6492.3 |
| Z6 | 5' - F - TTG*AAUU***CCCGGG**TCCAAA - 3' | 6641.7 | 6642.0 |
| cZ3 | 3' - Q - TTC*UUAA***GGGCCC**AG - 5' | 5474.1 | 5473.7 |
| cZ4 | 3' - Q - TTC*UUAA***GGGCCC**AG - 5' | 5631.3 | 5630.8 |
| cZ5 | 3' - Q - TTC*UUAA***GGGCCC**AG - 5' | 5788.5 | 5793.3 |

**Table S1**: (•) solid circles: 3-propyl-indole linked nucleotide; (○) open circles: 3-(prop-2-yn-1-yl)-indole linked nucleotide; F: 5'-(6-FAM)-labeled; Q: 3'-(Dabcyl)-labeled. *Italics* EcoR1 restriction site; **bold** XmaI, SmaI restriction site.

**A Melting points**

| | WT | Z1 | Z5 | Z6 | Z4 |
|---|---|---|---|---|---|
| cWT | 51°C | 44°C | 43°C | 39°C | 33°C |
| cZ3 | 44°C | 41°C | 37°C | 35°C | 38°C |
| cZ4 | 39°C | 37°C | 36°C | 36°C | 39°C |
| cZ5 | 33°C | 31°C | 30°C | 32°C | 43°C |

| | Z11 | Z55 | Z44 |
|---|---|---|---|
| cWT | 42°C | 34°C | 33°C |

**B Structural analysis using CD**

| | WT | Z1 | Z5 | Z6 | Z4 |
|---|---|---|---|---|---|
| cWT | B | BZ | B | B | Distorted B |
| cZ3 | Distorted B | Distorted B | BZ | B | Distorted B |
| cZ4 | Distorted B | Distorted B | BZ | Z | Distorted B |
| cZ5 | BZ | Distorted B | Z | BZ | B |

| | Z11 | Z55 | Z44 |
|---|---|---|---|
| cWT | B | B | B |

**Figure S1:** Melting points and structures of Zimera duplexes. (A) Grid showing the melting points of Zimera duplexes compared to wildtype. Melting point was monitored using FRET assays with a Fluorophore-labeled sense strand (WT; Z1-Z55) and a quencher- labeled antisense strand (cWT; cZ3-cZ5). (B) Grid showing structural variation of Zimera duplexes compared to wildtype. Analysis of the structure was performed by CD spectroscopy using 1: 2.5 ratio of WT/Z1-Z55: cWT/Z3/Z4/Z5 in PBS, 20°C.

### 3. Thermal studies of ZImera

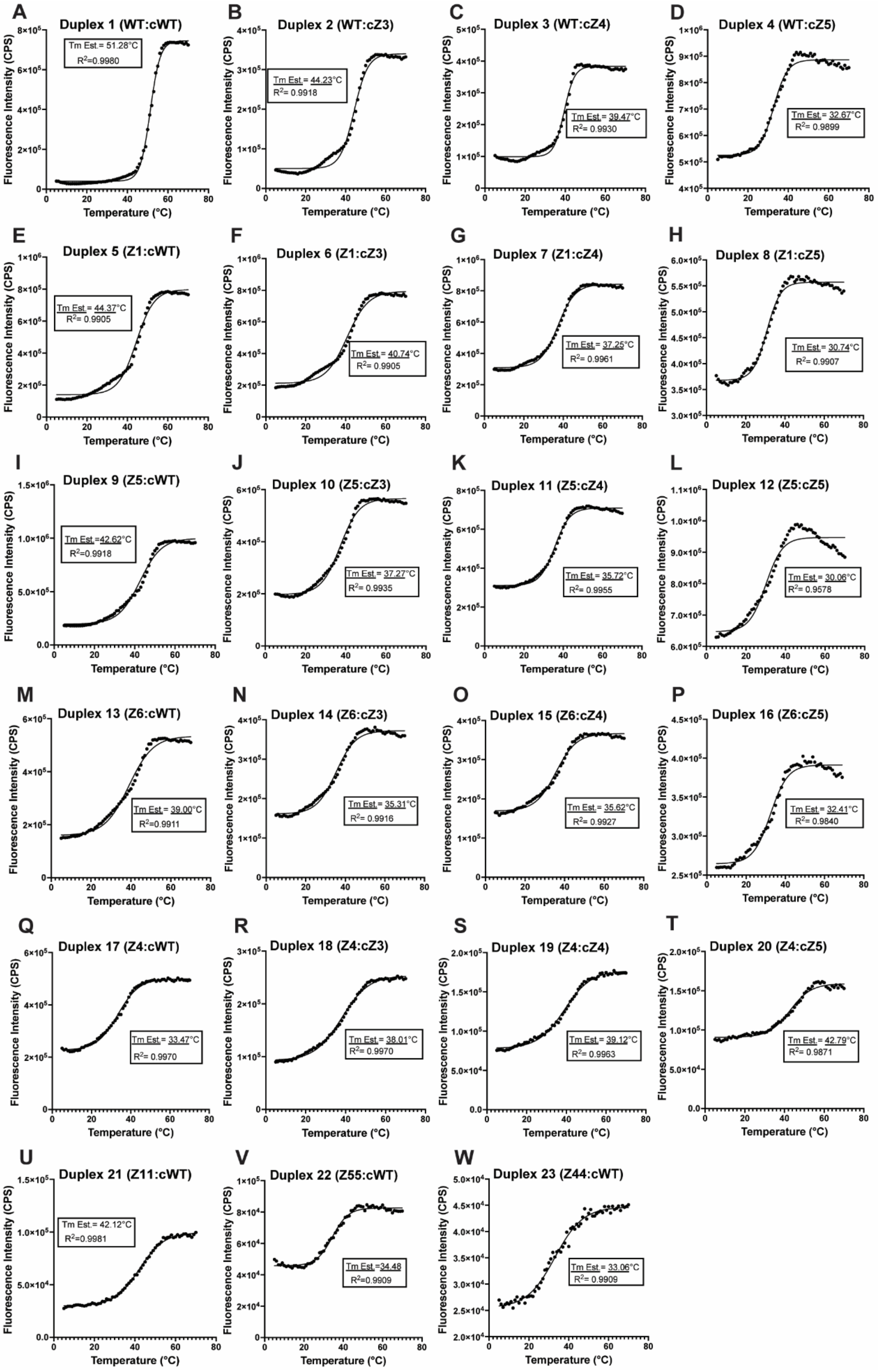


**Figure S2**. Thermal analysis of ZImera was performed by recording Tm measurements in a quartz cuvette with a 3 mm pathlength. The temperature was varied in 1°C increments across a range from 5 °C to 70 °C.

### 4. FRET analysis

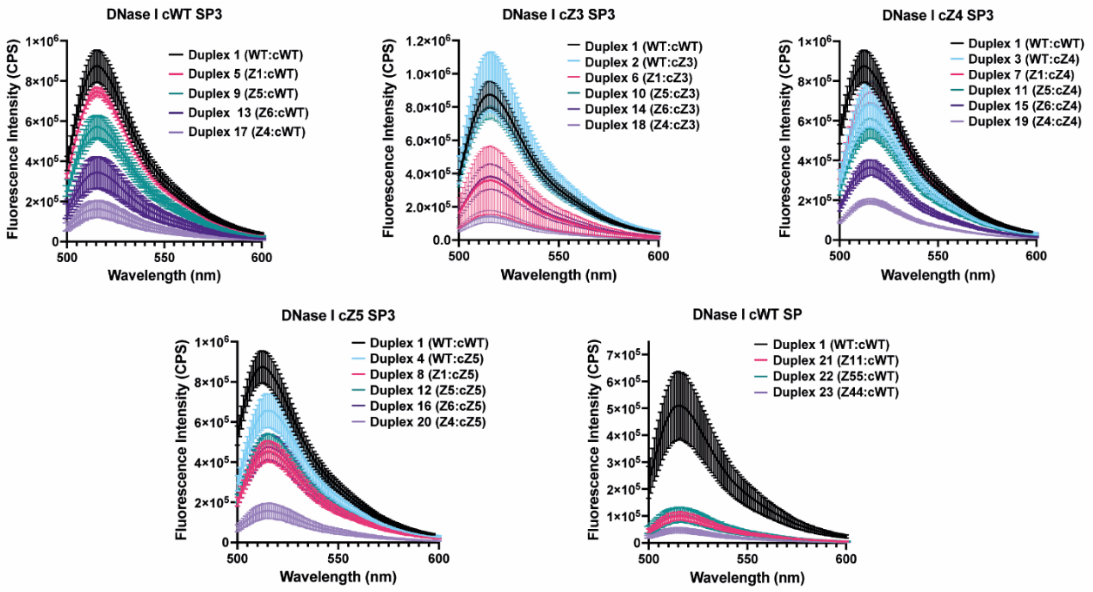


**Figure S3**. Analysis of DNase 1 activity of ZImera. The DNase I activity of ZImera were analyzed by preparing 10 µmol DNA samples in a 10 µL volume. The samples were treated with 1.4 units of DNase I (Thermo Scientific, REF# EN0521, 0.56 u/µL) for 15 minutes at 37 °C. Fluorescent measurements were then conducted at 20 °C, within the wavelength range of 500 nm to 600 nm.

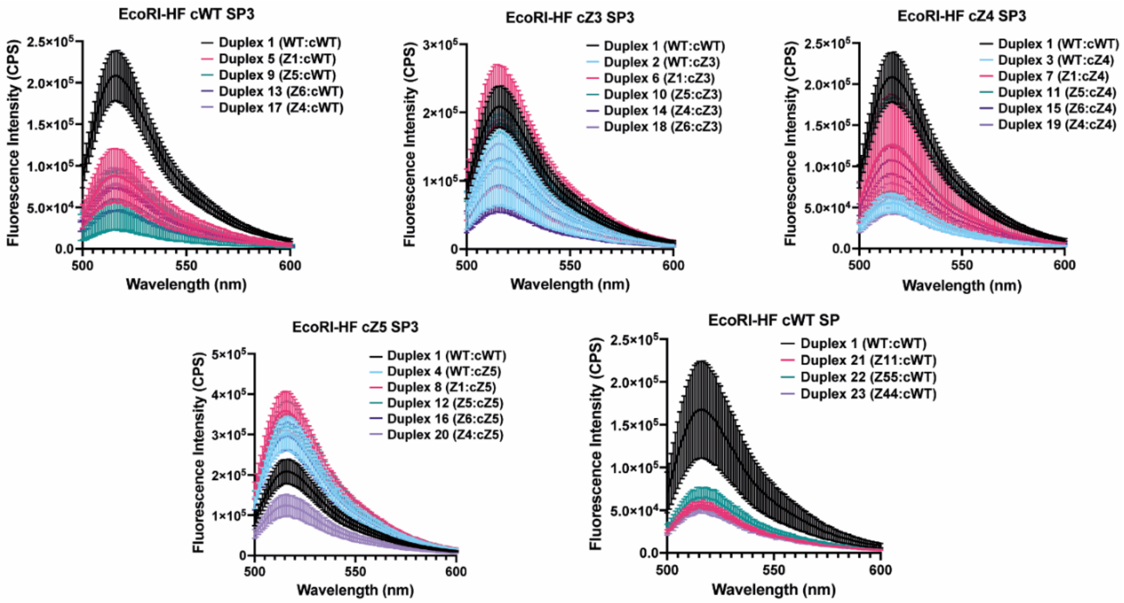


**Figure S4**. Analysis of EcoRI activity of ZImera. The EcoRI activity of ZImera were analyzed by preparing 10 µmol DNA samples in a 10 µL volume. The samples were treated with 70 units of EcoRI-HF (NEB,

R3101M, 28 u/µL) for overnight at 37 °C. Fluorescent measurements were then conducted at 20 °C, within the wavelength range of 500 nm to 600 nm.

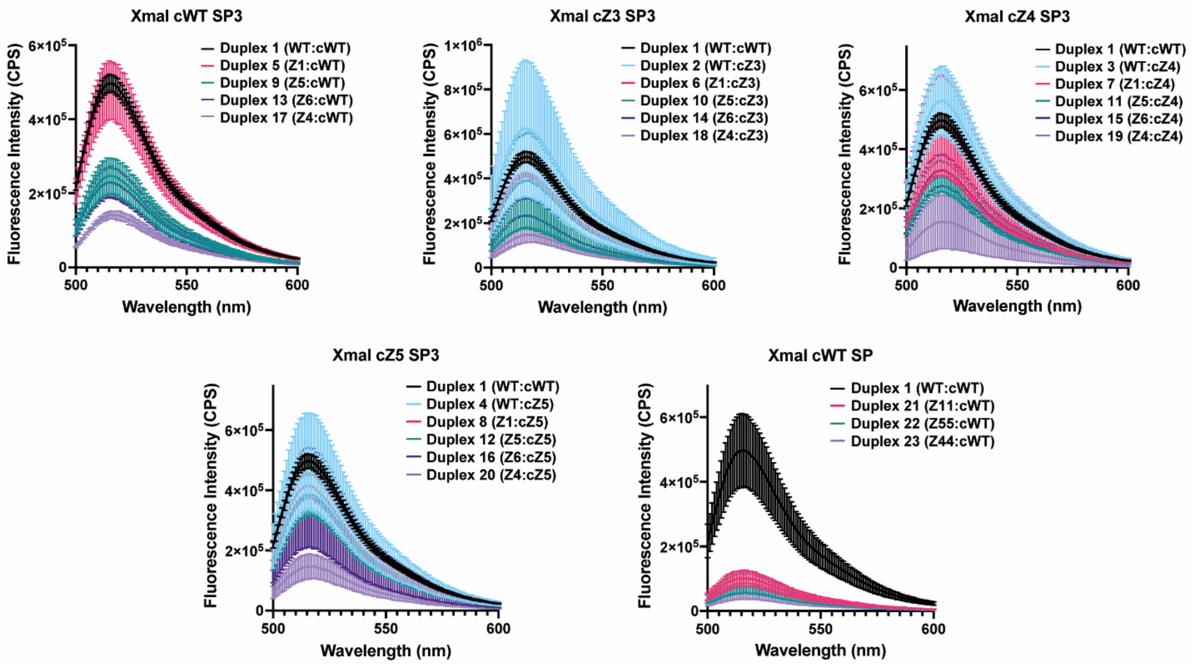


**Figure S5**. Analysis of XmaI activity of ZImera. The XmaI activity of ZImera were analyzed by preparing 10 µmol DNA samples in a 10 µL volume. The samples were treated with 7 units of XmaI (NEB, R0180S, 2.8 u/µL) for overnight at 37 °C. Fluorescent measurements were then conducted at 20 °C, within the wavelength range of 500 nm to 600 nm.

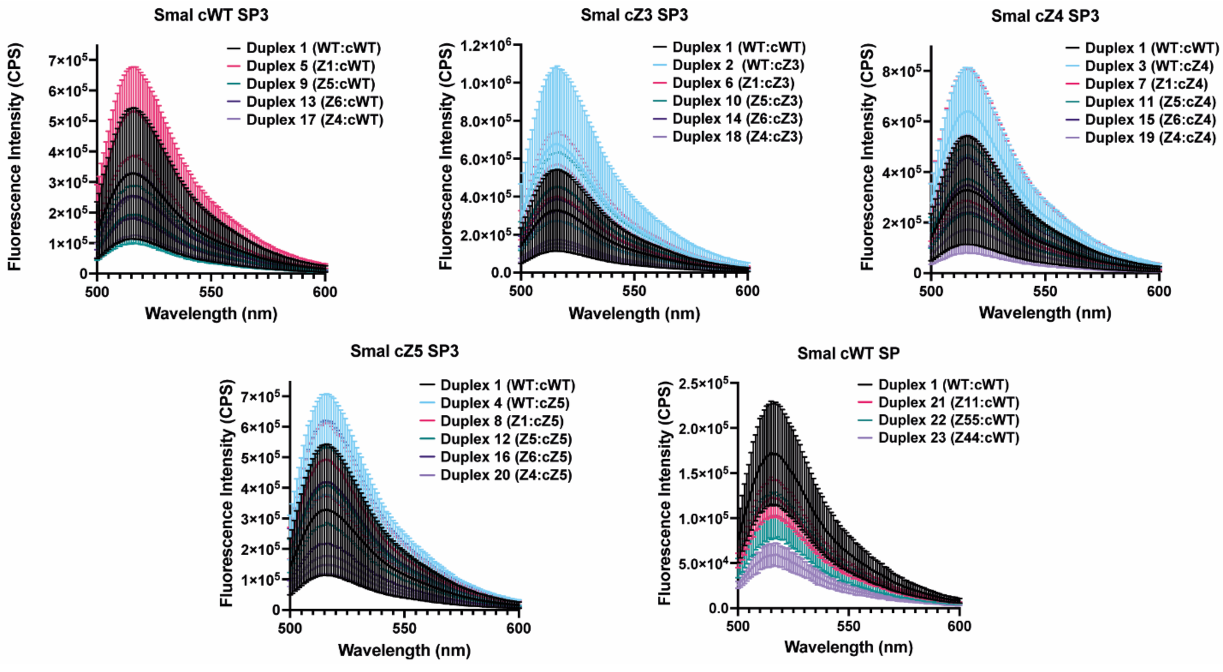


**Figure S6**. Analysis of SmaI activity of ZImera. The SmaI activity of ZImera were analyzed by preparing 10 µmol DNA samples in a 10 µL volume. The samples were treated with 7 units of SmaI

(Thermoscientific, REF# ER0665, 2.8 u/μL) for overnight at 25 °C. Fluorescent measurements were then conducted at 20 °C, within the wavelength range of 500 nm to 600 nm.

### 5. NMR Spectra:

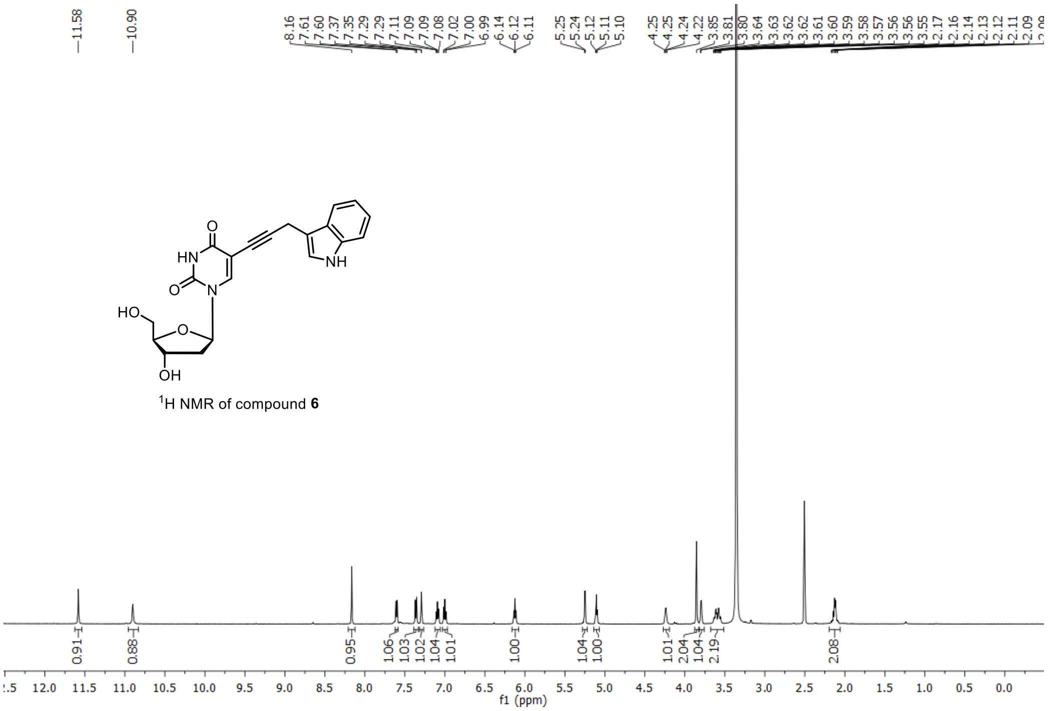

$^1$H NMR of compound **6**

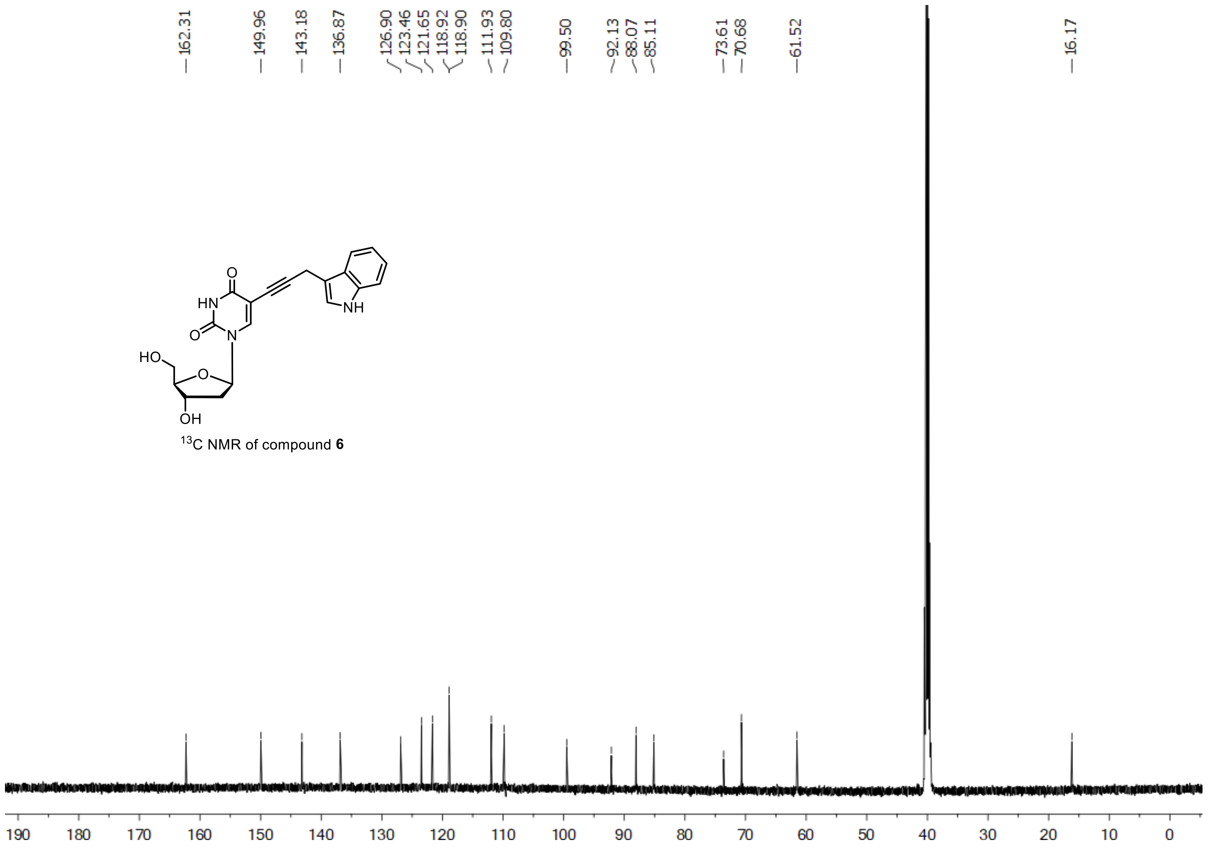

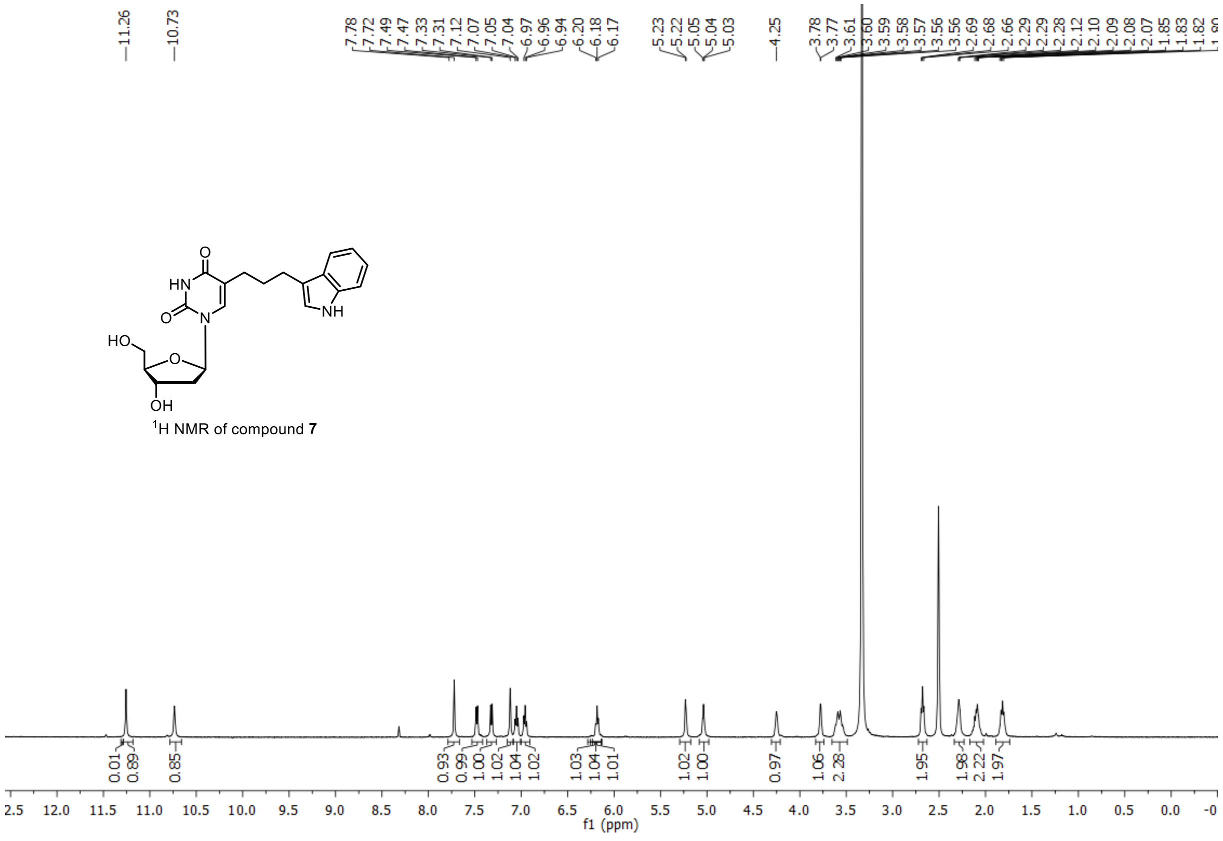

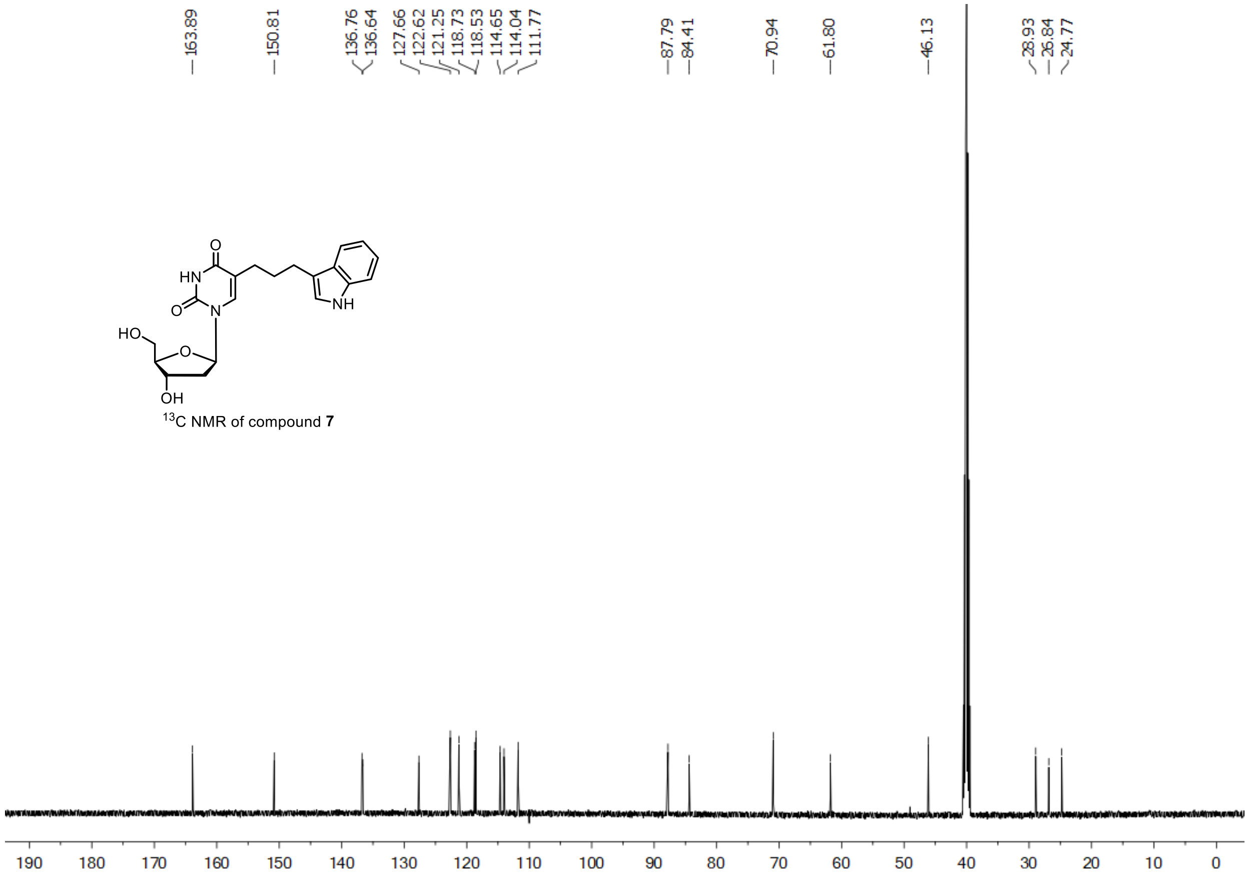


$^{13}$C NMR of compound **7**

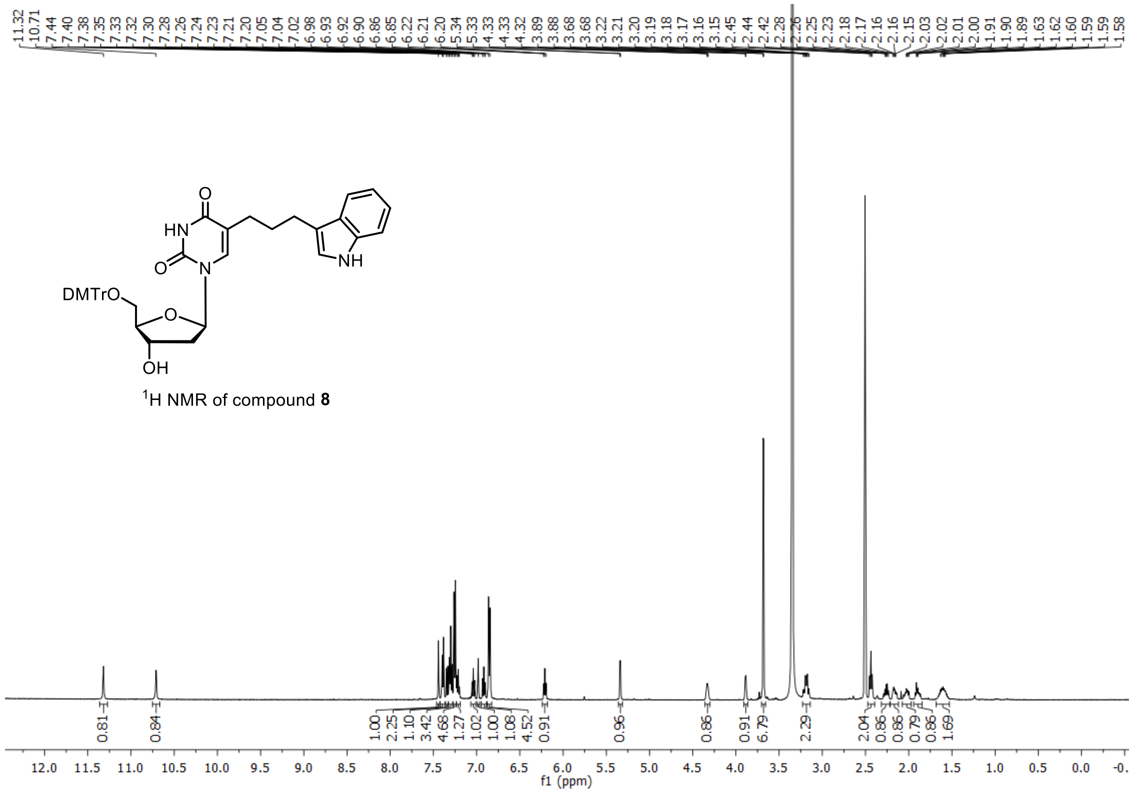


$^{1}$H NMR of compound **8**

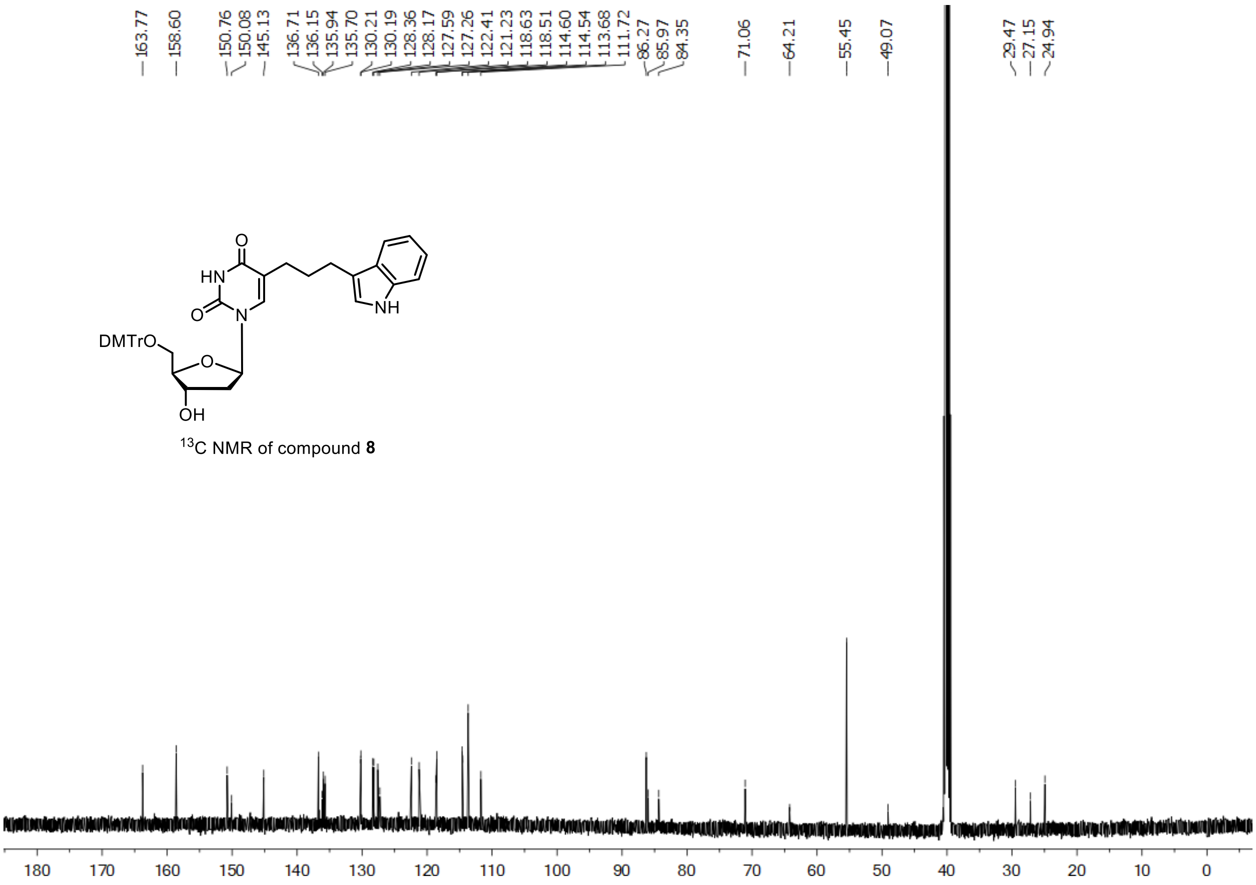


$^{13}$C NMR of compound **8**

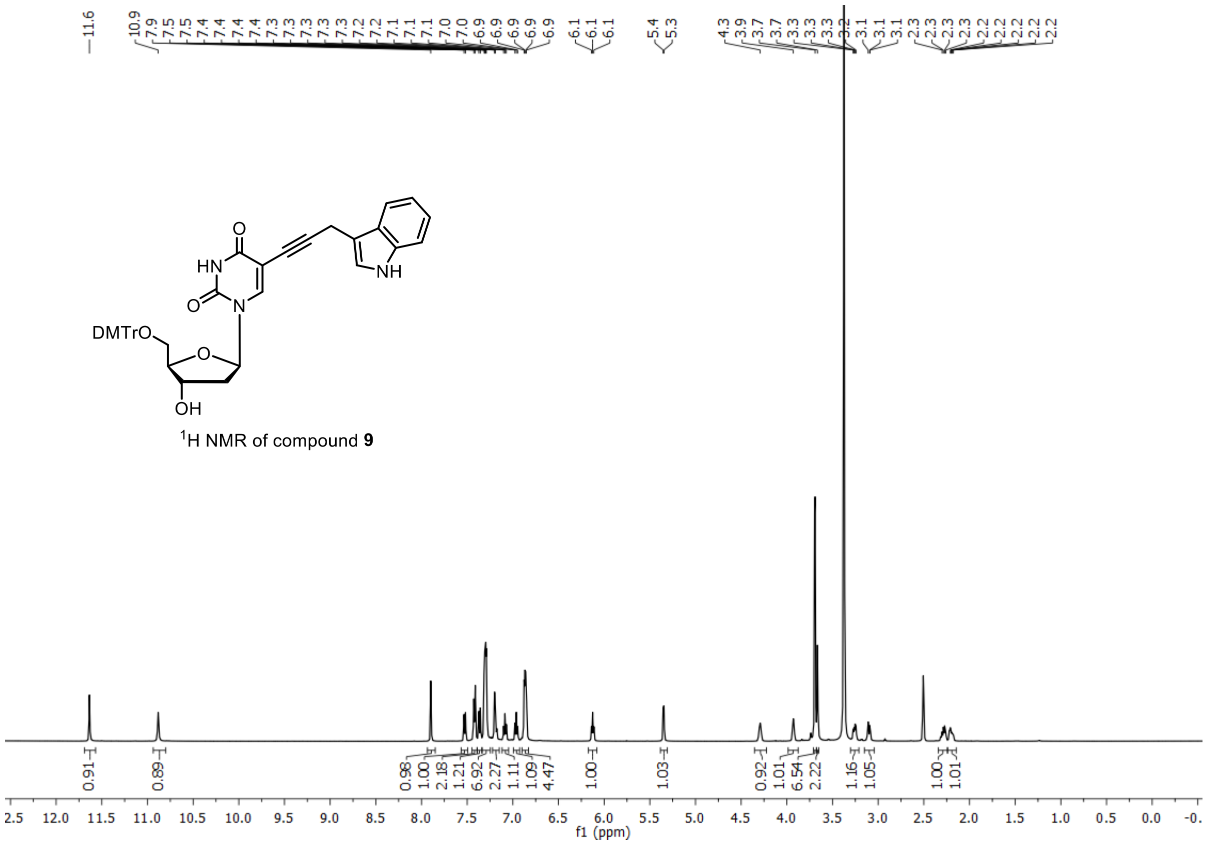


[1]H NMR of compound **9**

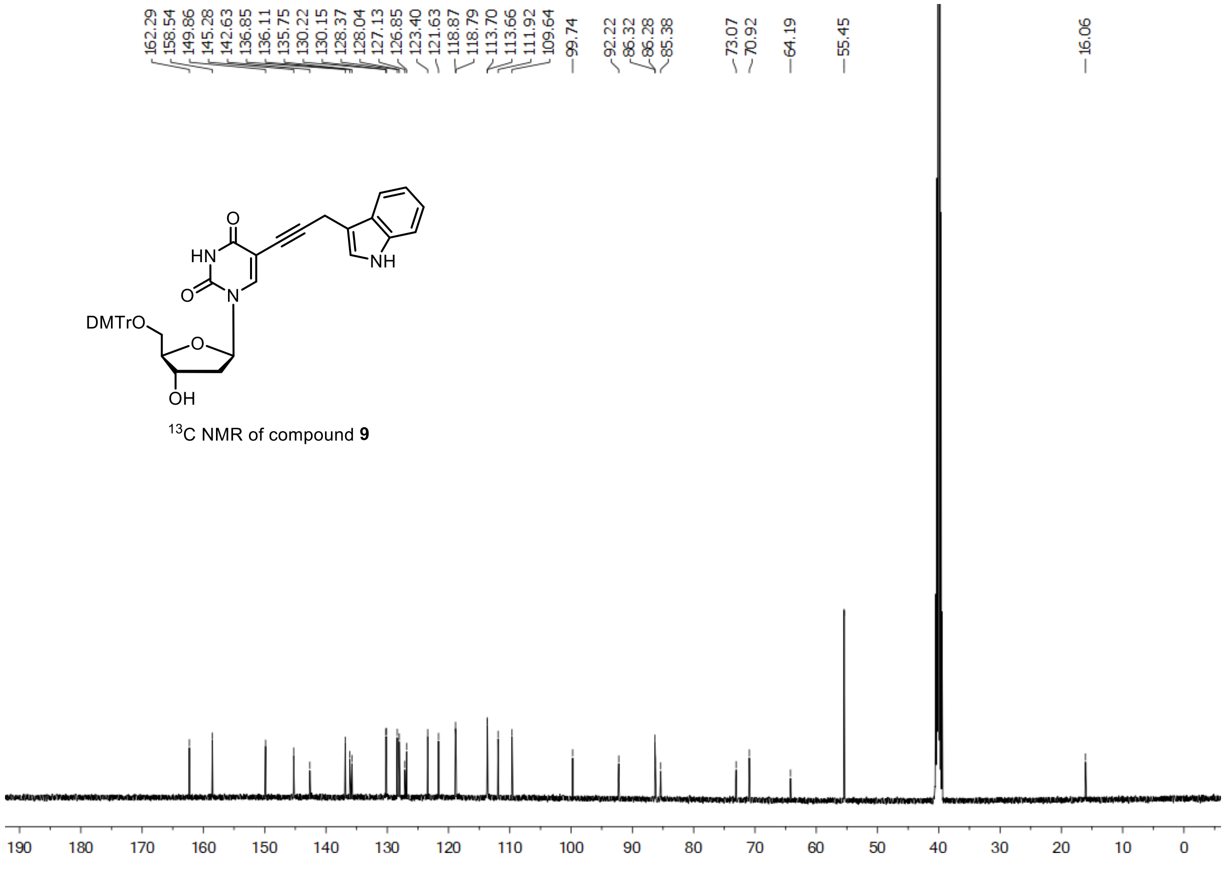


[13]C NMR of compound **9**

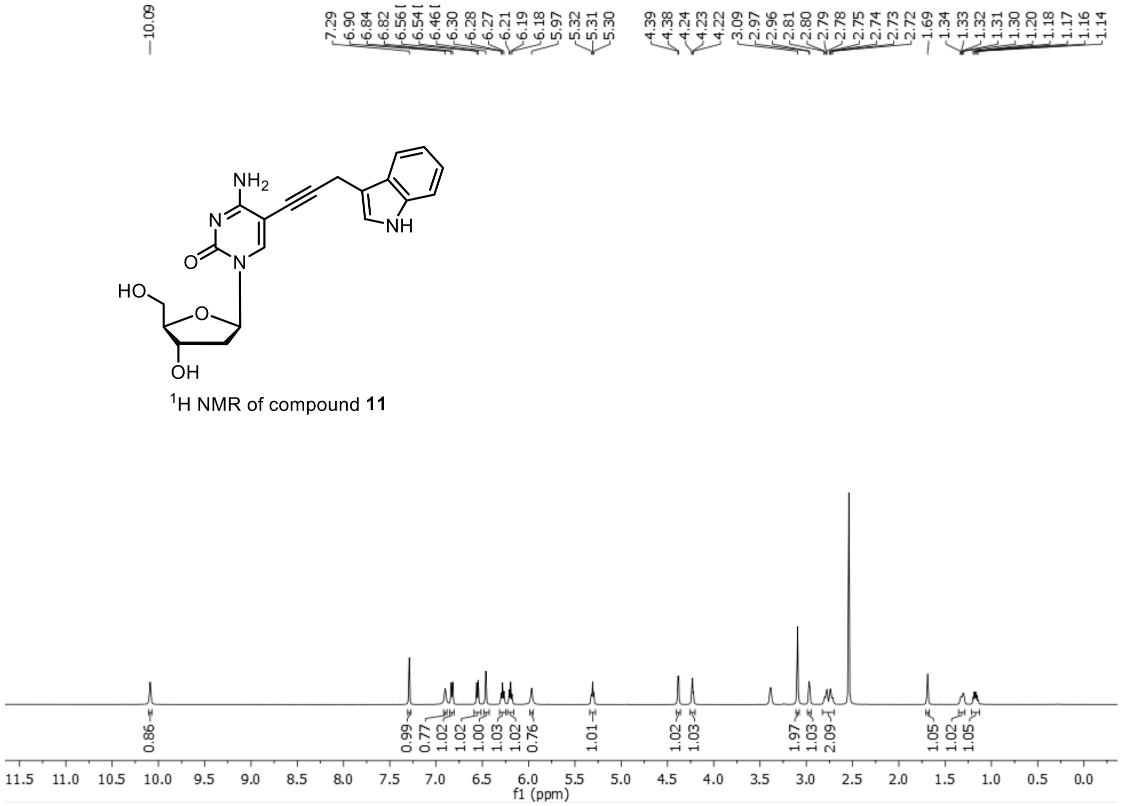


$^1$H NMR of compound **11**

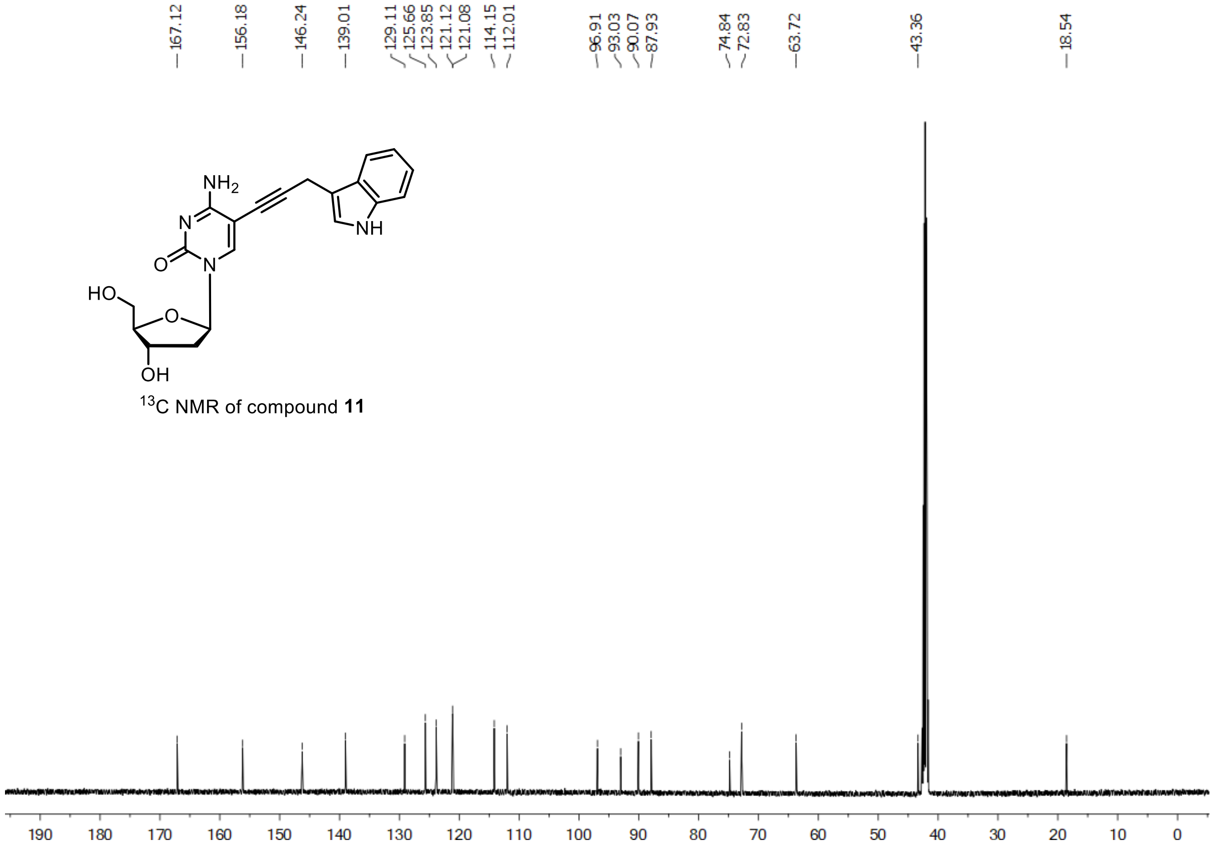


$^{13}$C NMR of compound **11**

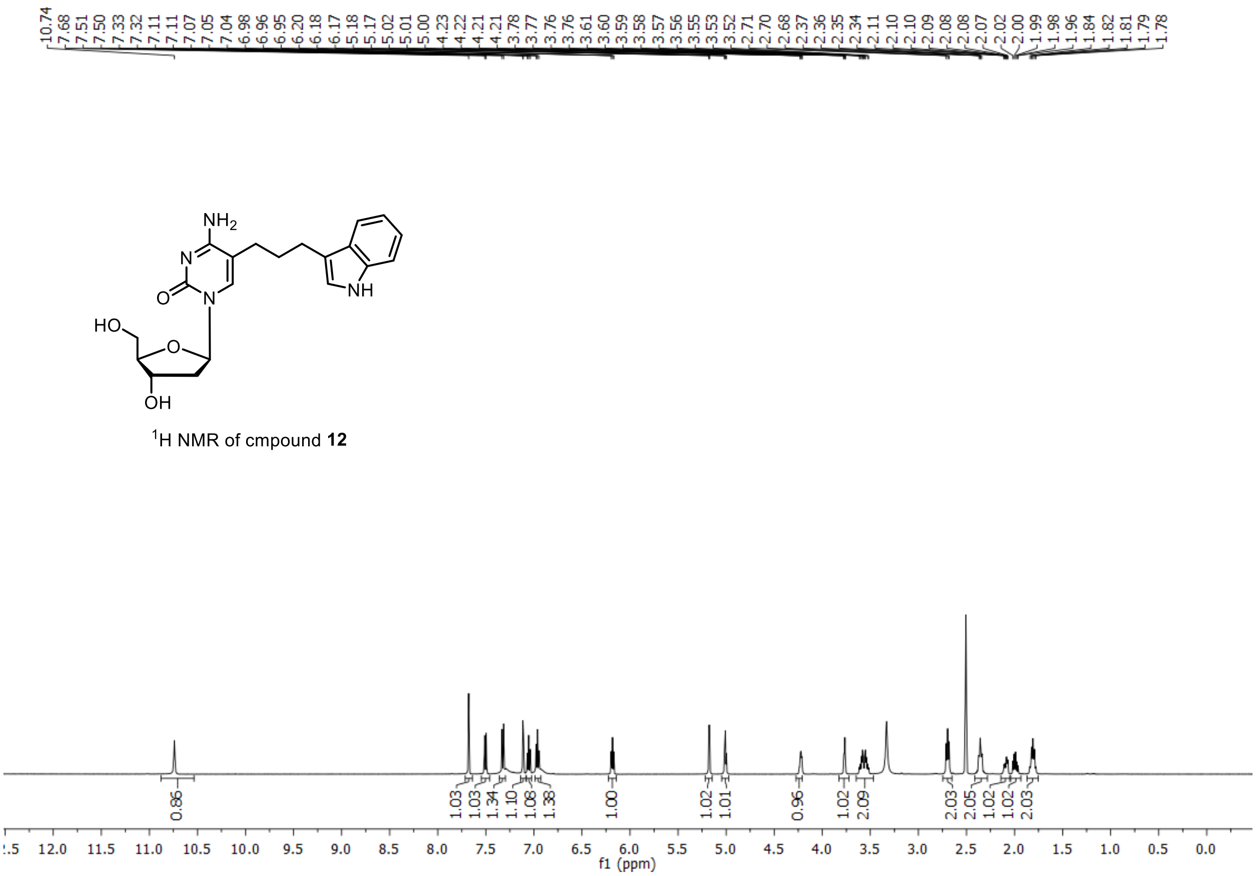


$^1$H NMR of cmpound **12**

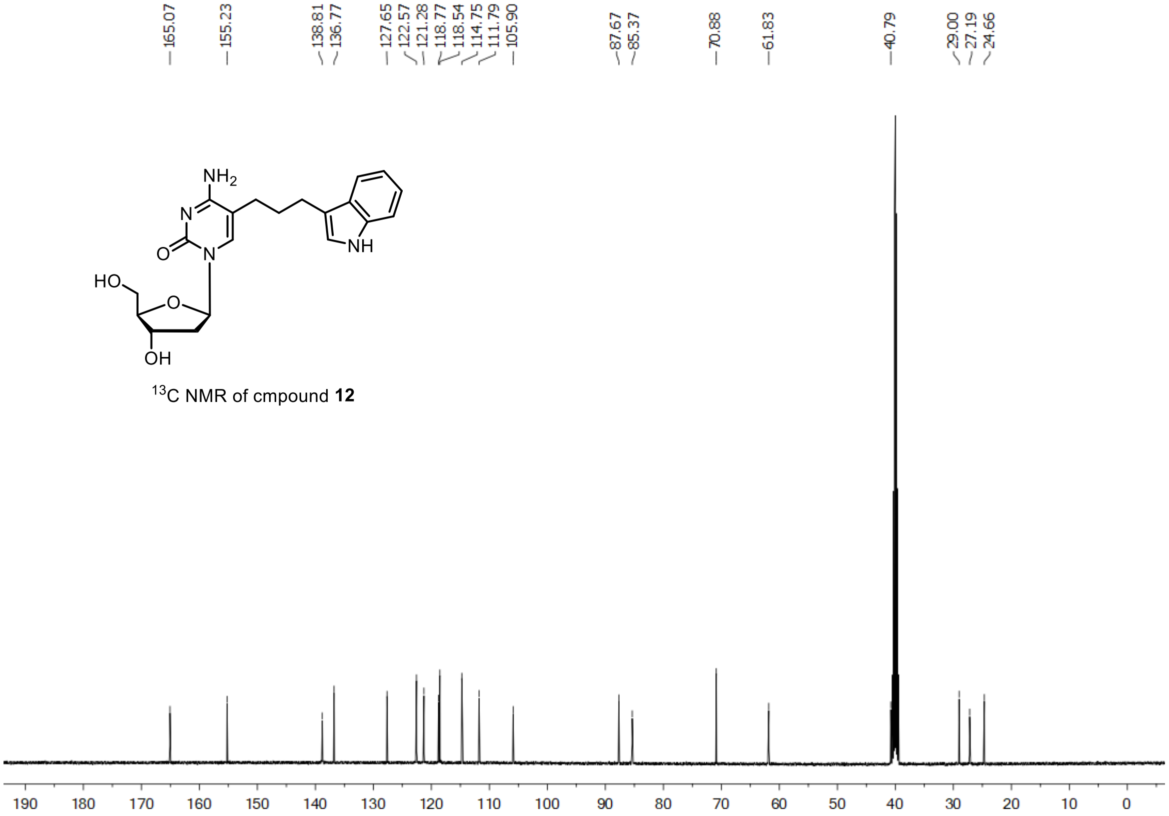


$^{13}$C NMR of cmpound **12**

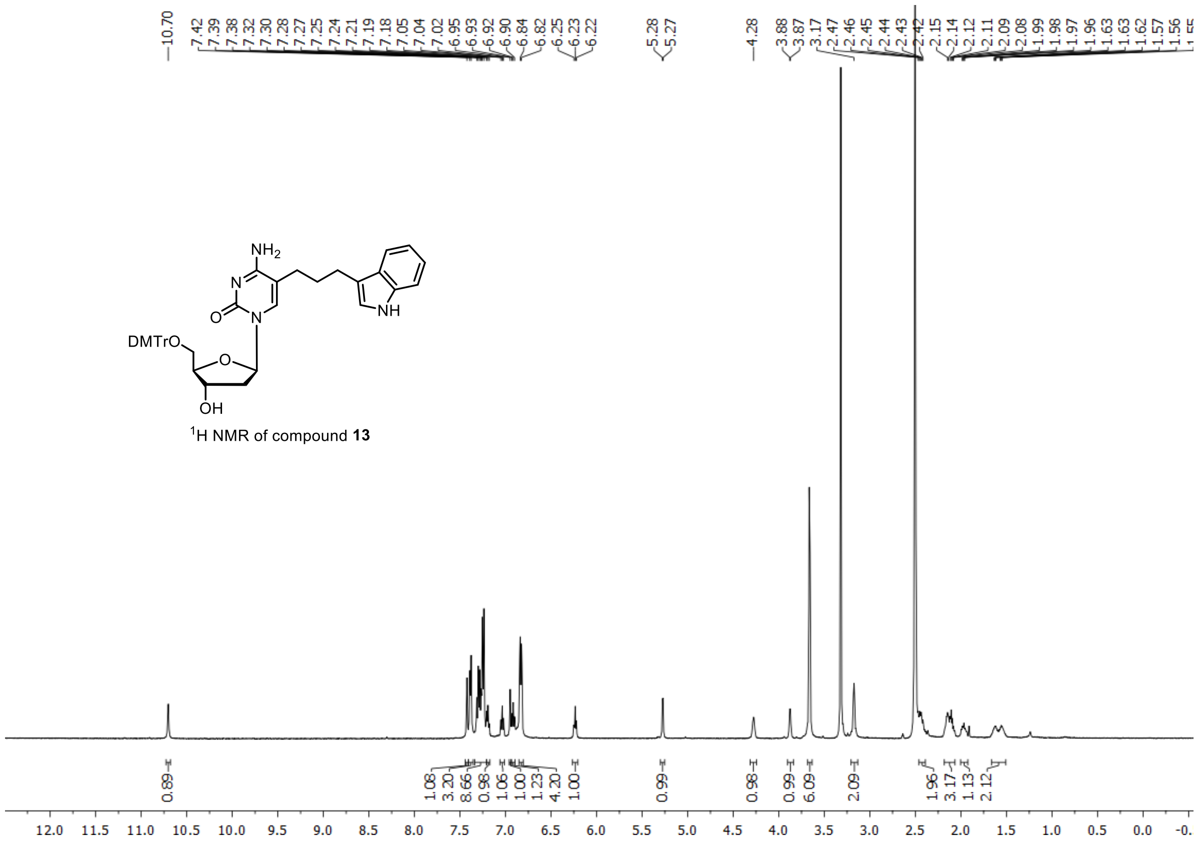


$^1$H NMR of compound **13**

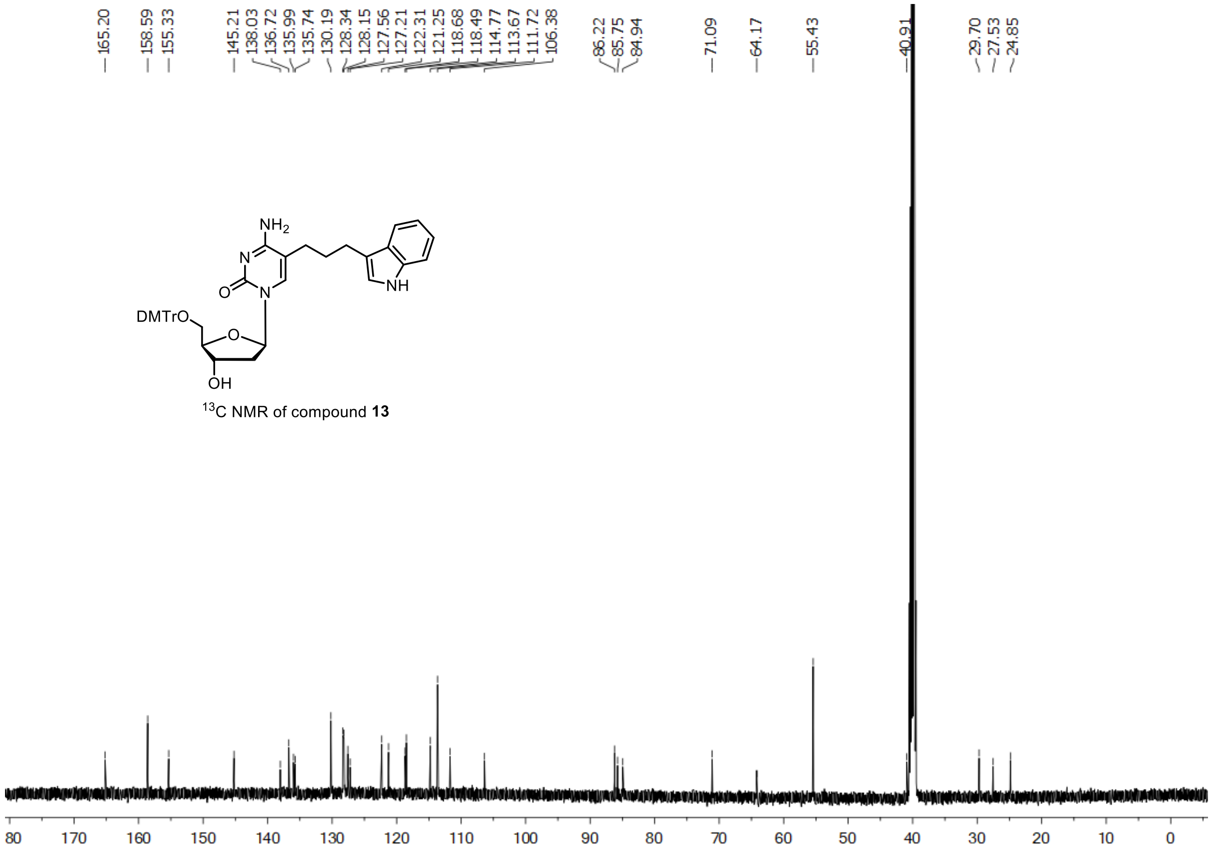


$^{13}$C NMR of compound **13**

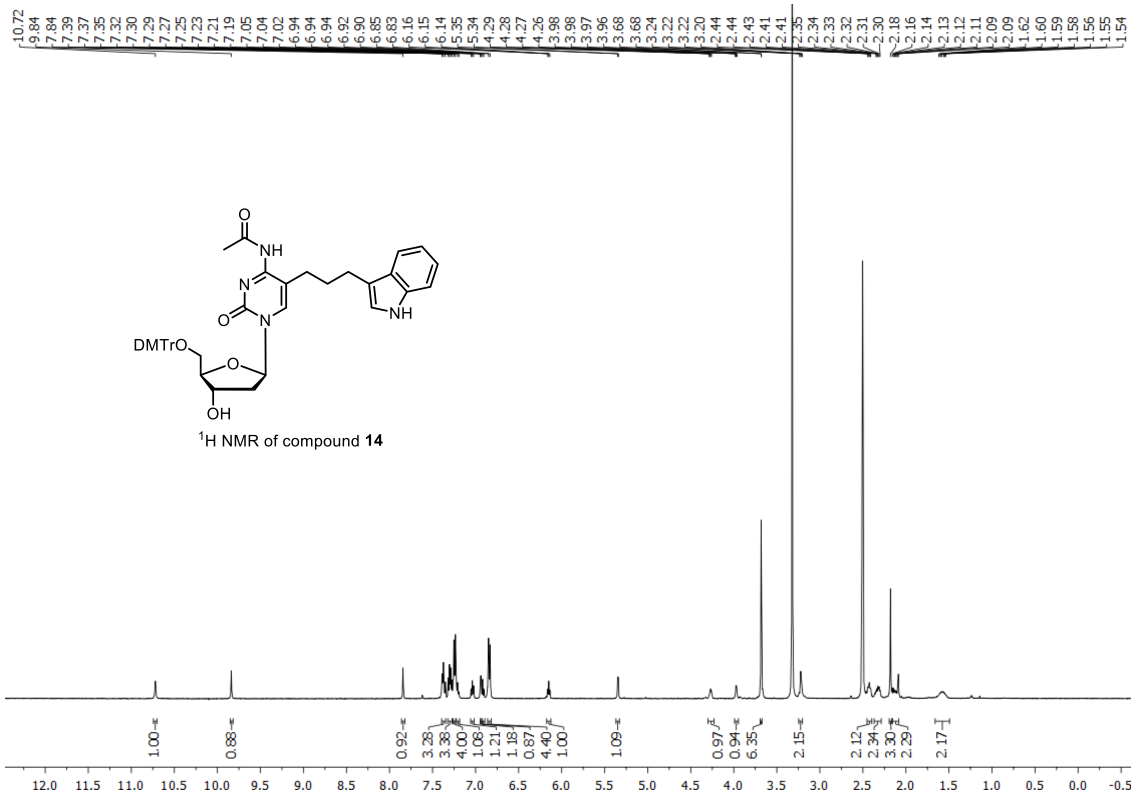

[1]H NMR of compound **14**

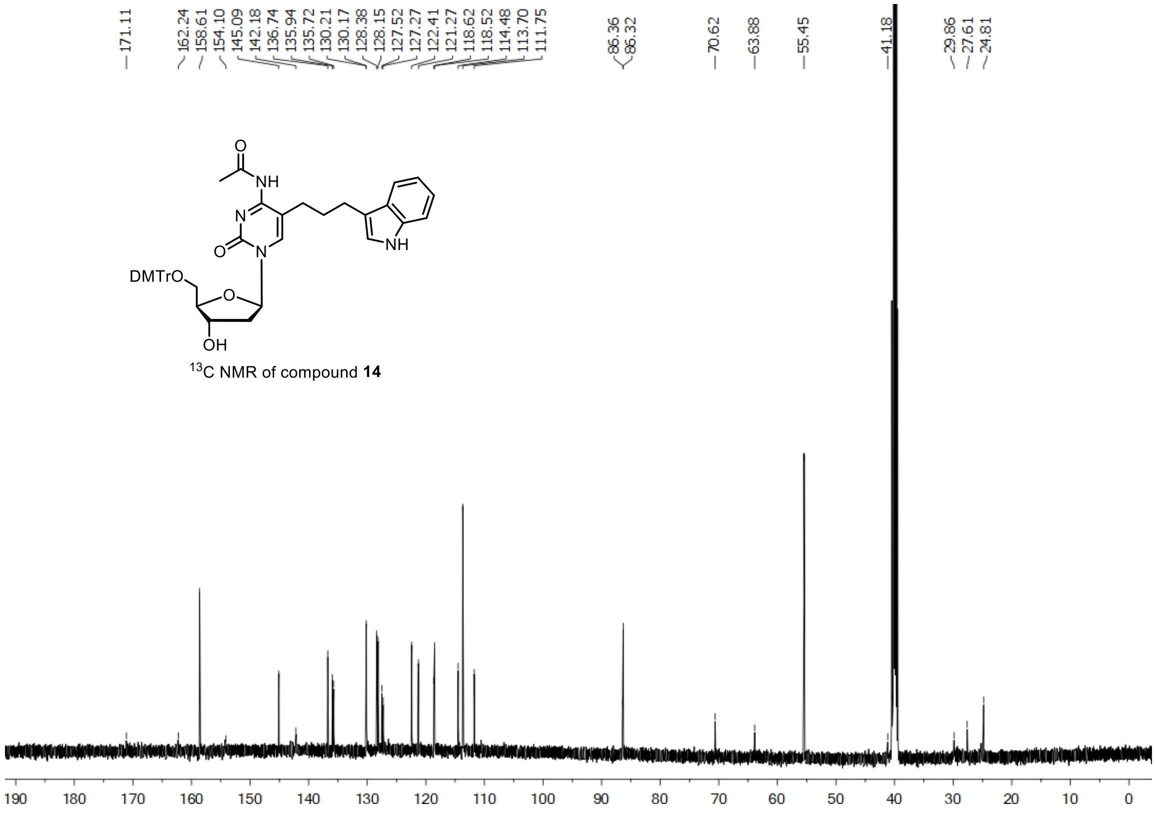

[13]C NMR of compound **14**

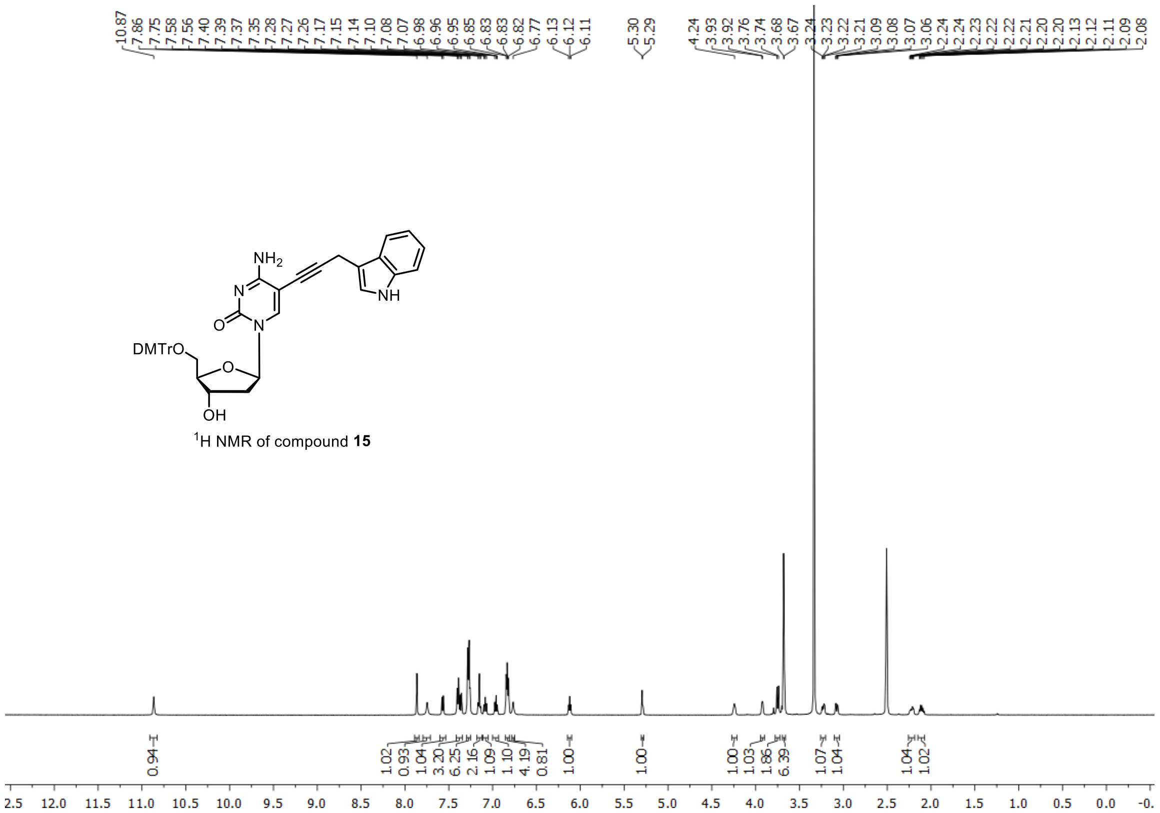


$^{1}H$ NMR of compound **15**

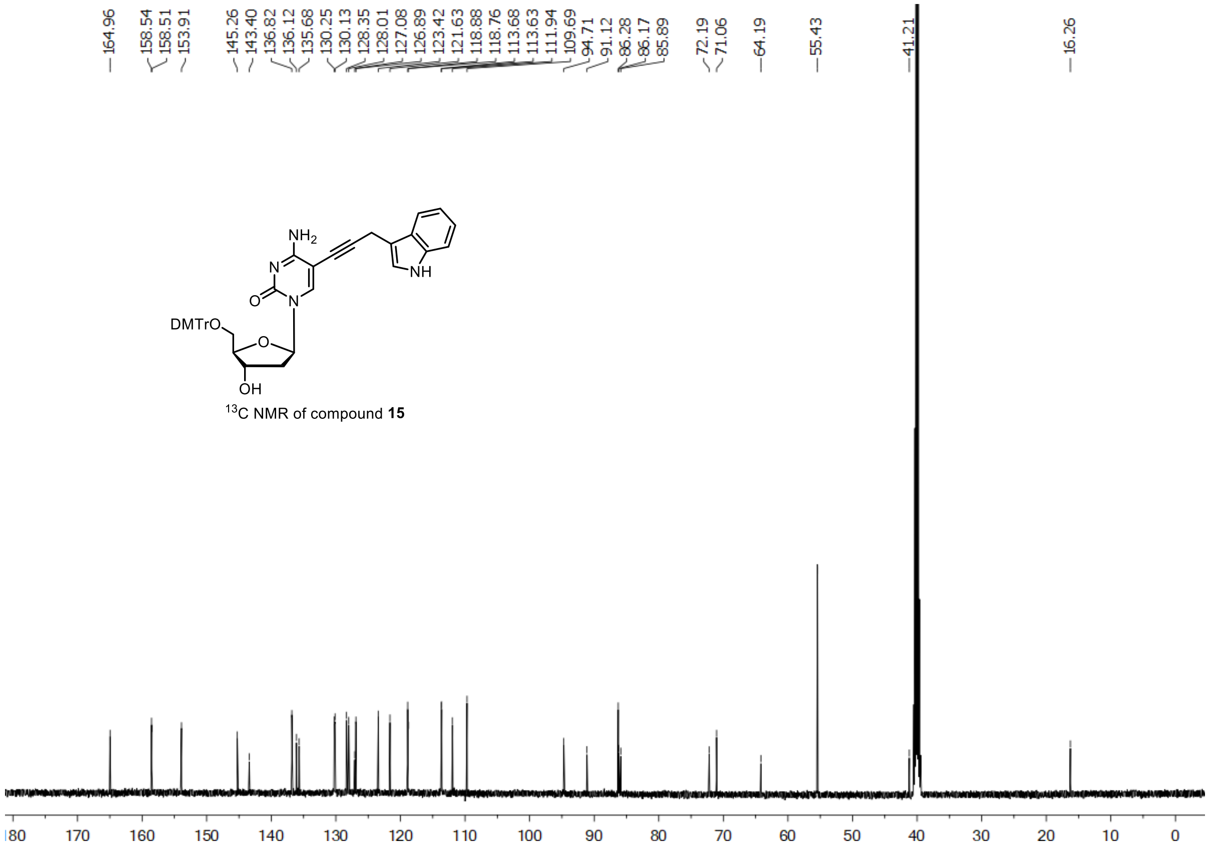


$^{13}C$ NMR of compound **15**

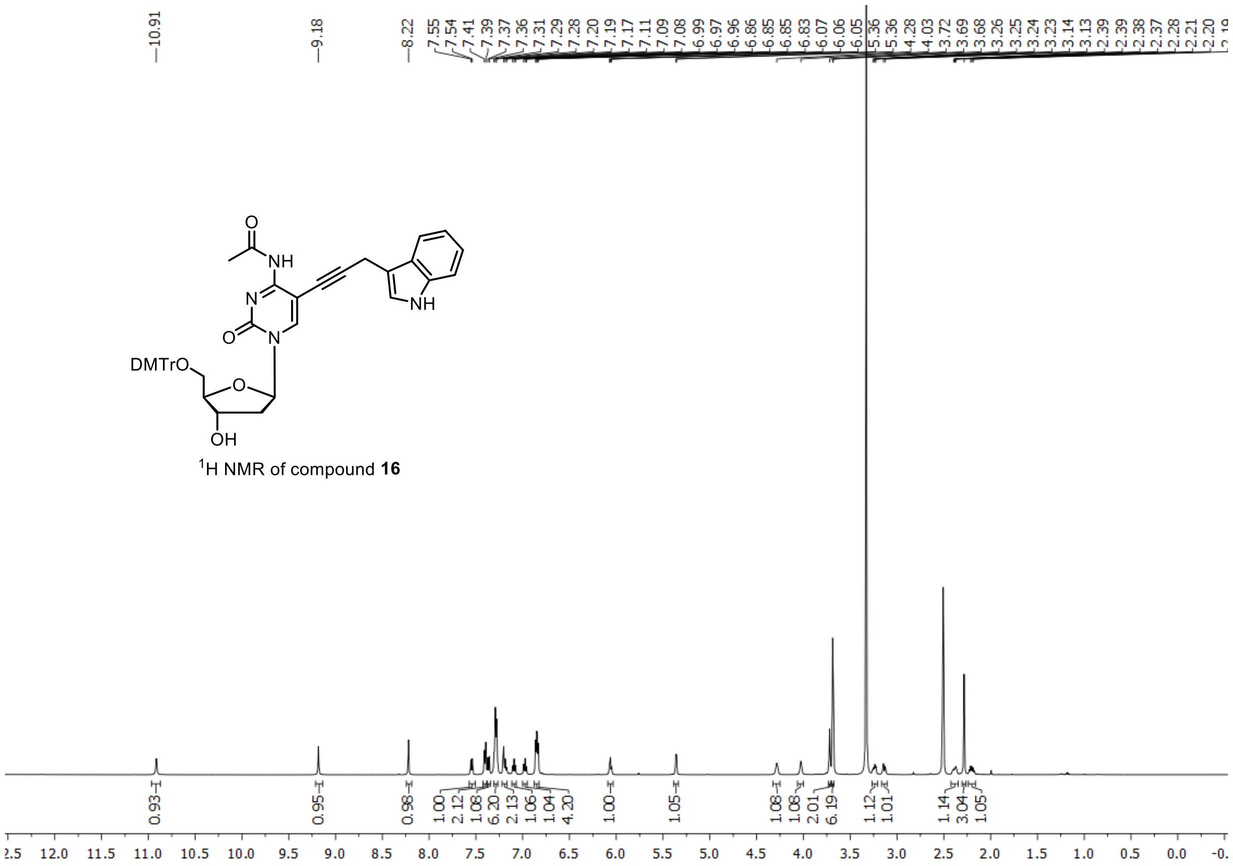

$^1H$ NMR of compound **16**

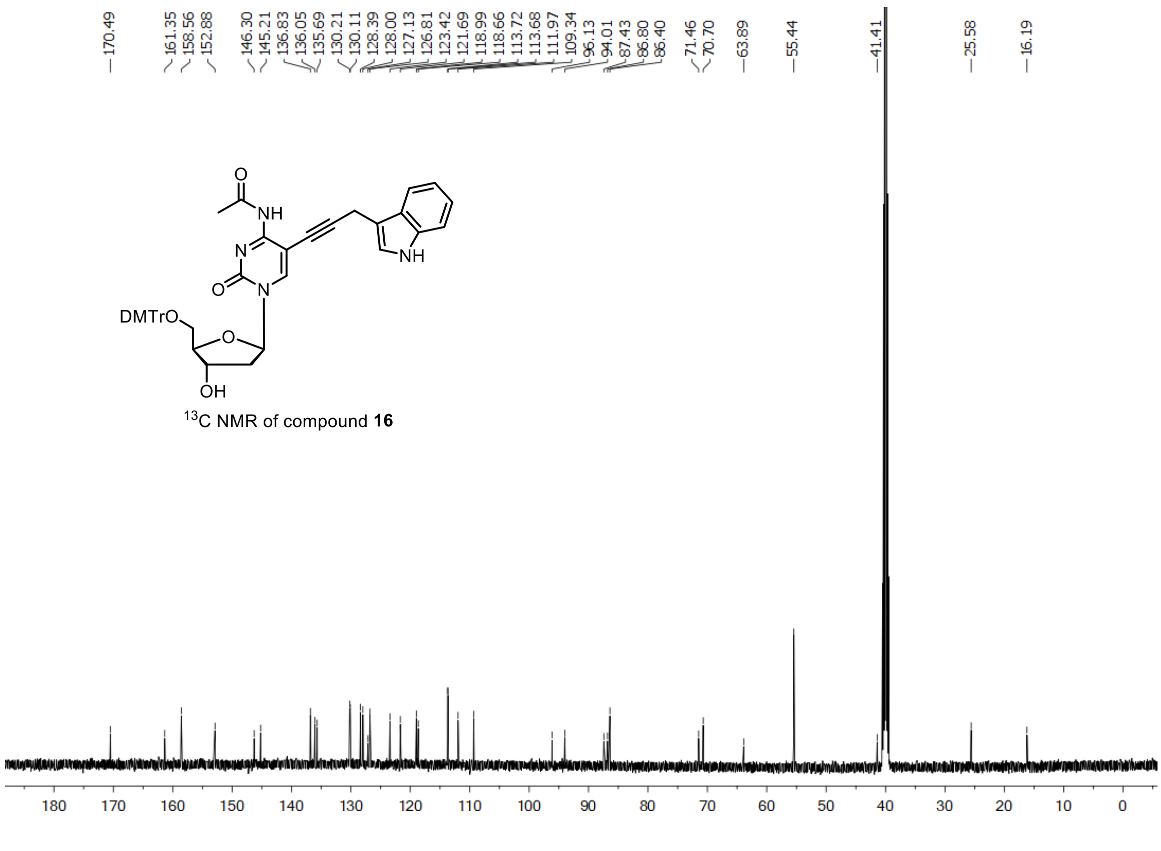

$^{13}C$ NMR of compound **16**

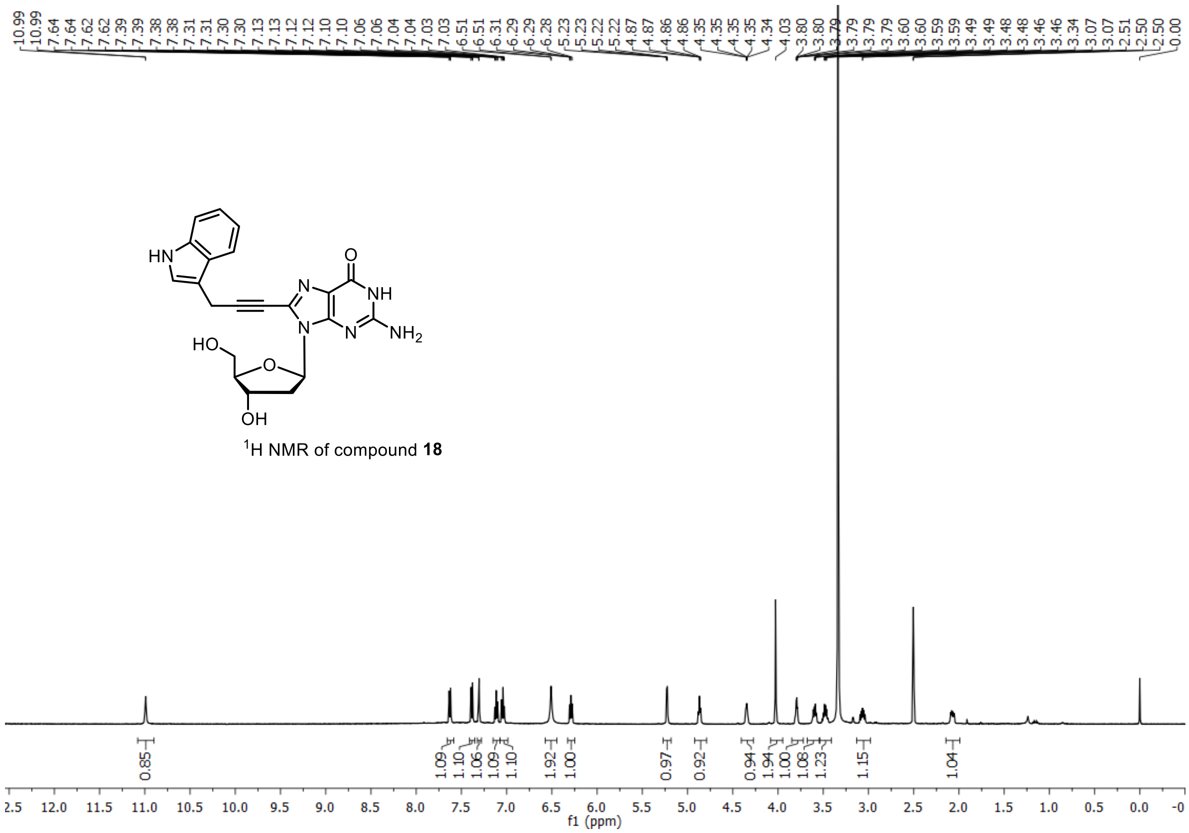


$^{1}$H NMR of compound **18**

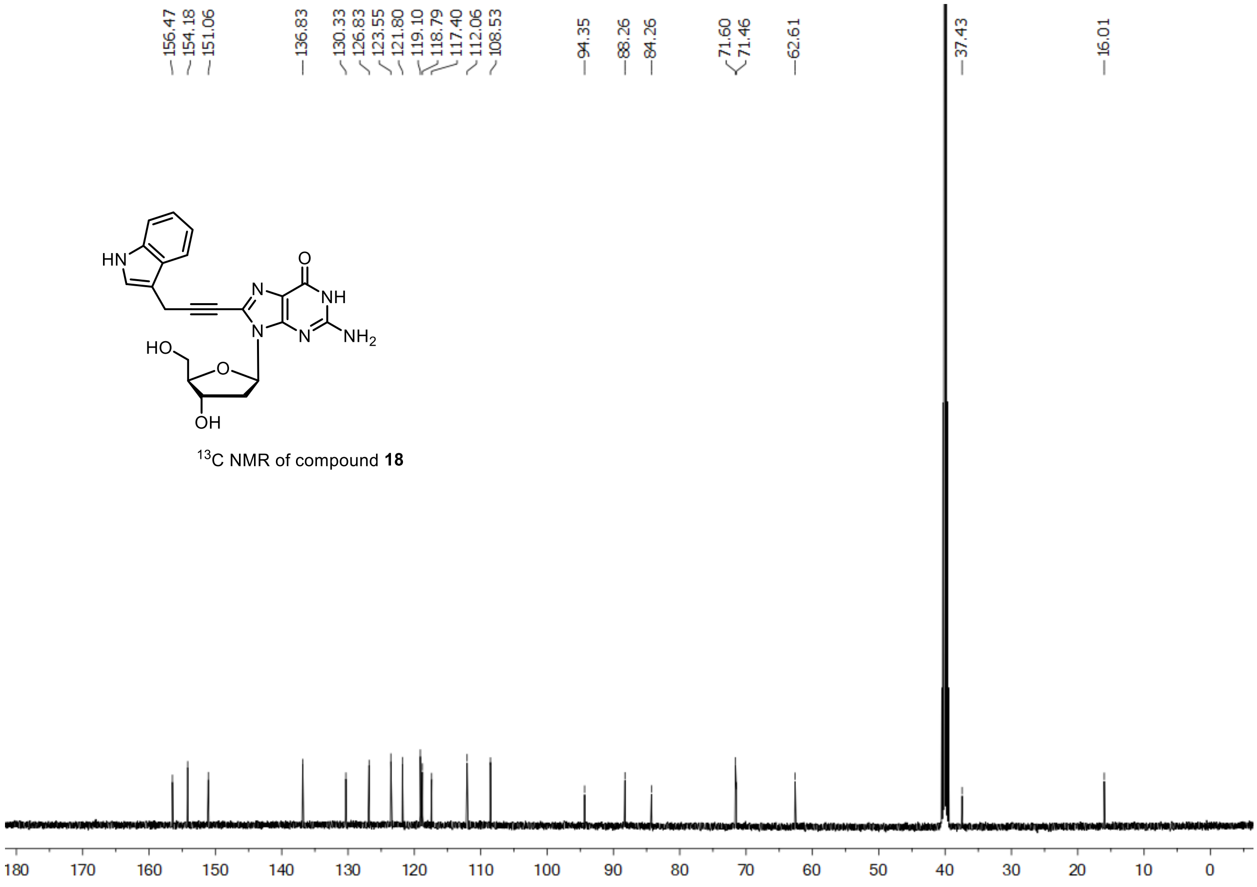


$^{13}$C NMR of compound **18**

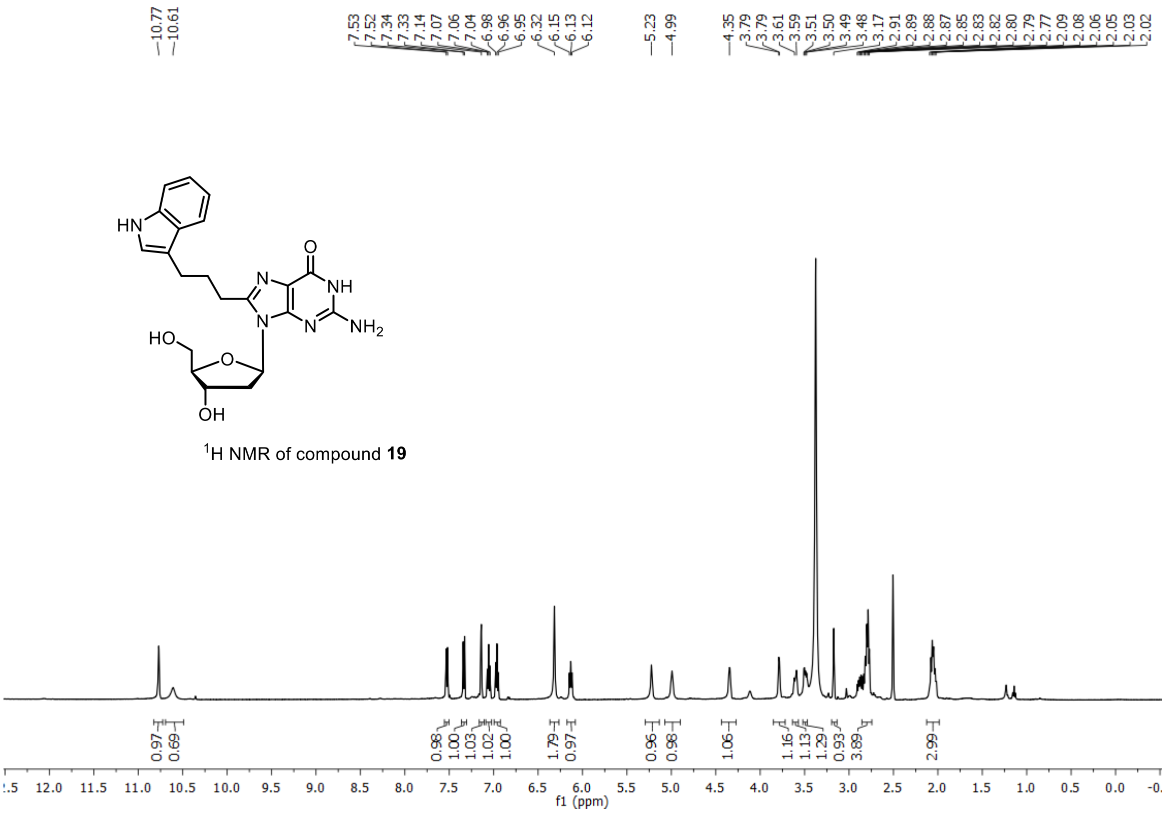


$^{1}$H NMR of compound **19**

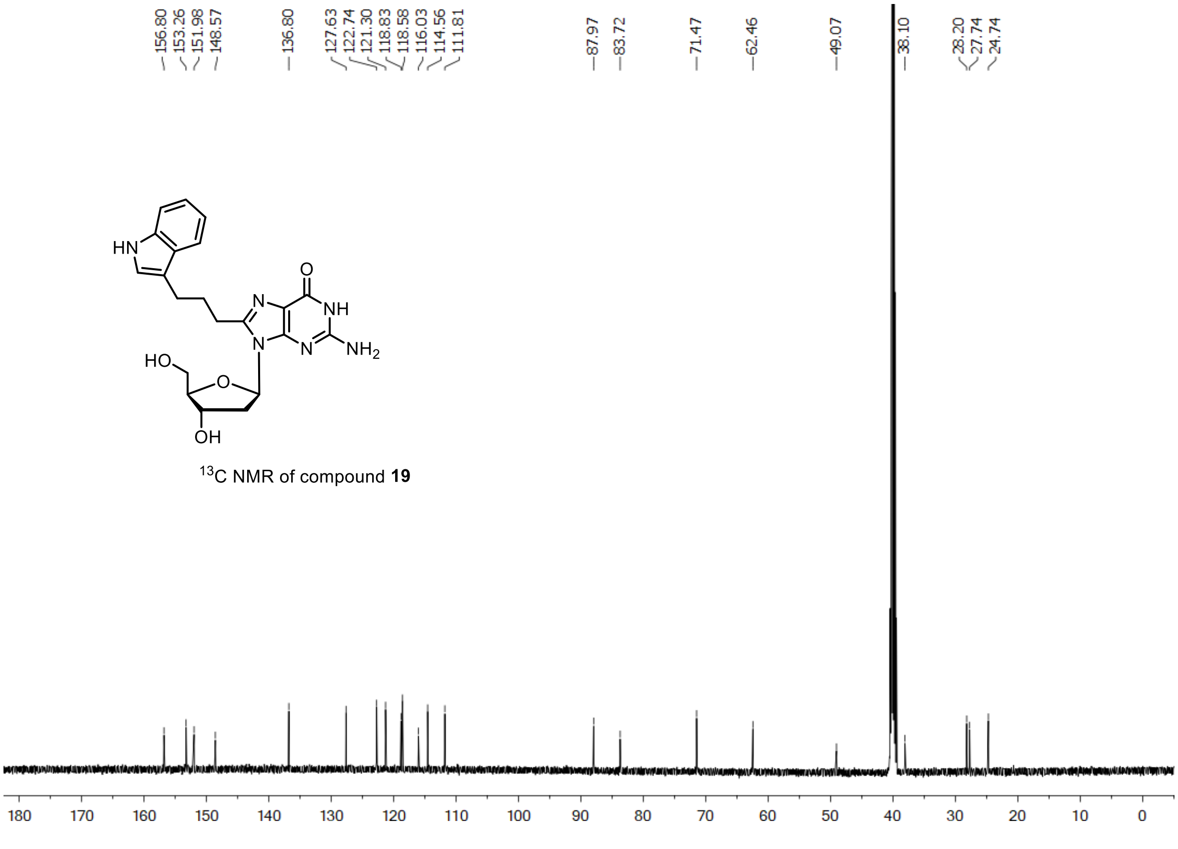


$^{13}$C NMR of compound **19**

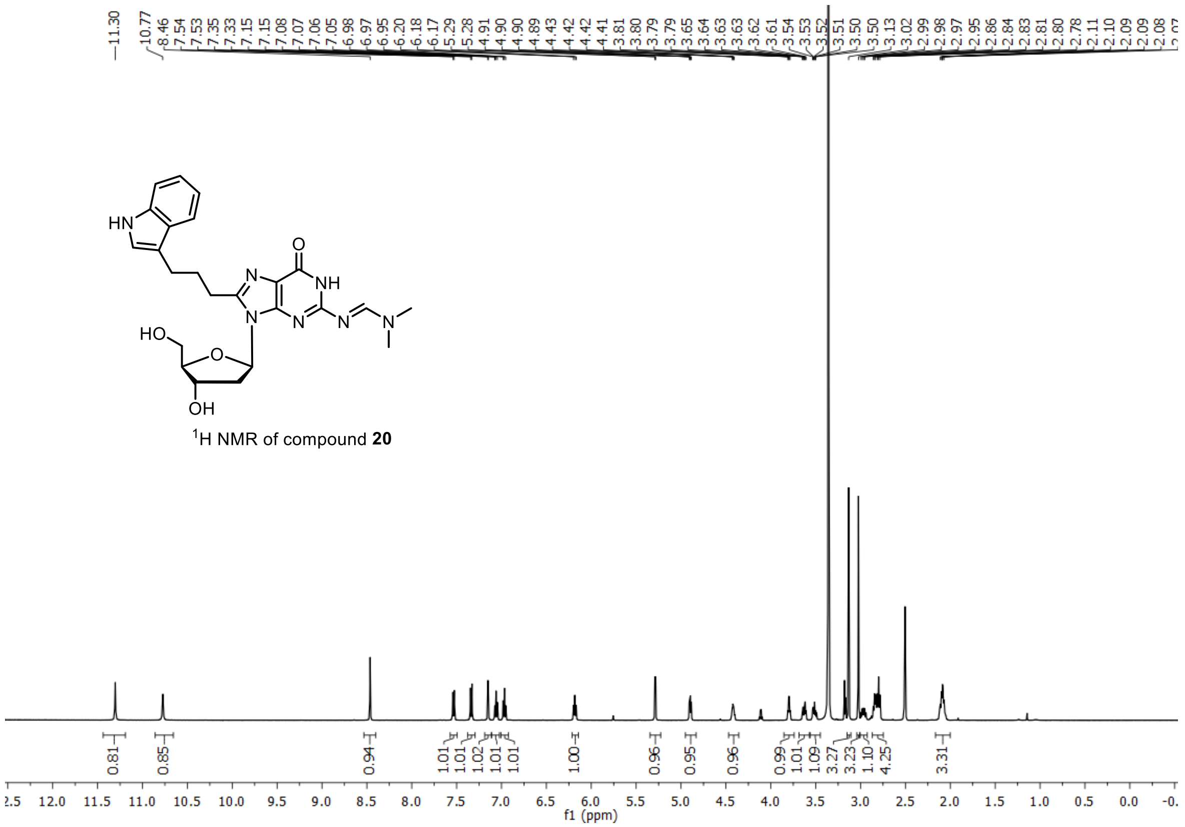

[1]H NMR of compound **20**

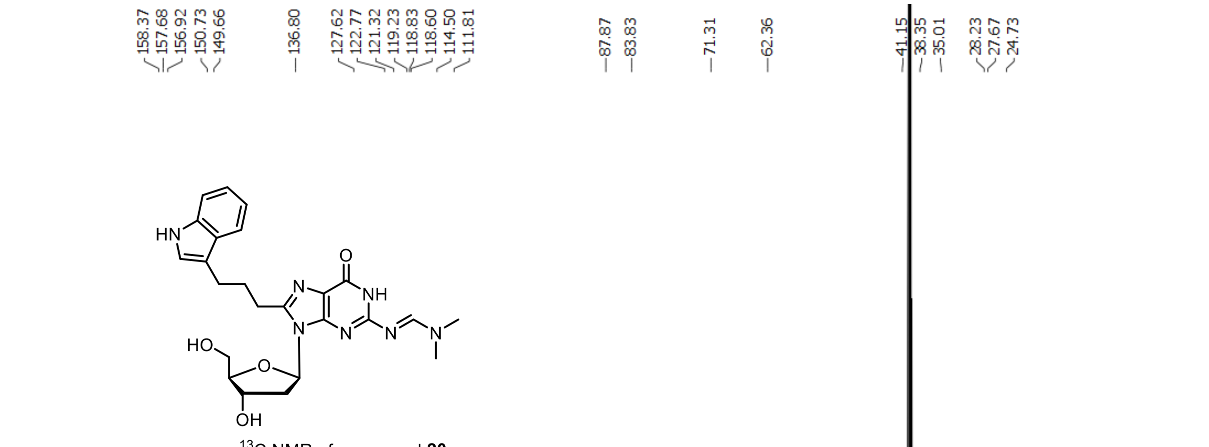

[13]C NMR of compound **20**

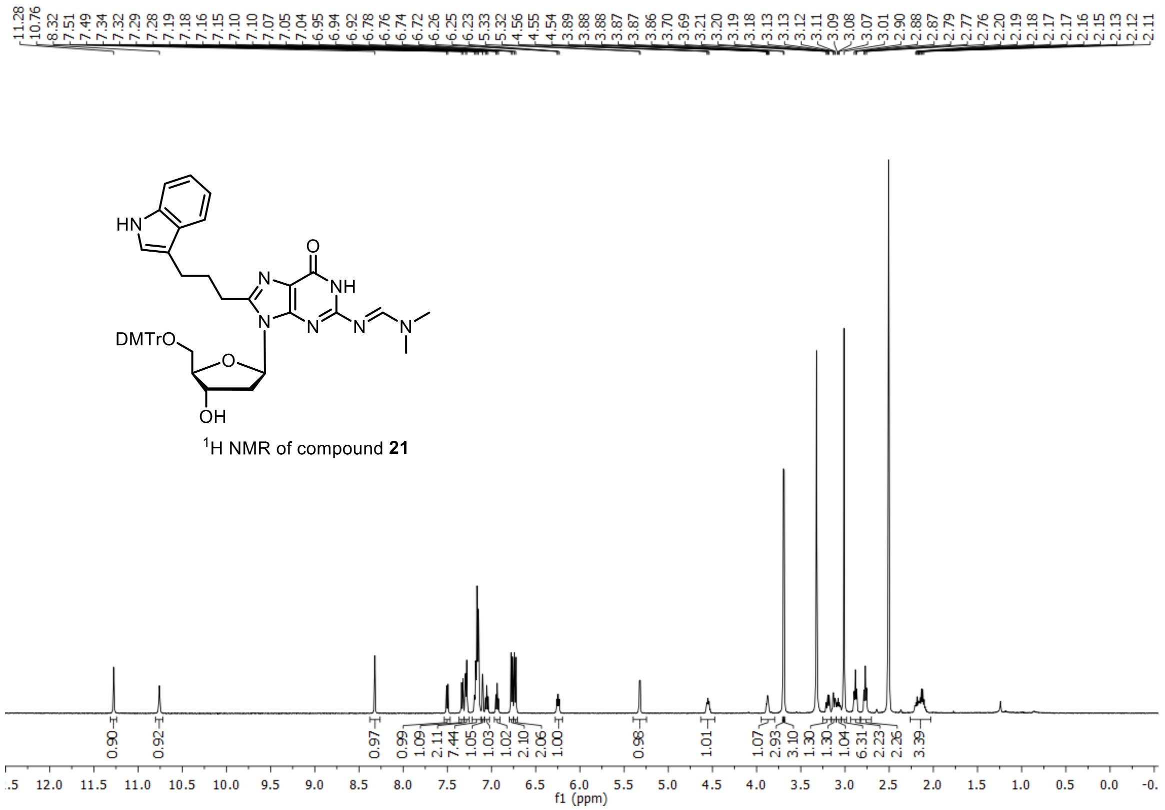

$^{1}H$ NMR of compound **21**

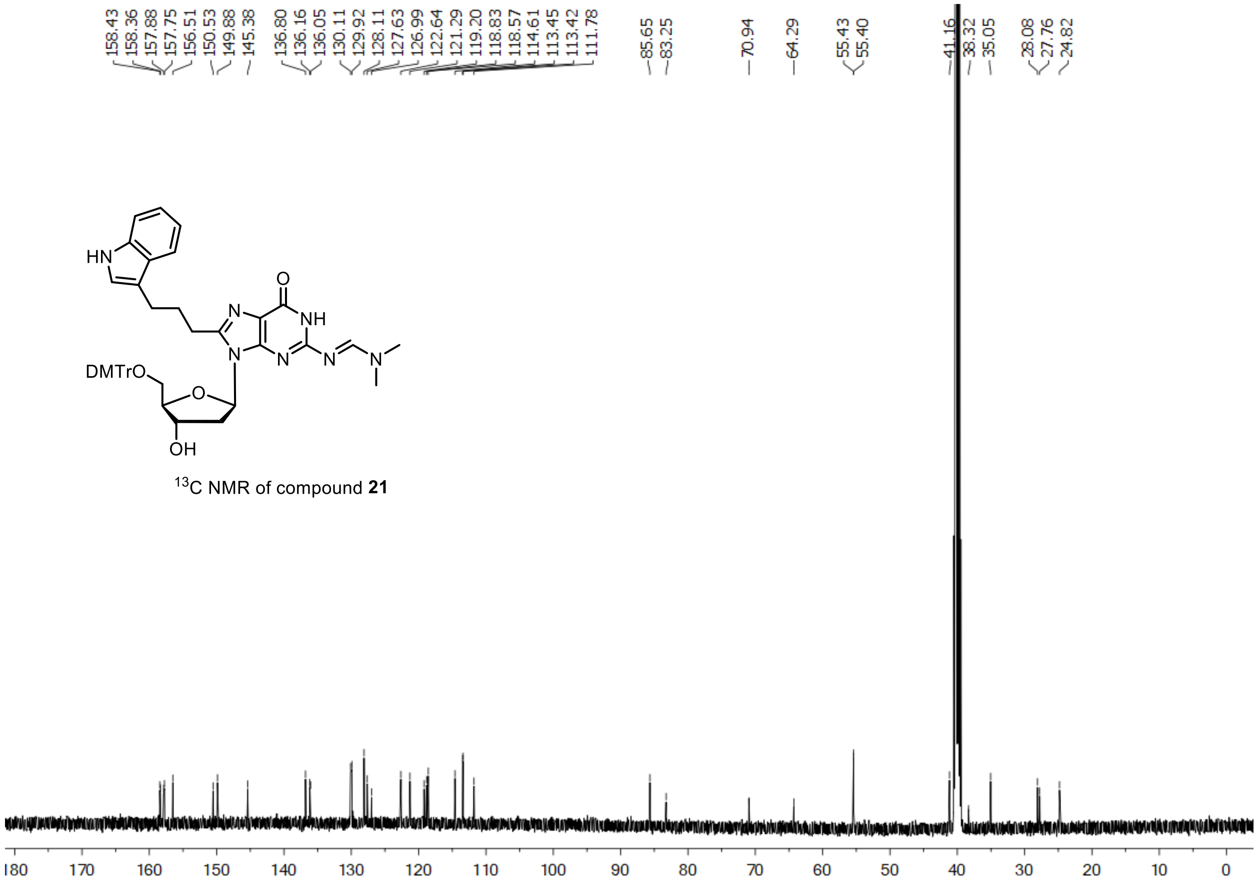

$^{13}C$ NMR of compound **21**

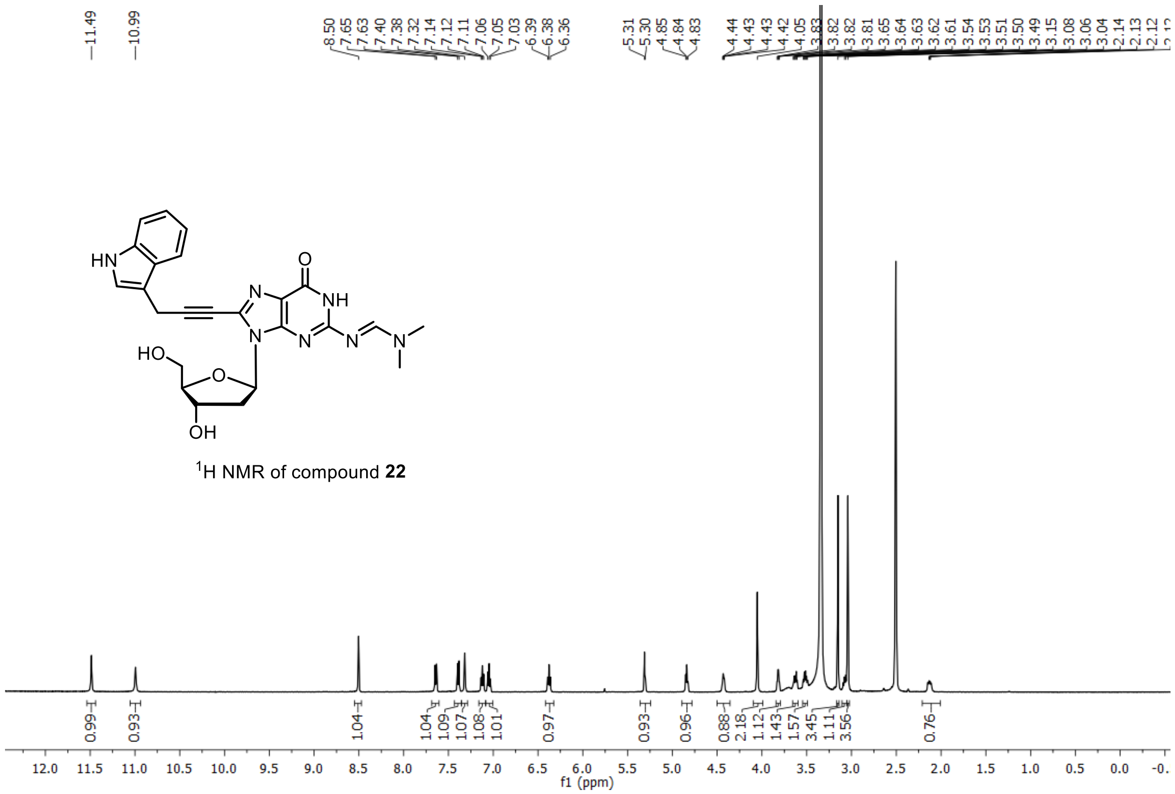

$^{1}$H NMR of compound **22**

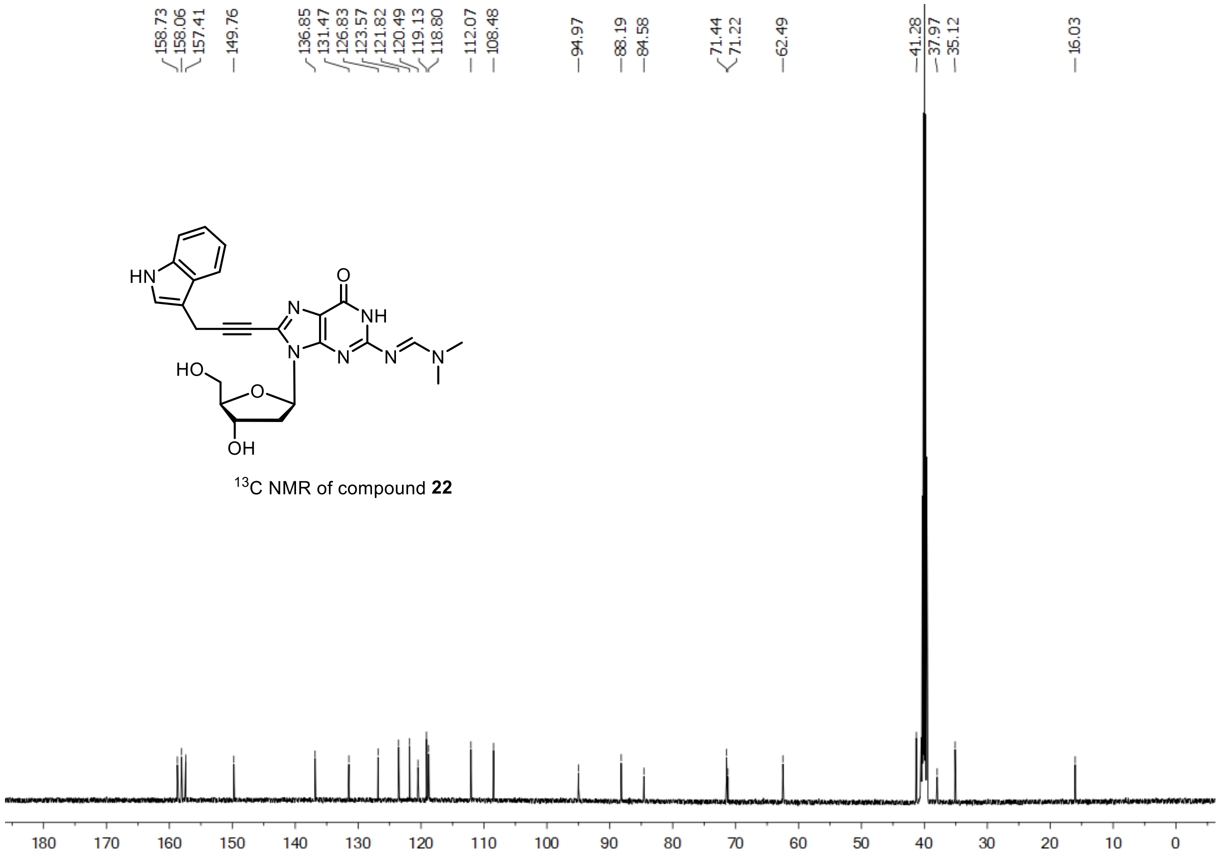

$^{13}$C NMR of compound **22**

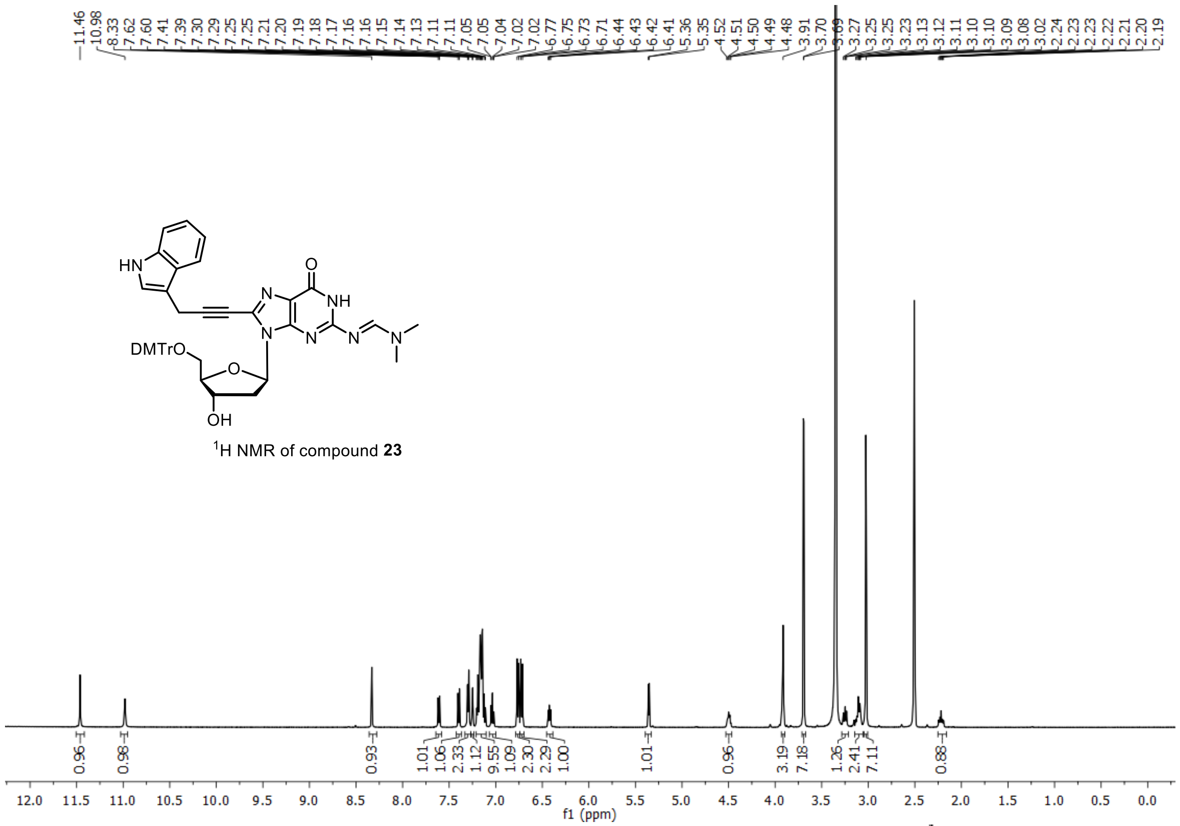

[1]H NMR of compound **23**

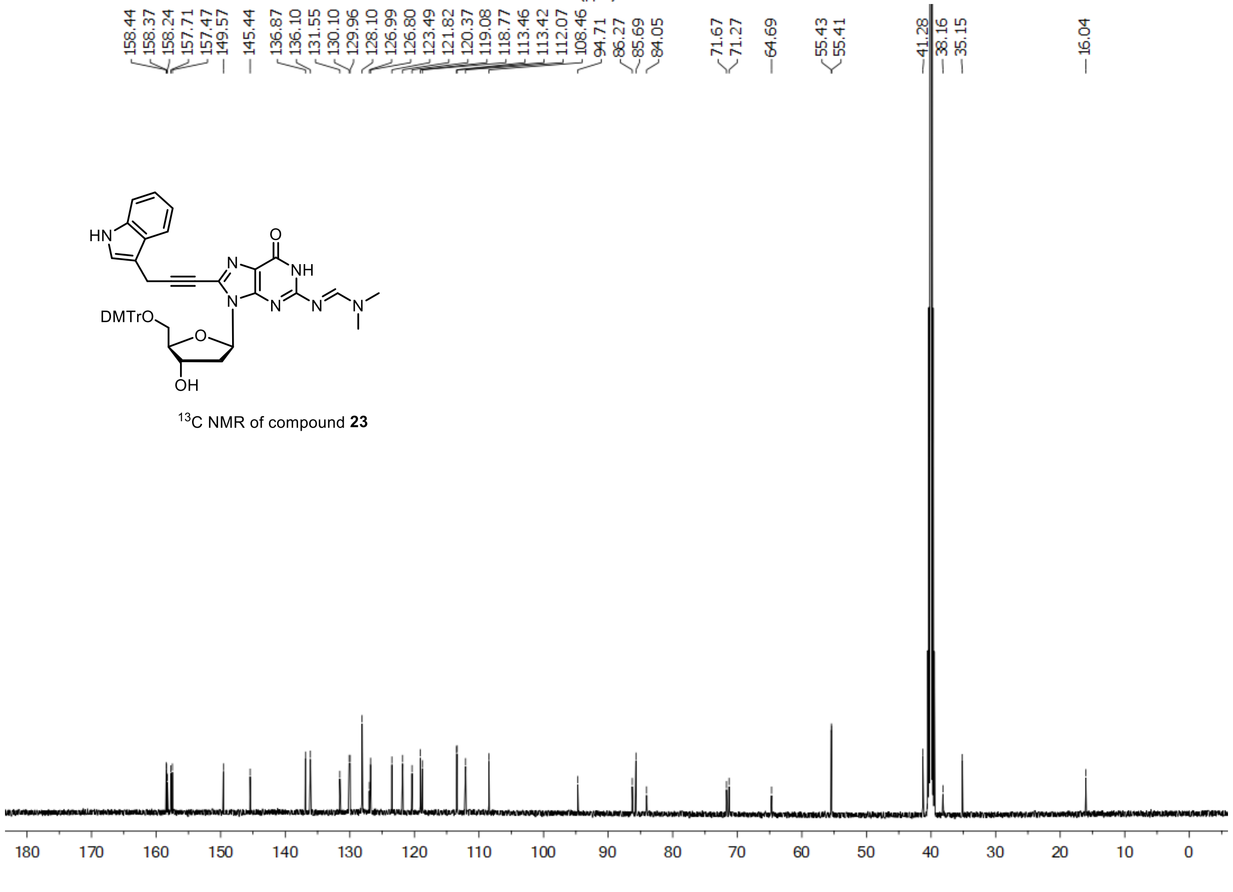

[13]C NMR of compound **23**

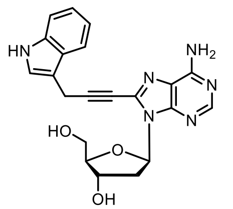


$^{1}H$ NMR of compound **25**

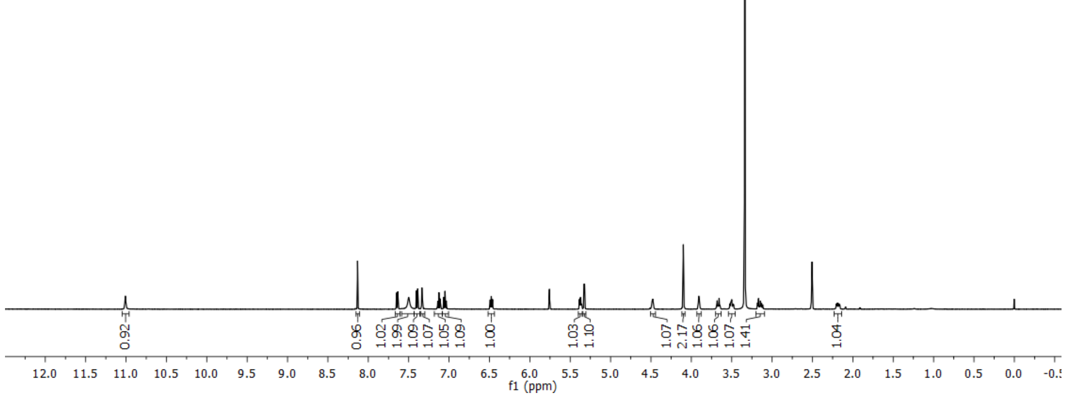


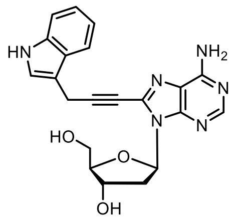


$^{13}C$ NMR of compound **25**

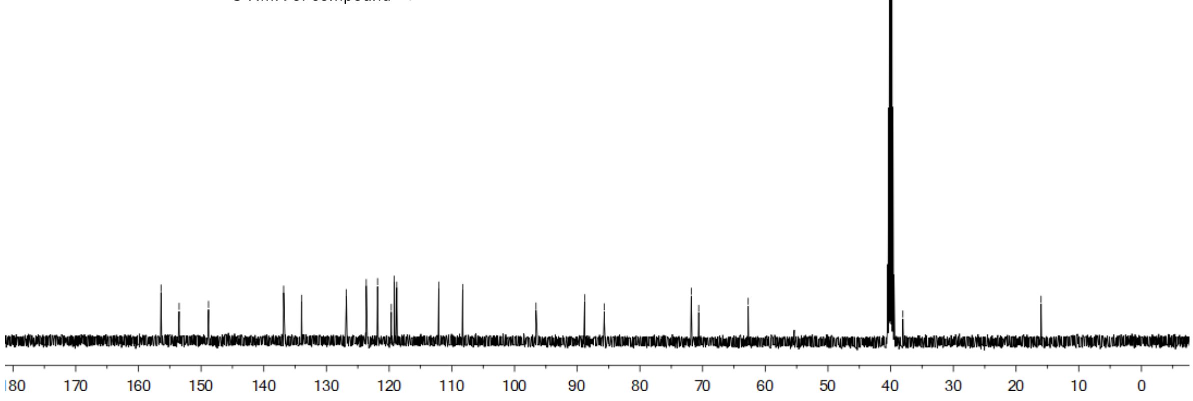

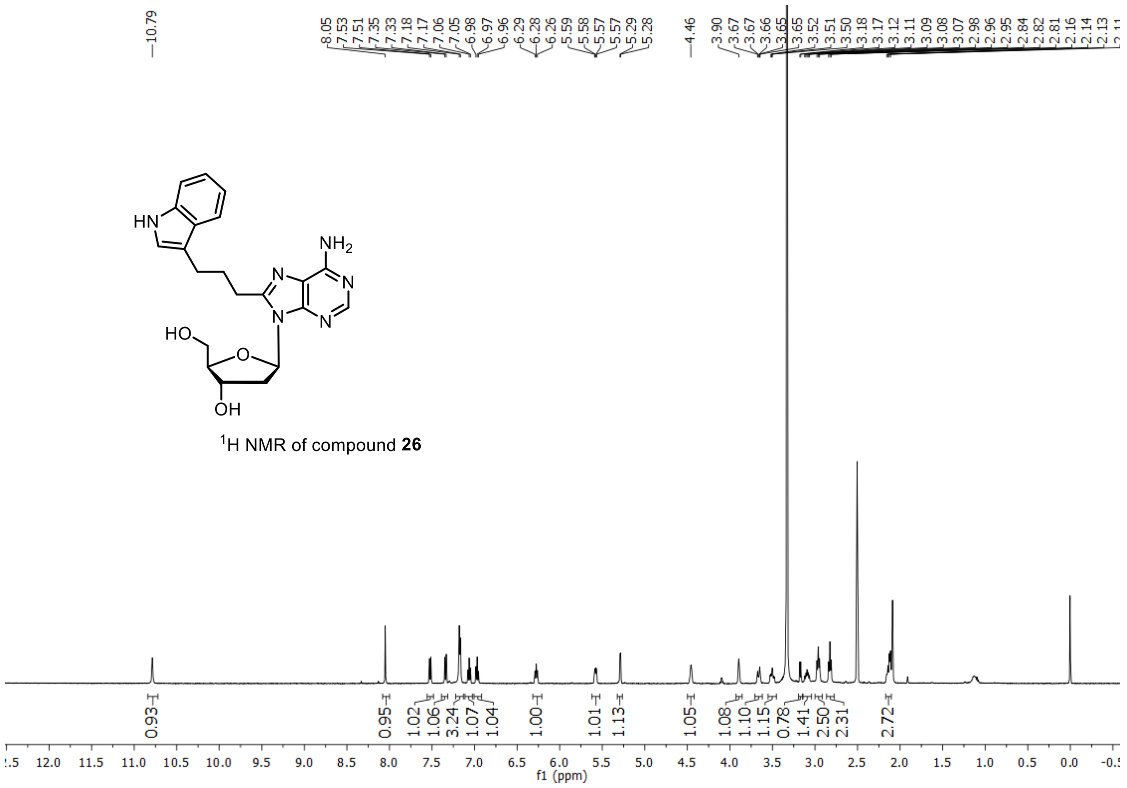

[1]H NMR of compound **26**

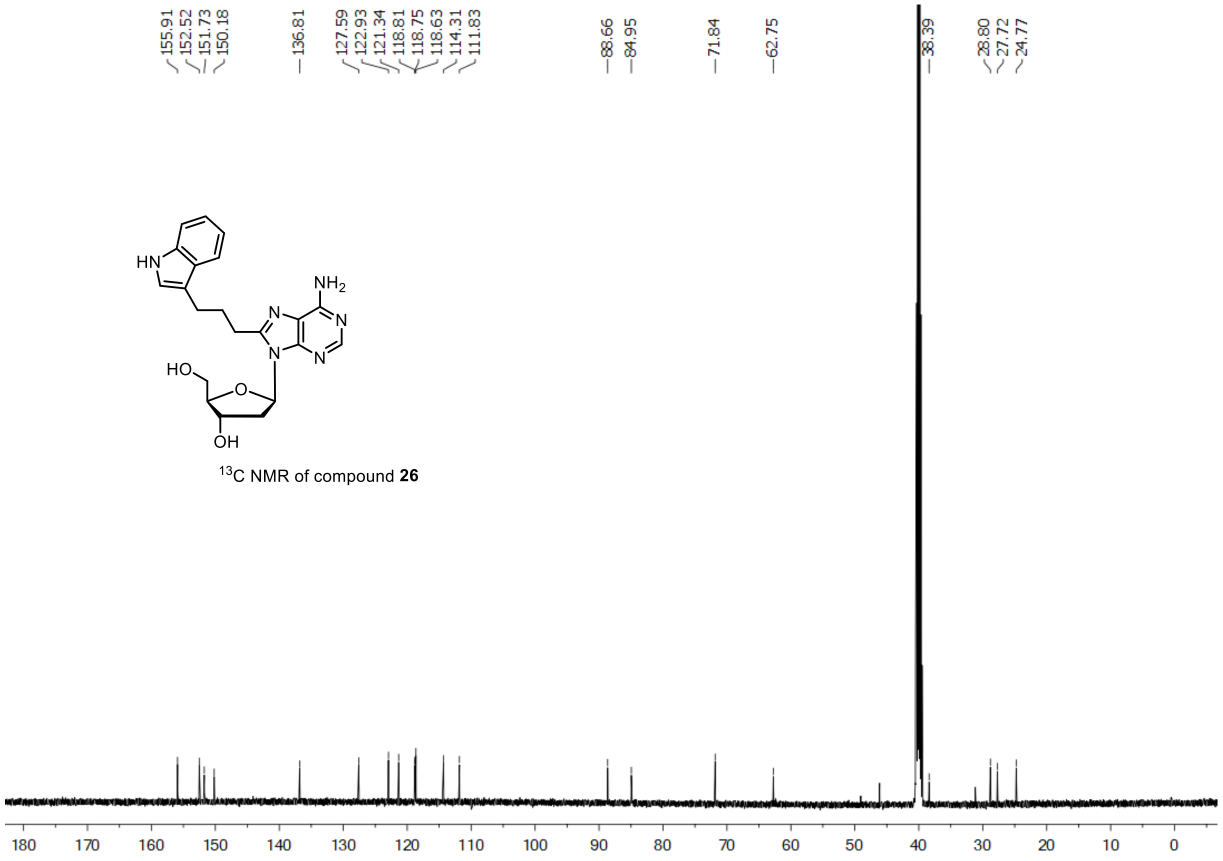

[13]C NMR of compound **26**

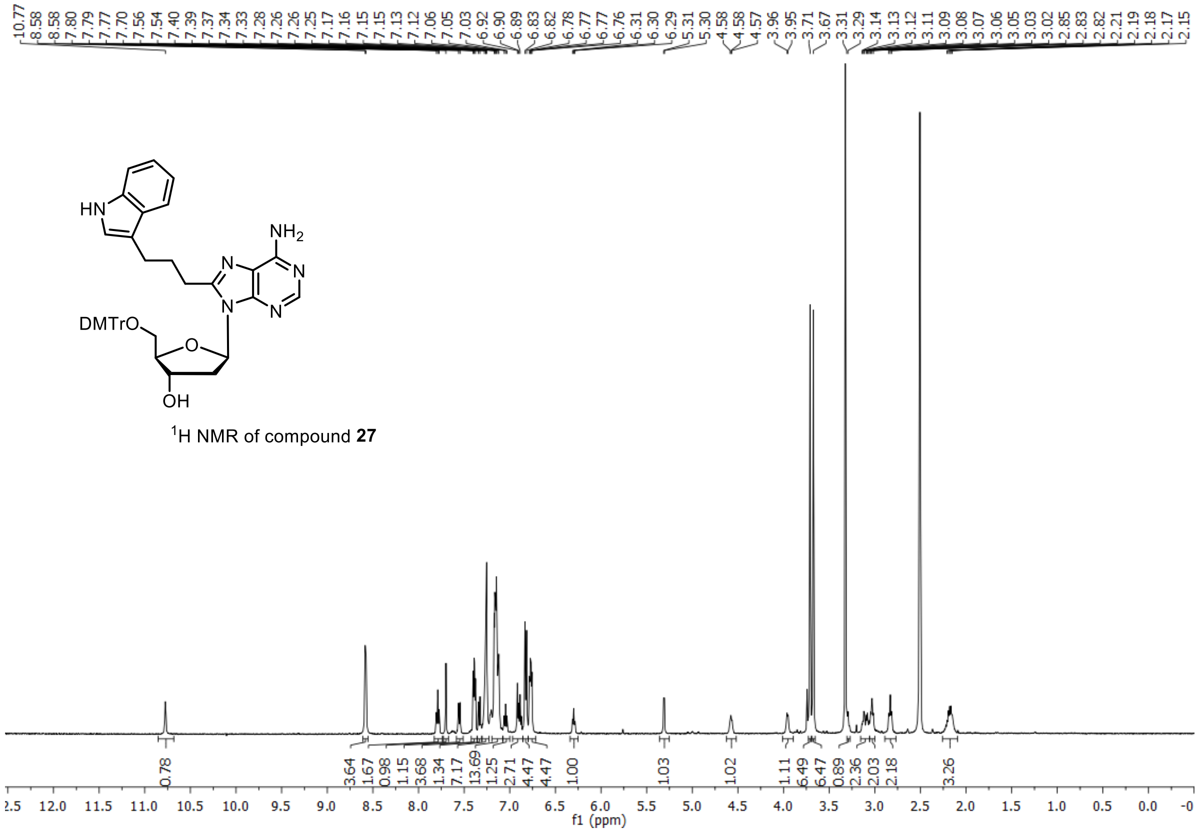

$^{1}$H NMR of compound **27**

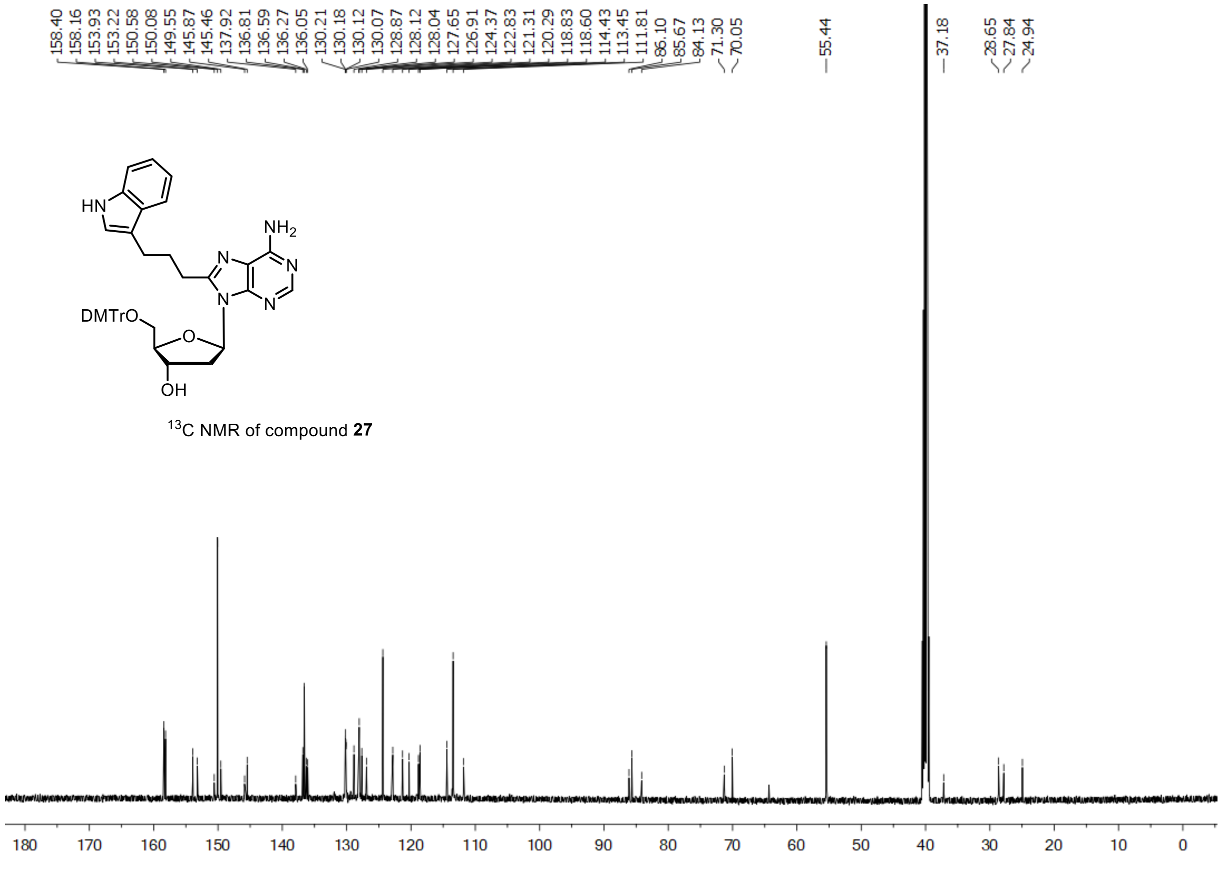

$^{13}$C NMR of compound **27**

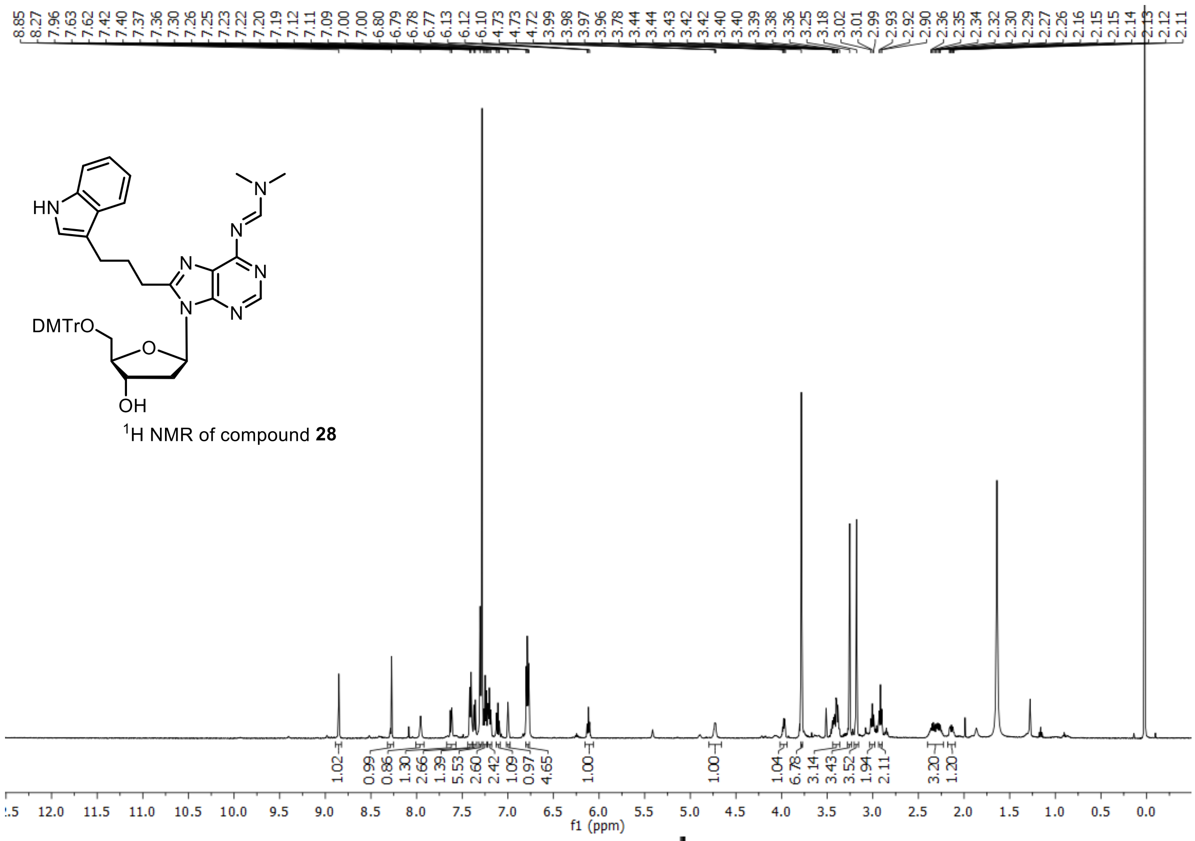


$^{1}$H NMR of compound **28**

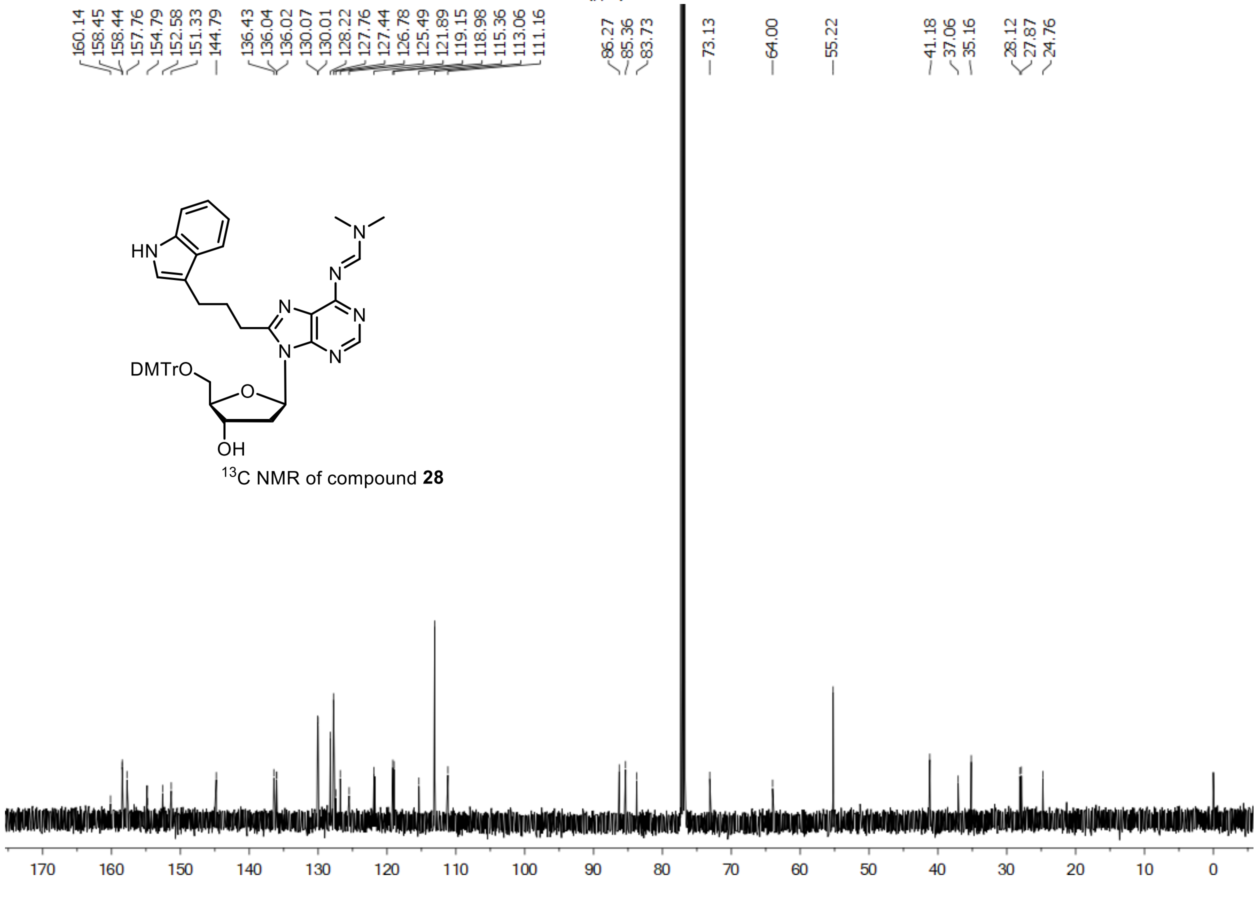


$^{13}$C NMR of compound **28**

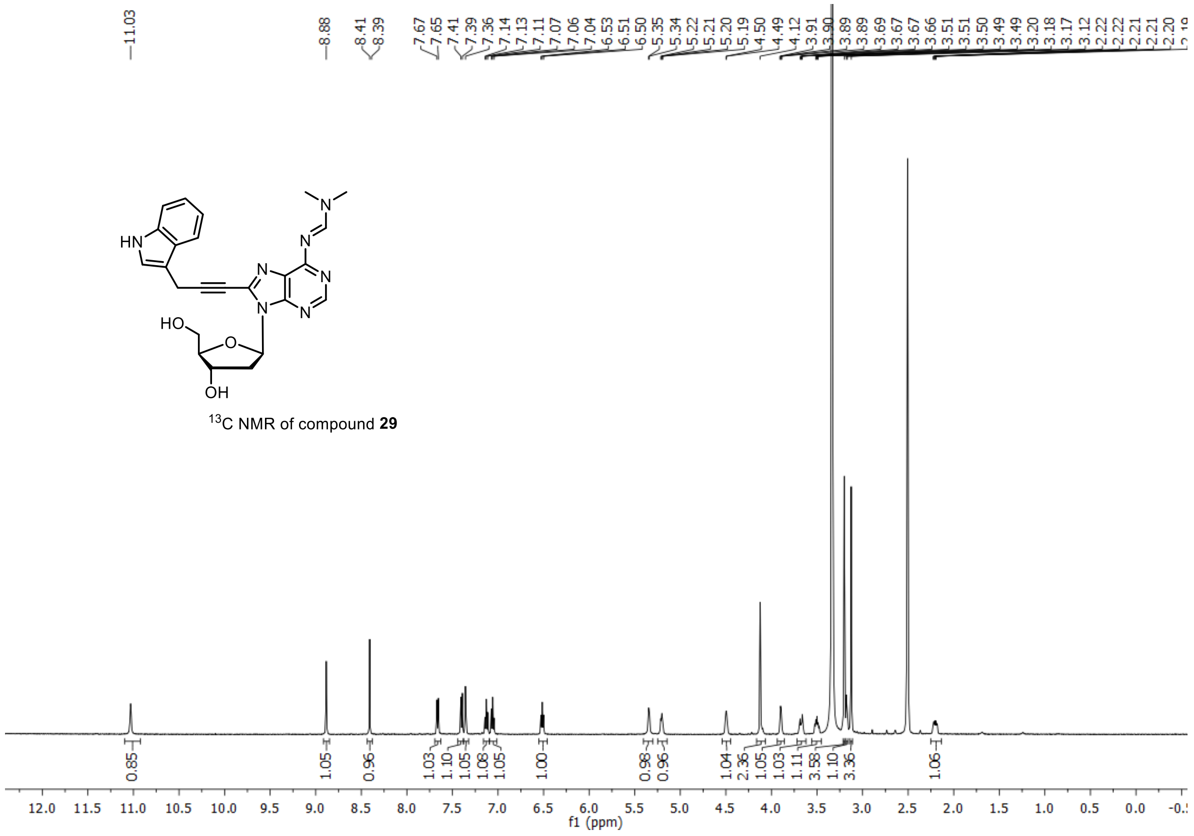


$^{13}$C NMR of compound **29**

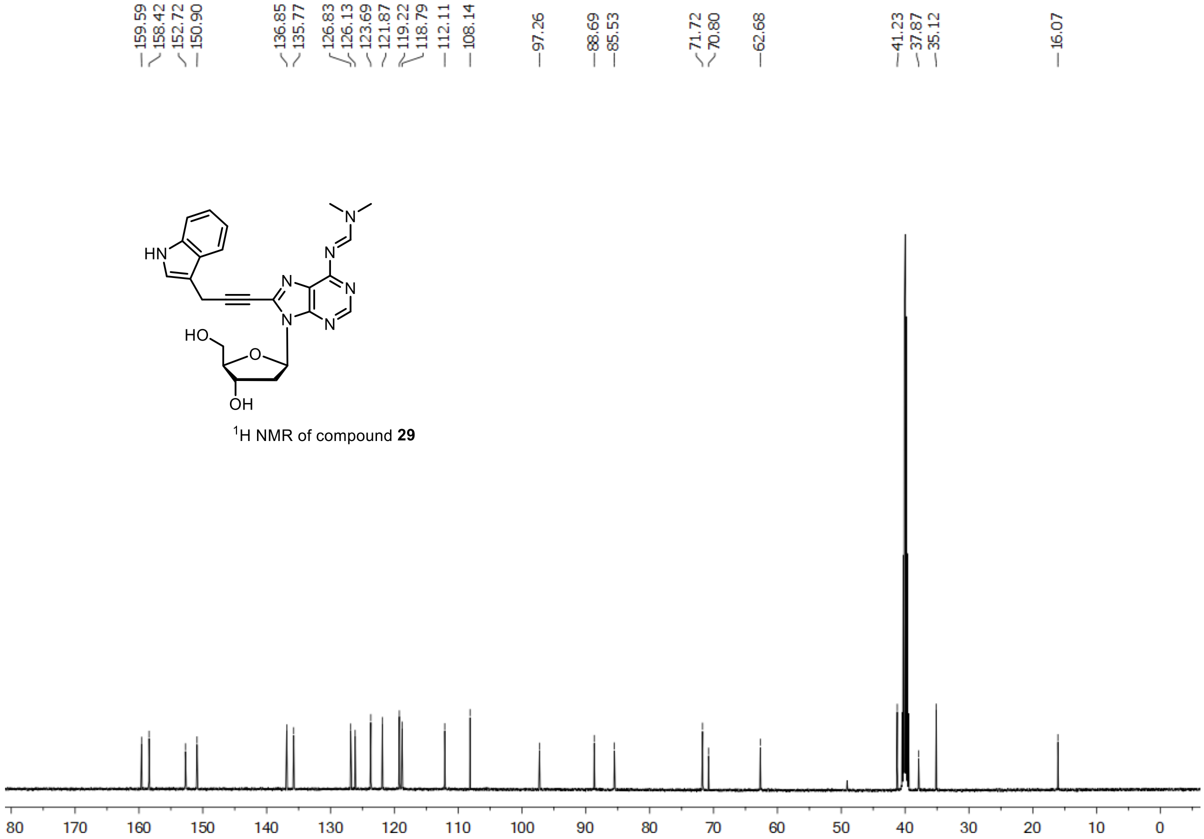


$^{1}$H NMR of compound **29**

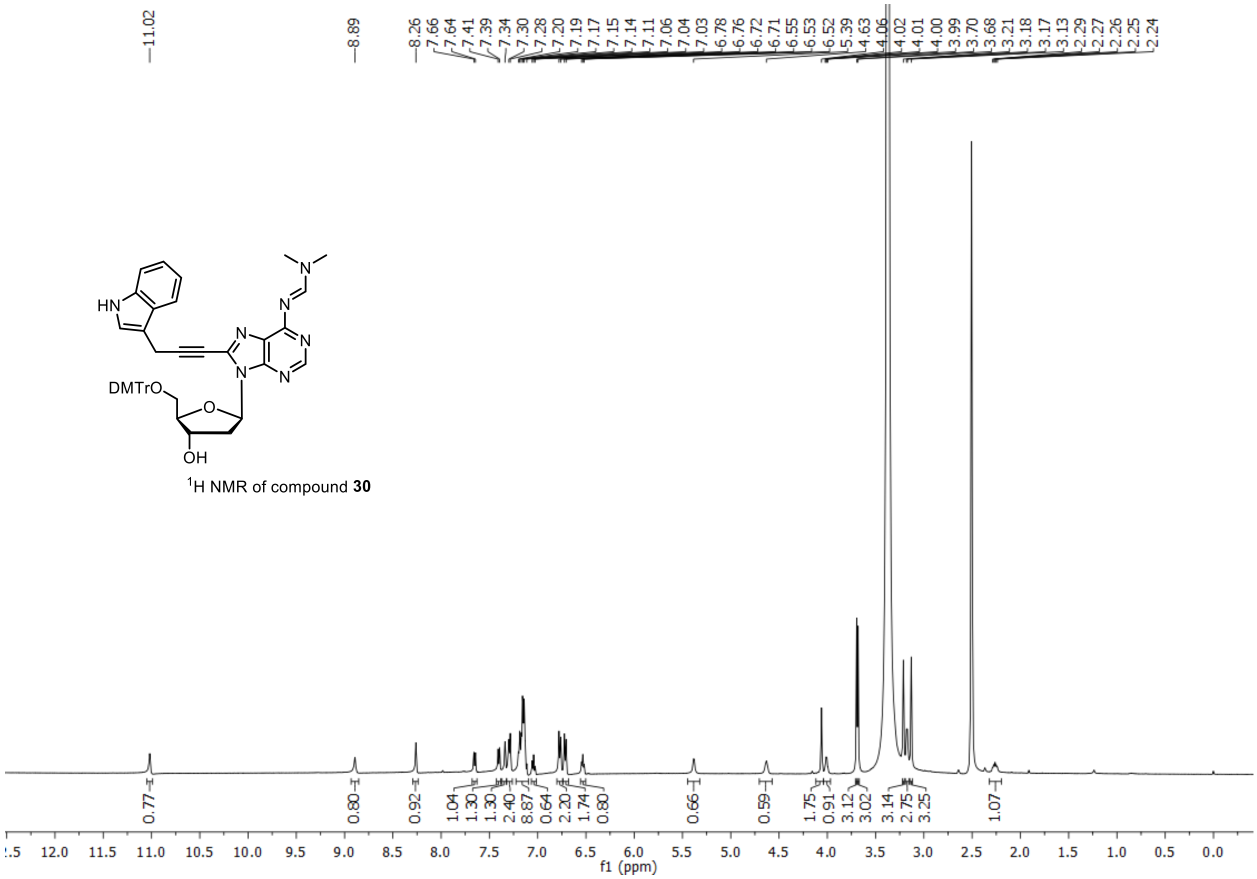


[1]H NMR of compound **30**

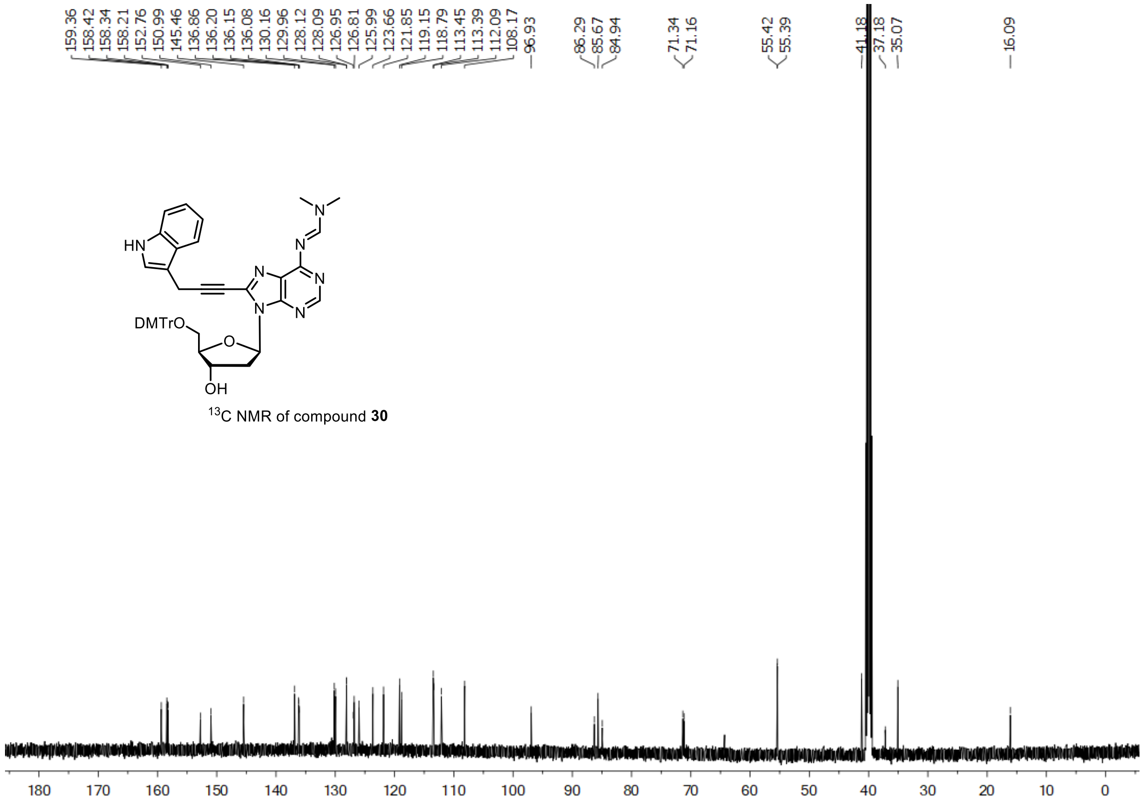


[13]C NMR of compound **30**

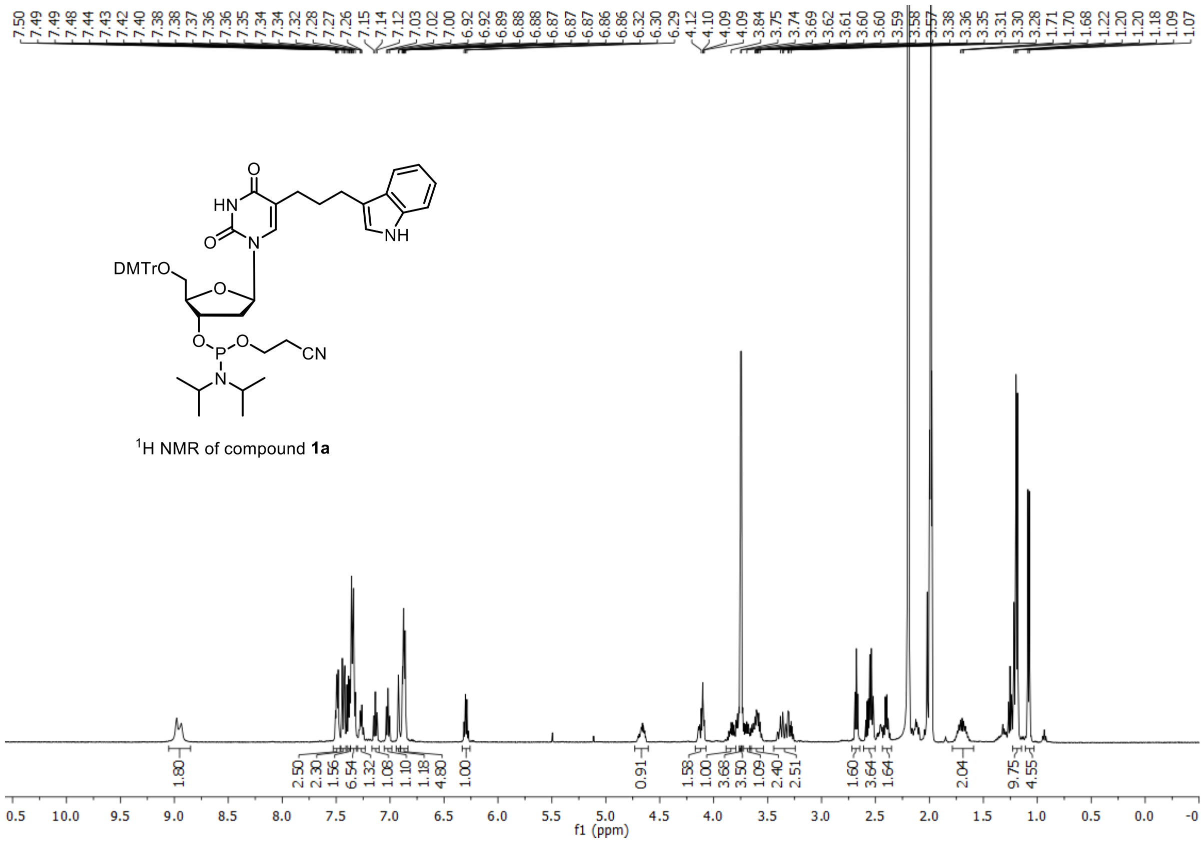


$^{1}$H NMR of compound **1a**

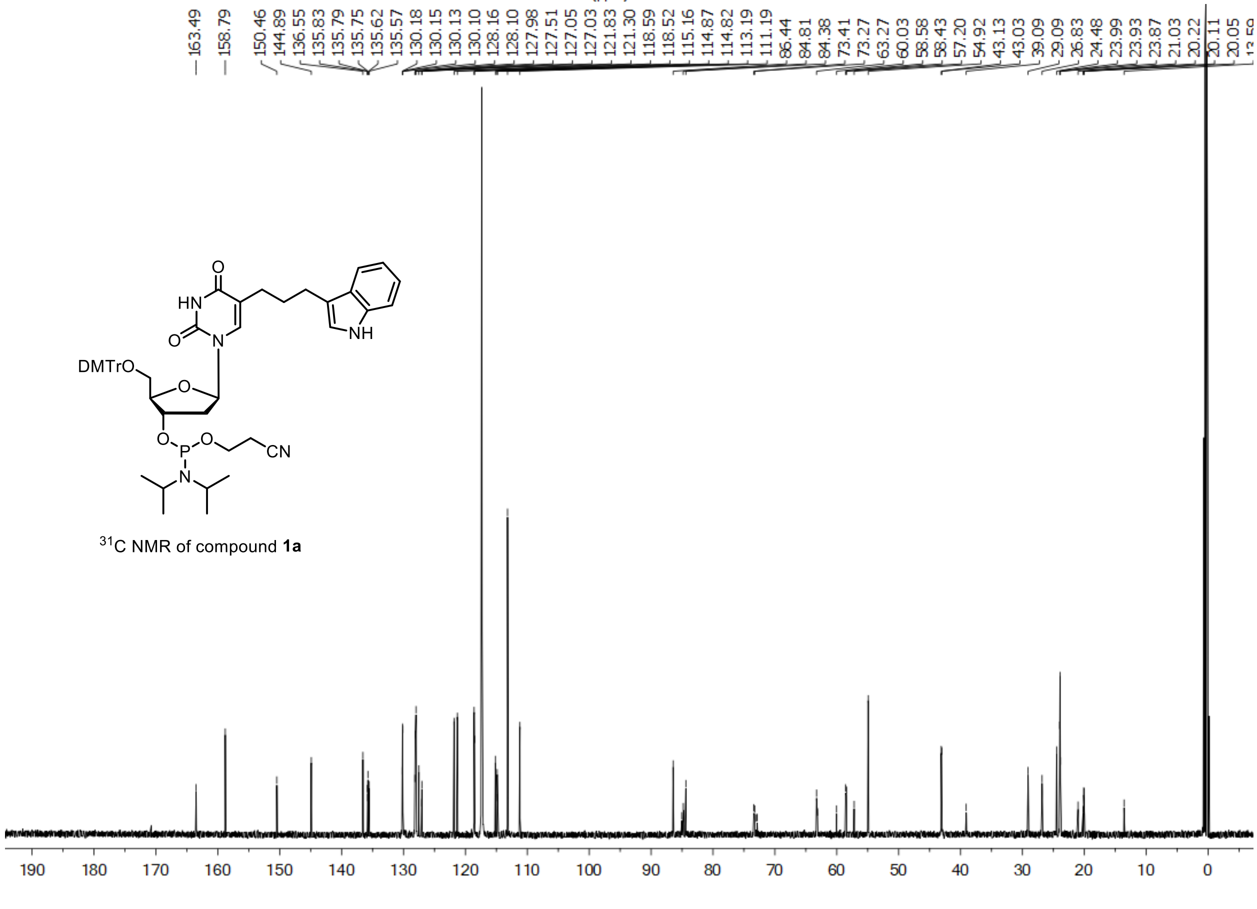


$^{31}$C NMR of compound **1a**

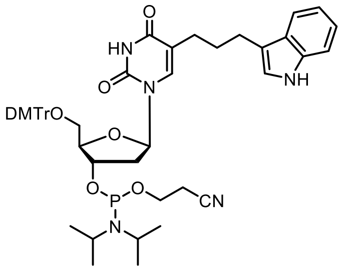

$^{31}$P NMR of compound **1a**

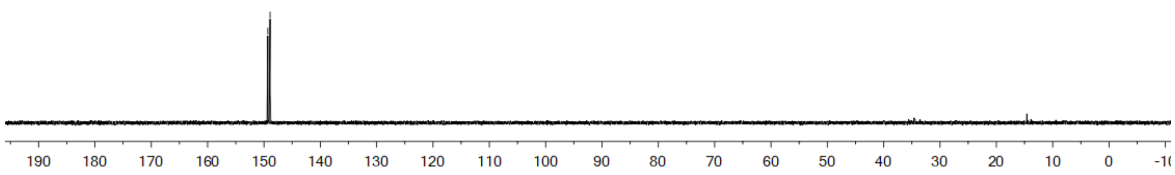


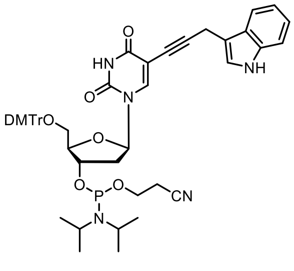

$^{1}$H NMR of compound **1b**

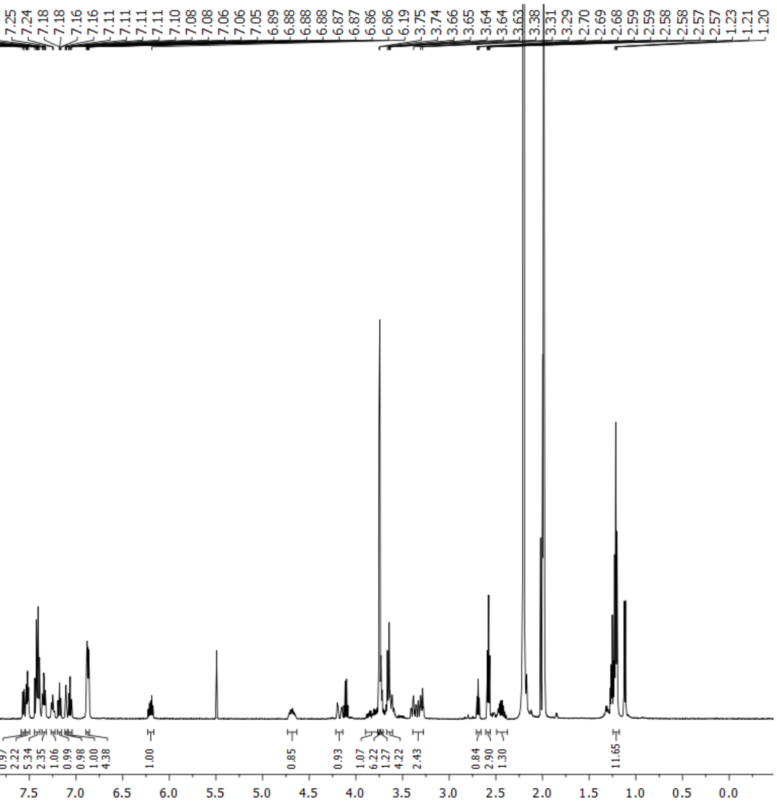

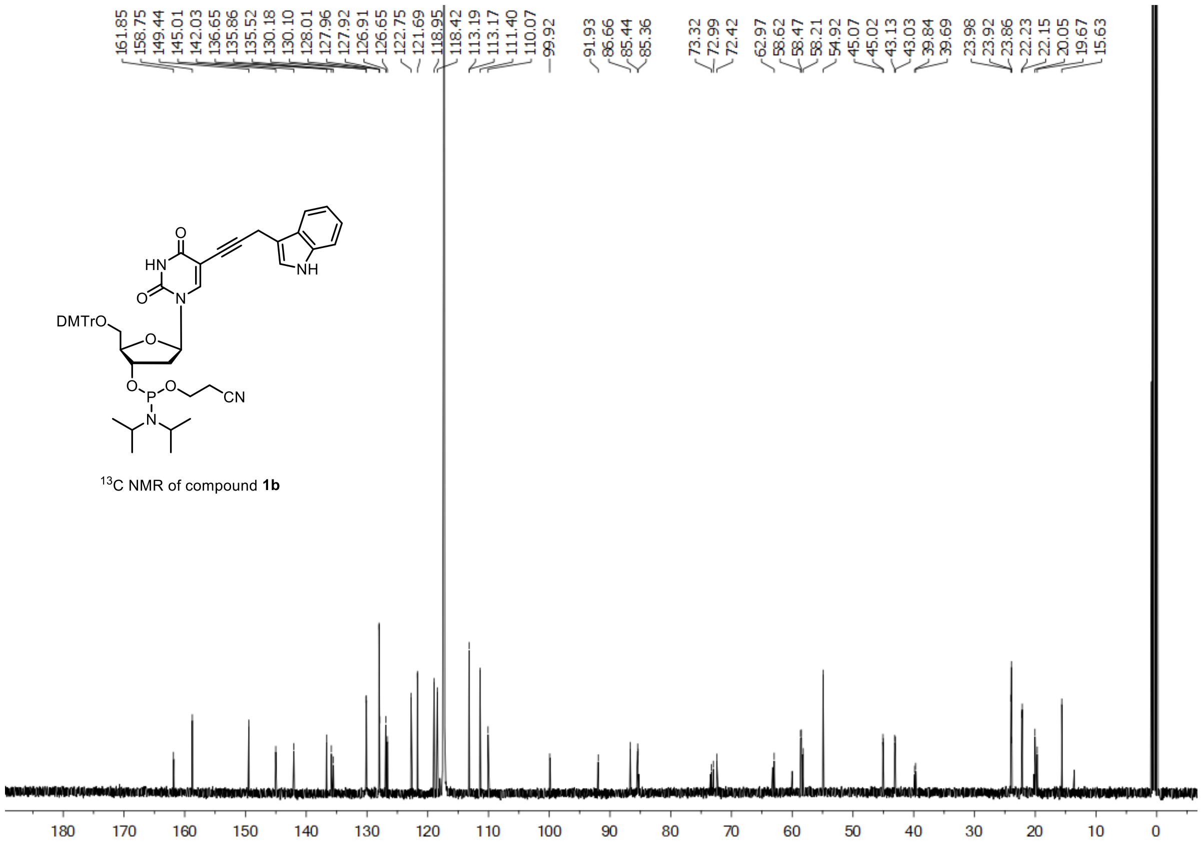


$^{13}$C NMR of compound **1b**

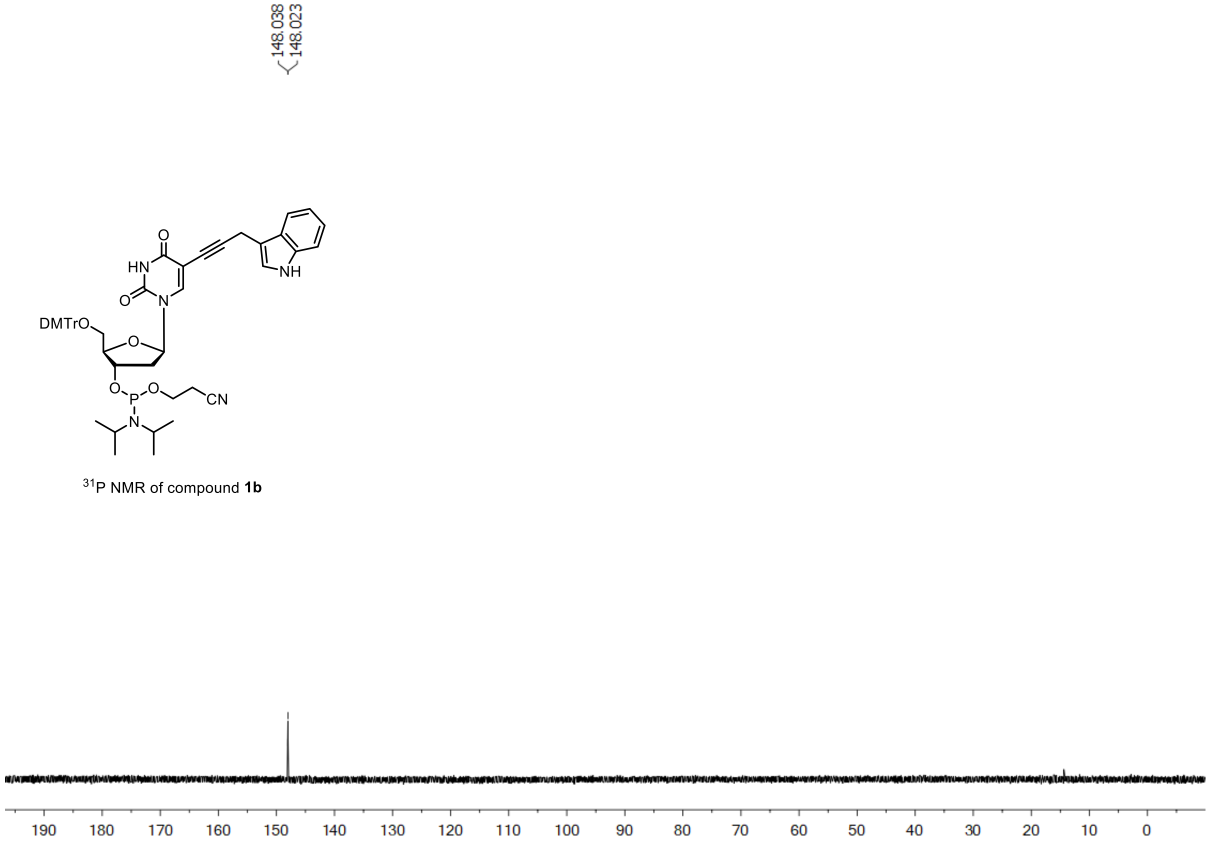


$^{31}$P NMR of compound **1b**

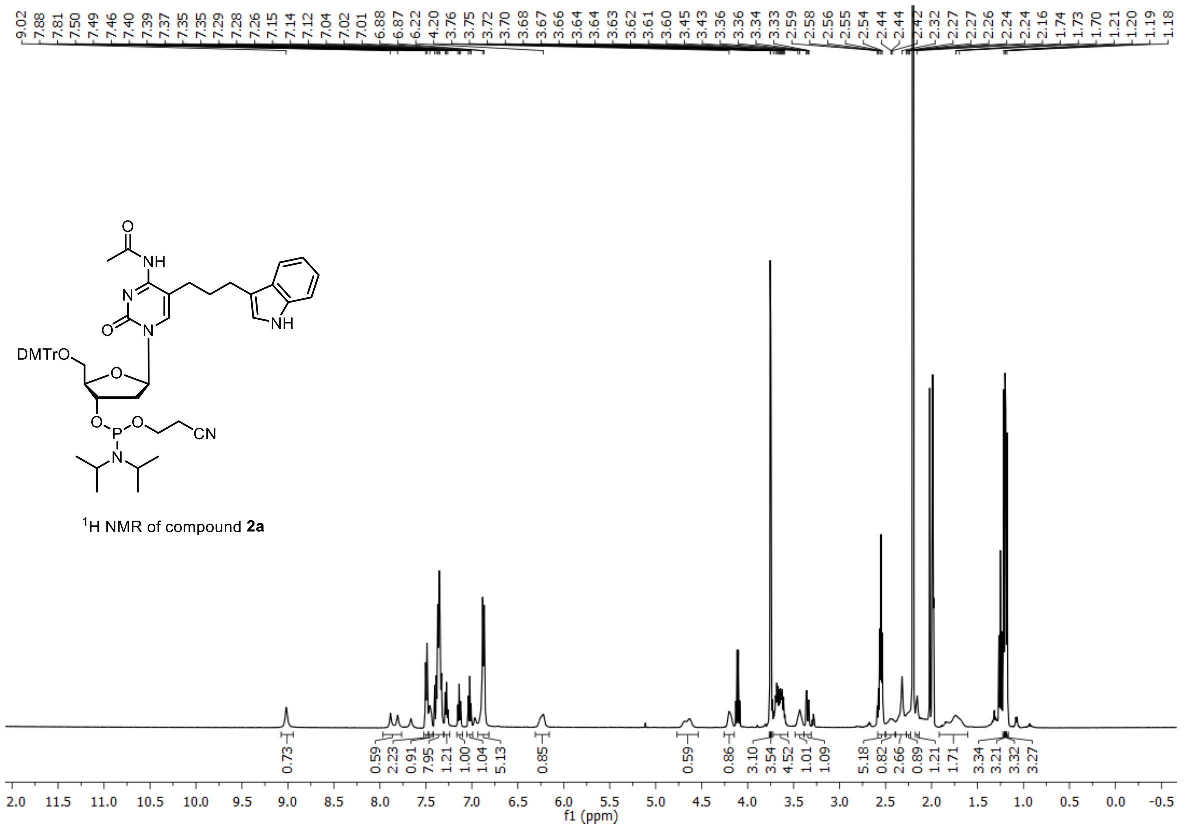

$^{1}$H NMR of compound **2a**

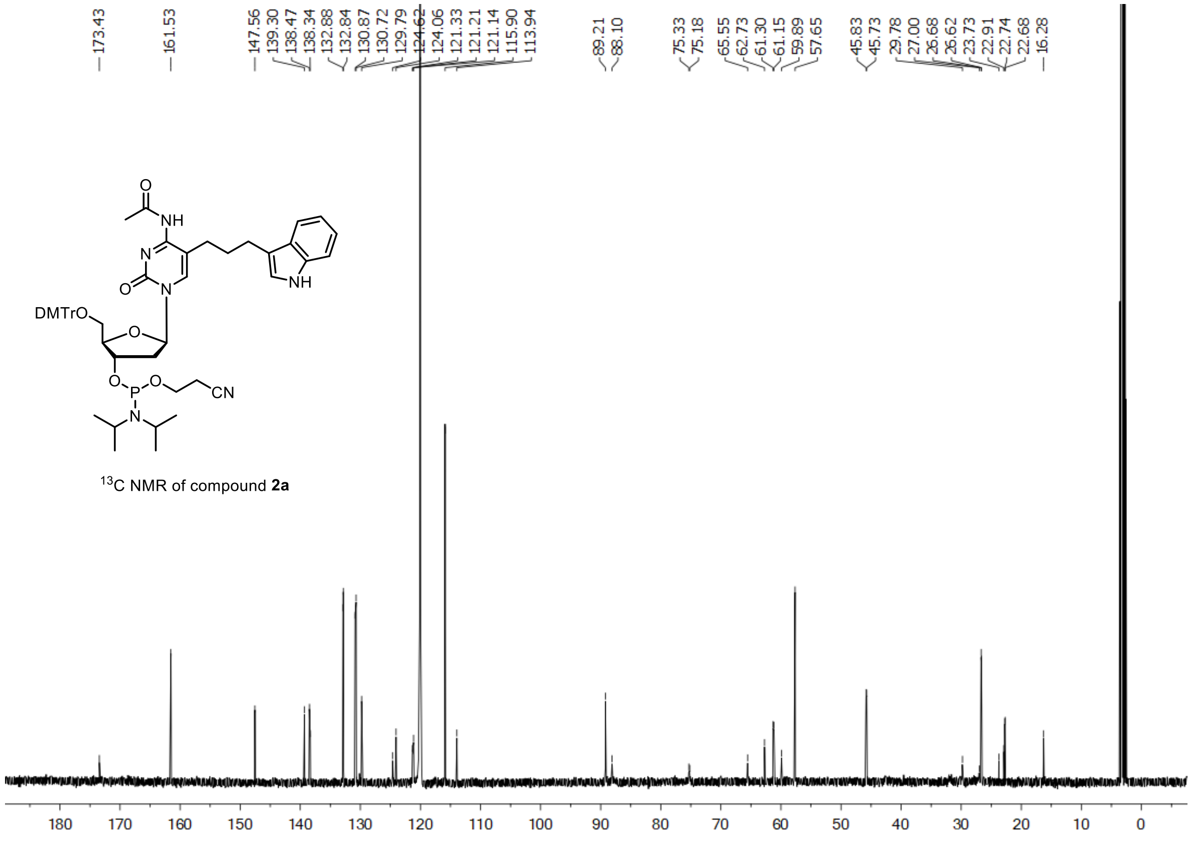

$^{13}$C NMR of compound **2a**

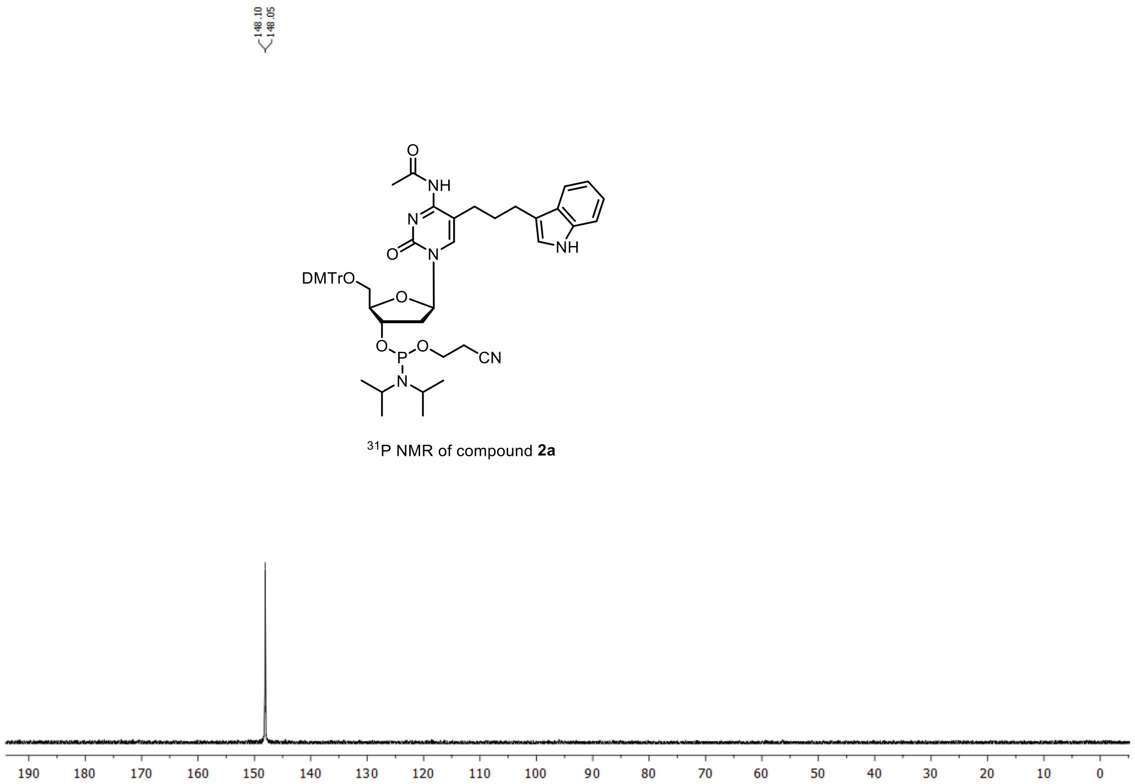


$^{31}$P NMR of compound **2a**

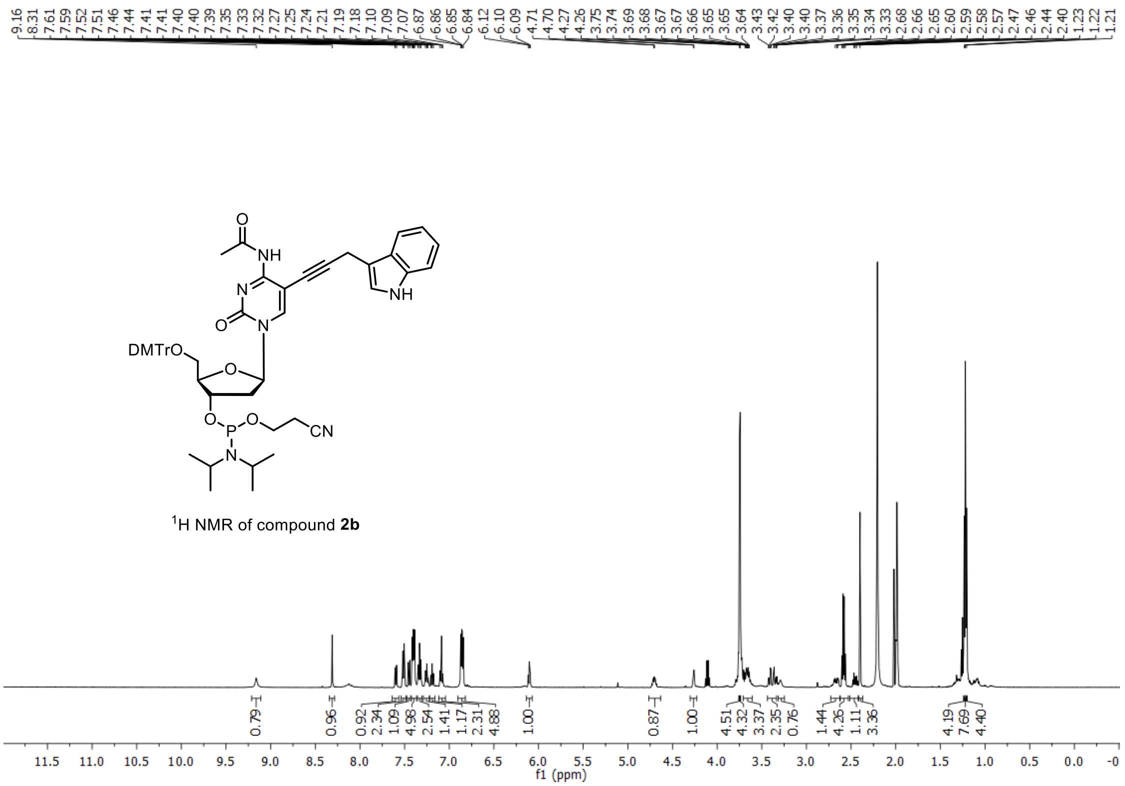


$^{1}$H NMR of compound **2b**

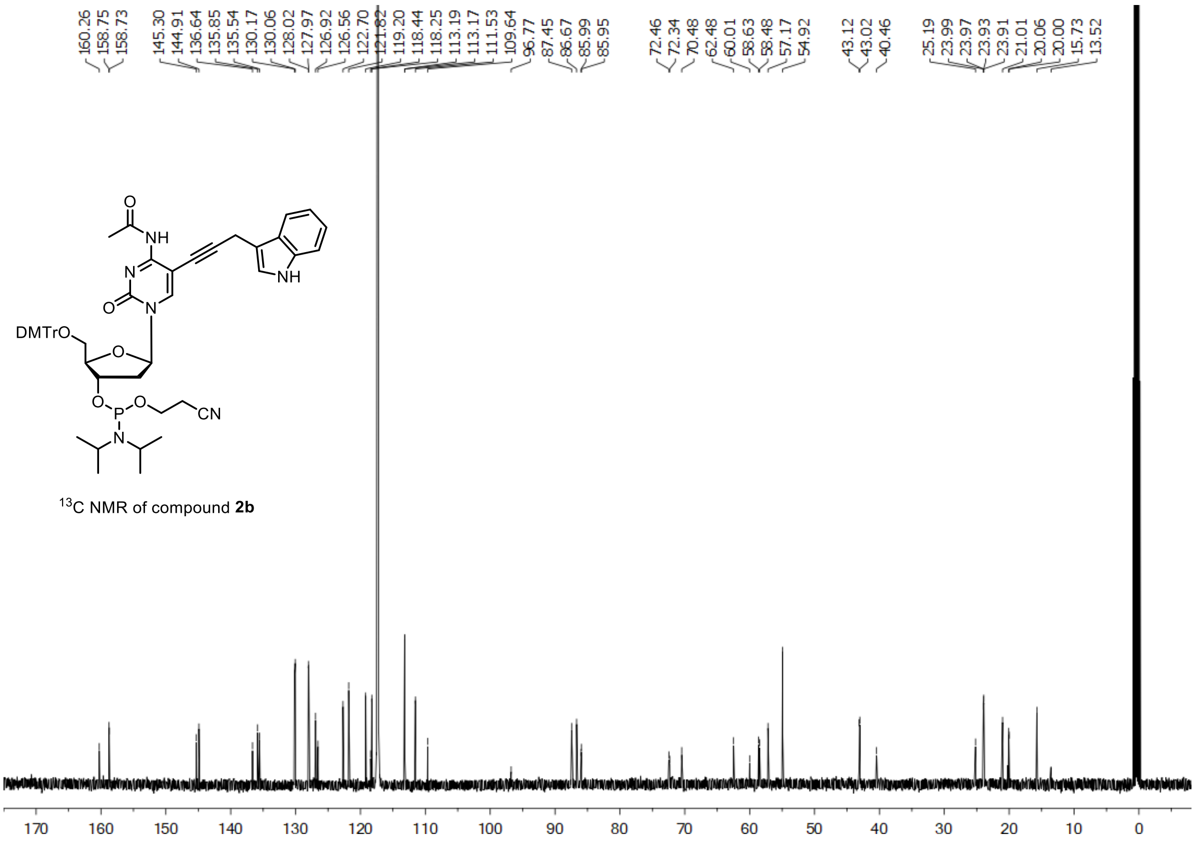


$^{13}C$ NMR of compound **2b**

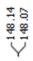


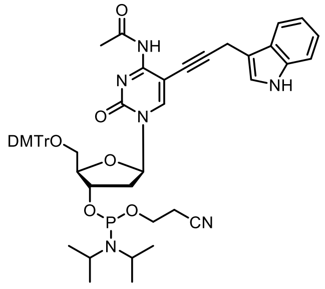


$^{31}P$ NMR of compound **2b**

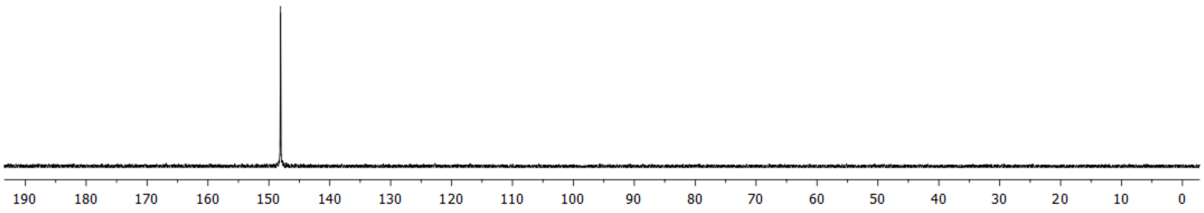

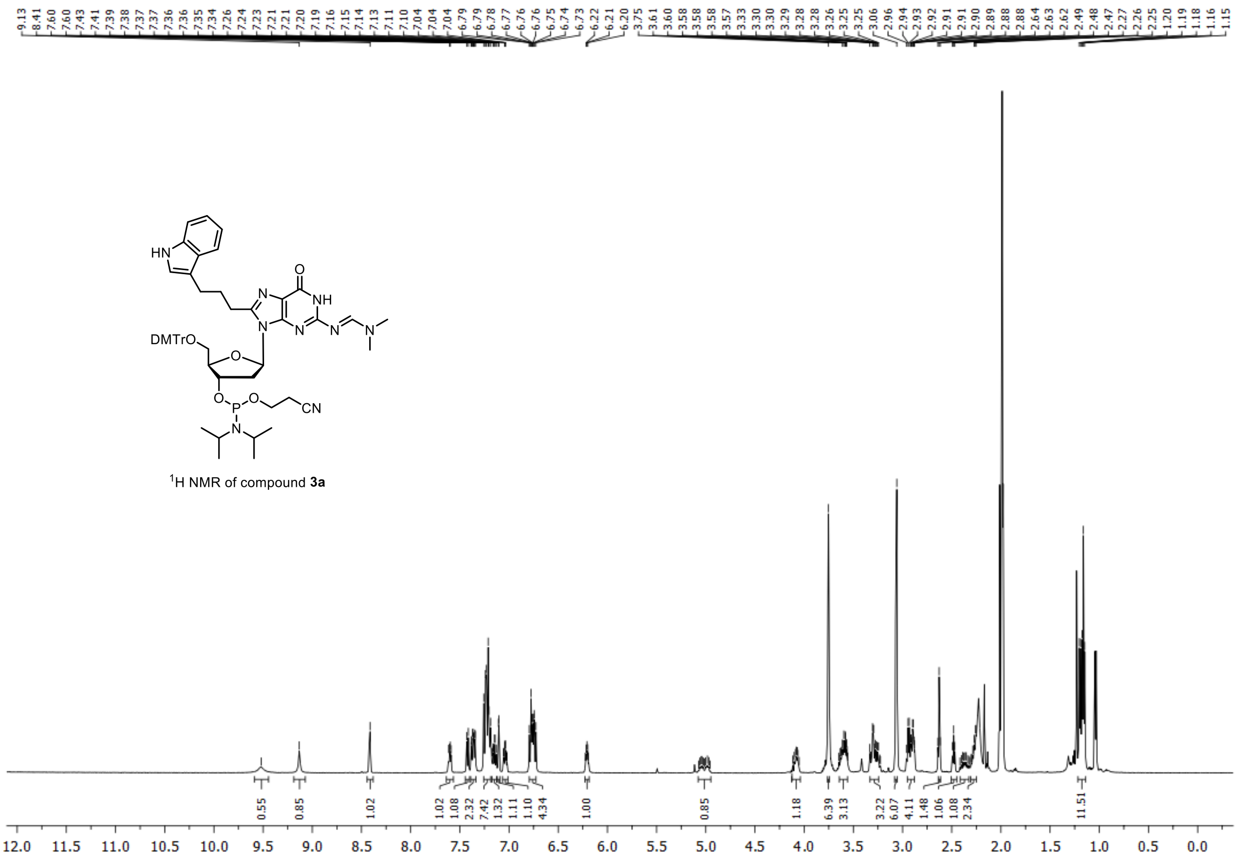

[1]H NMR of compound **3a**

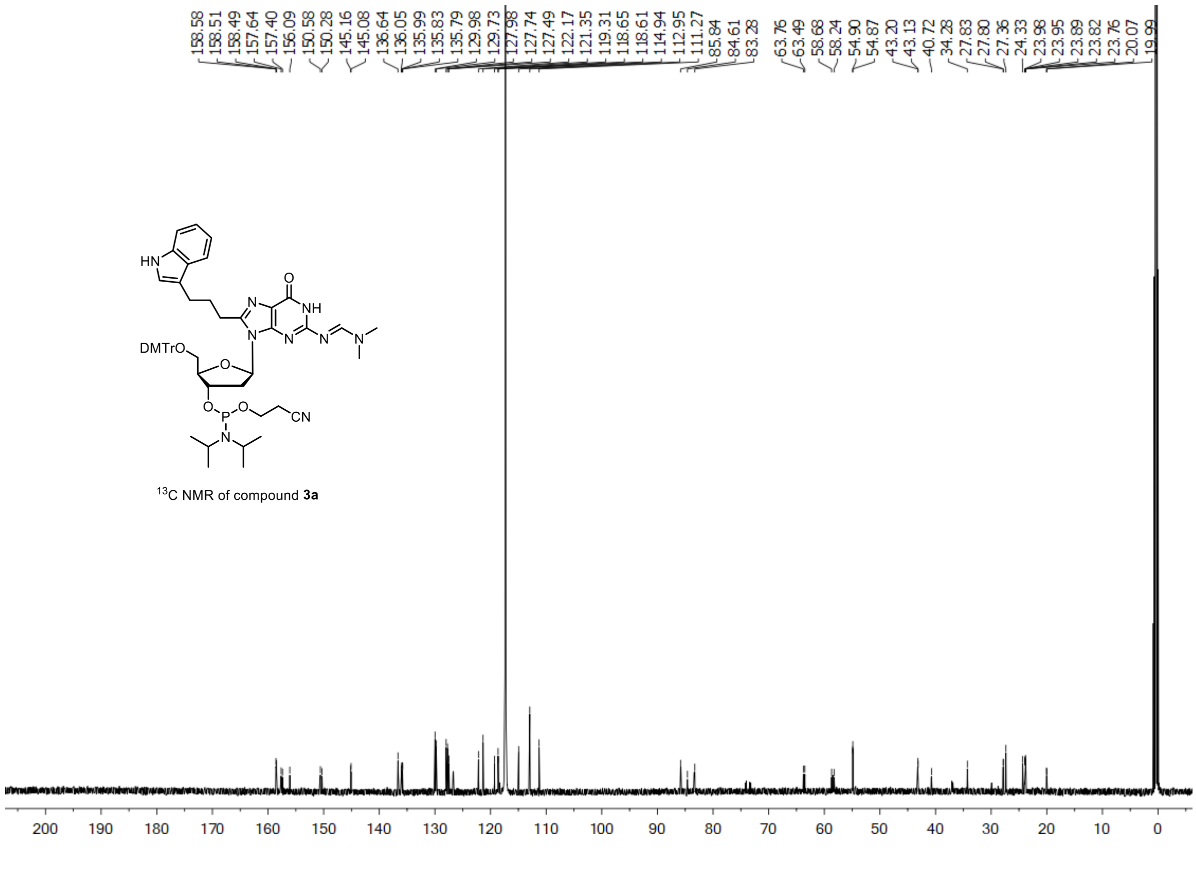

[13]C NMR of compound **3a**

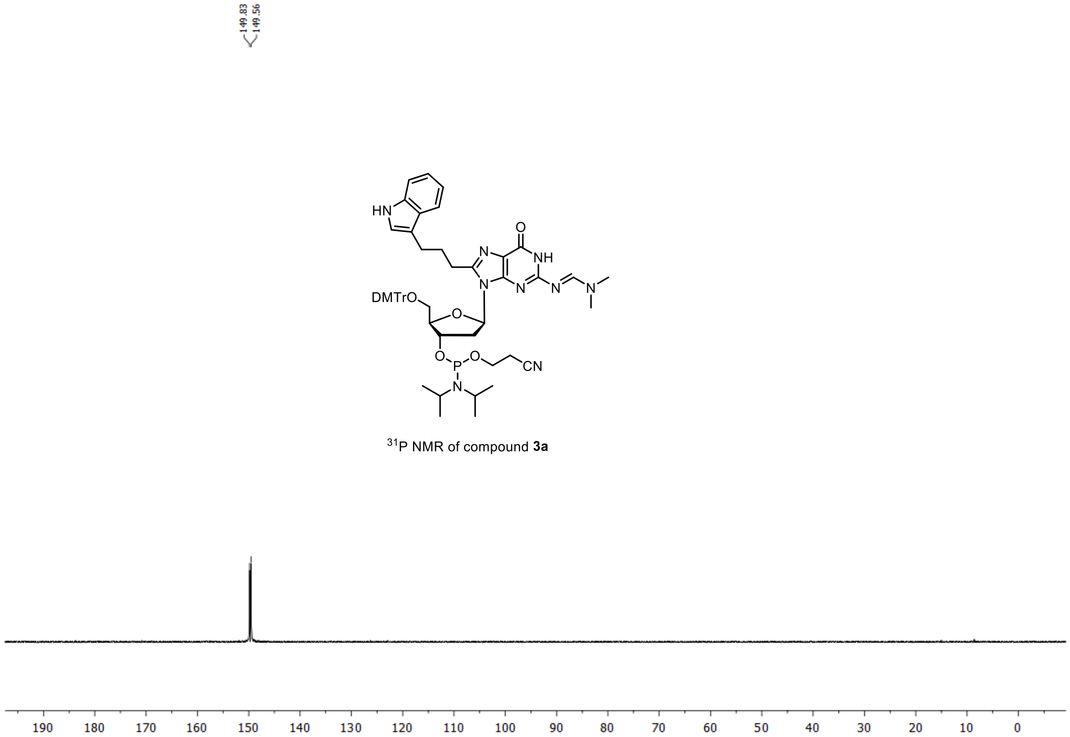


[31]P NMR of compound **3a**

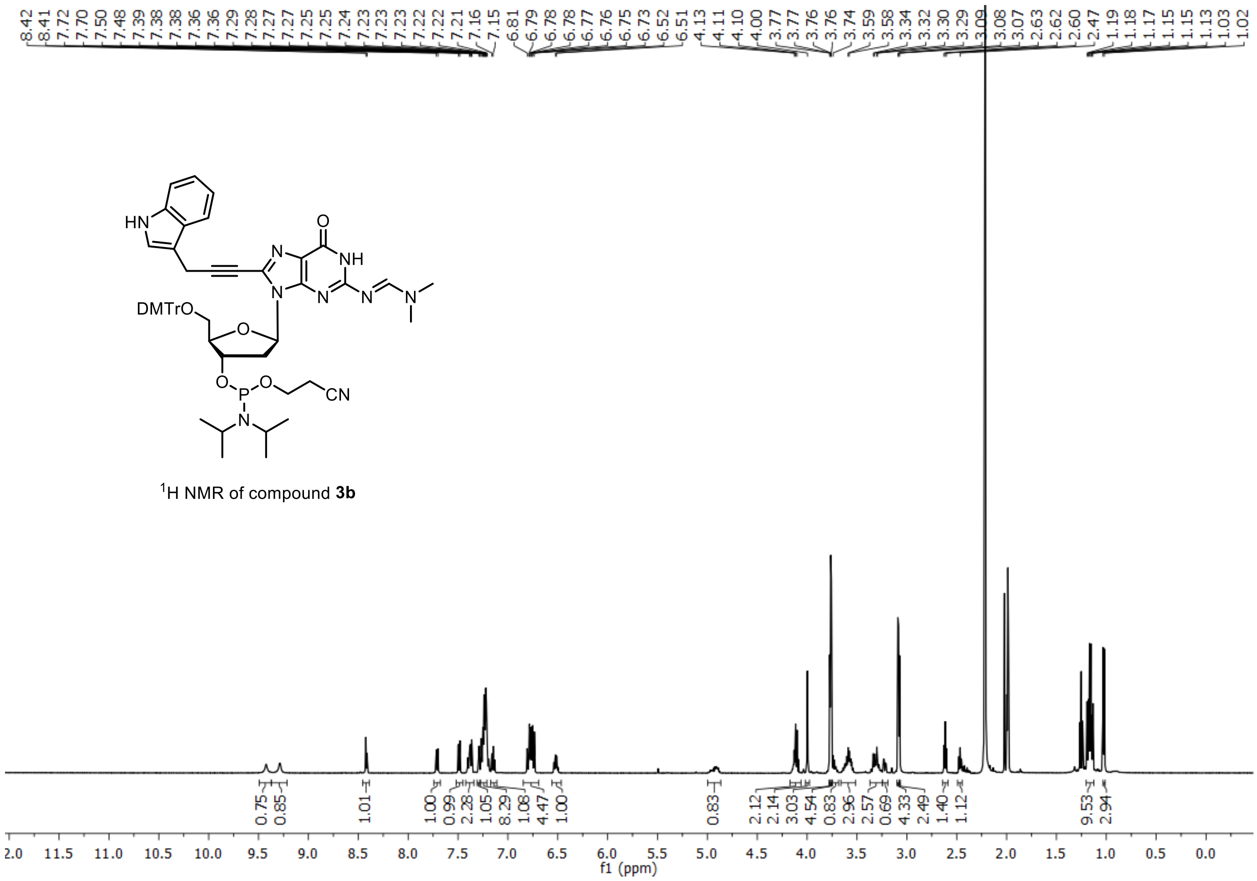


[1]H NMR of compound **3b**

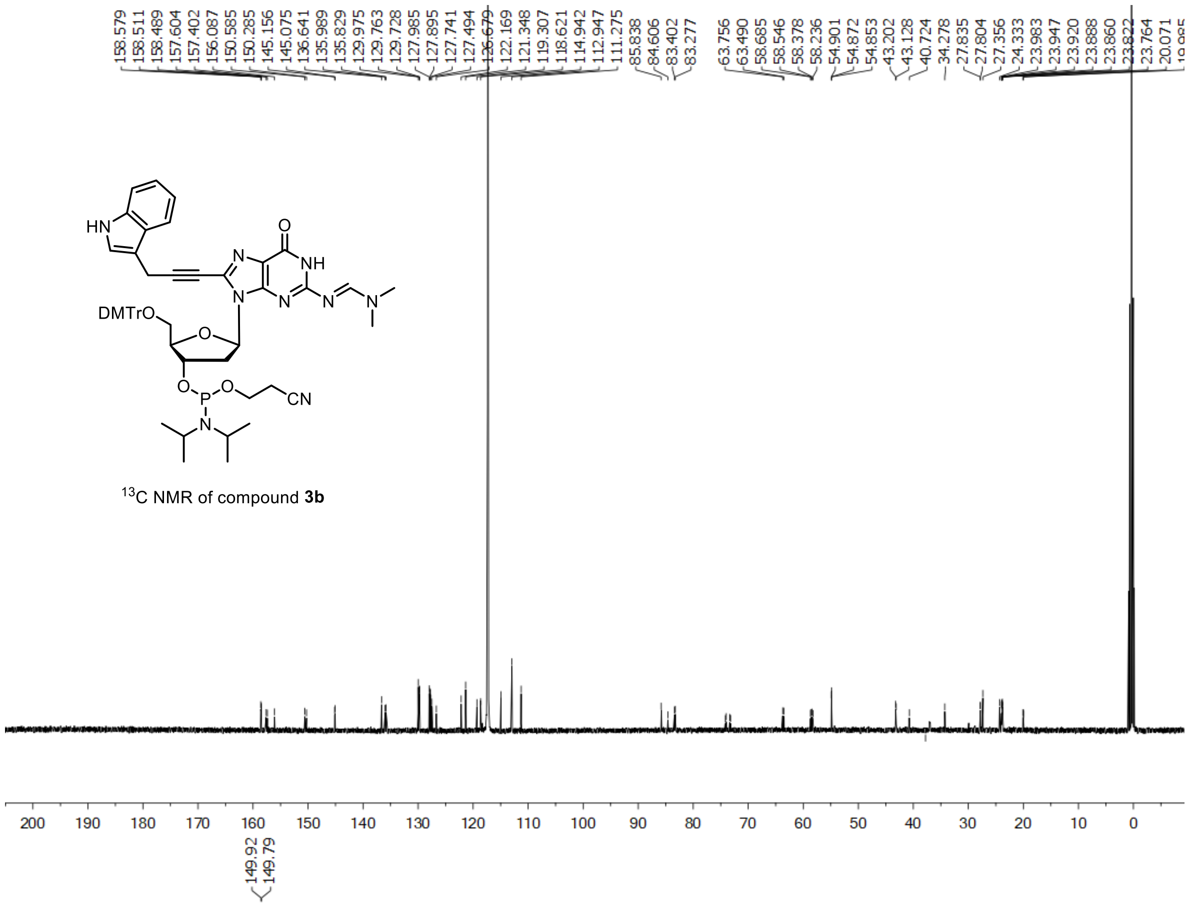


$^{13}$C NMR of compound **3b**

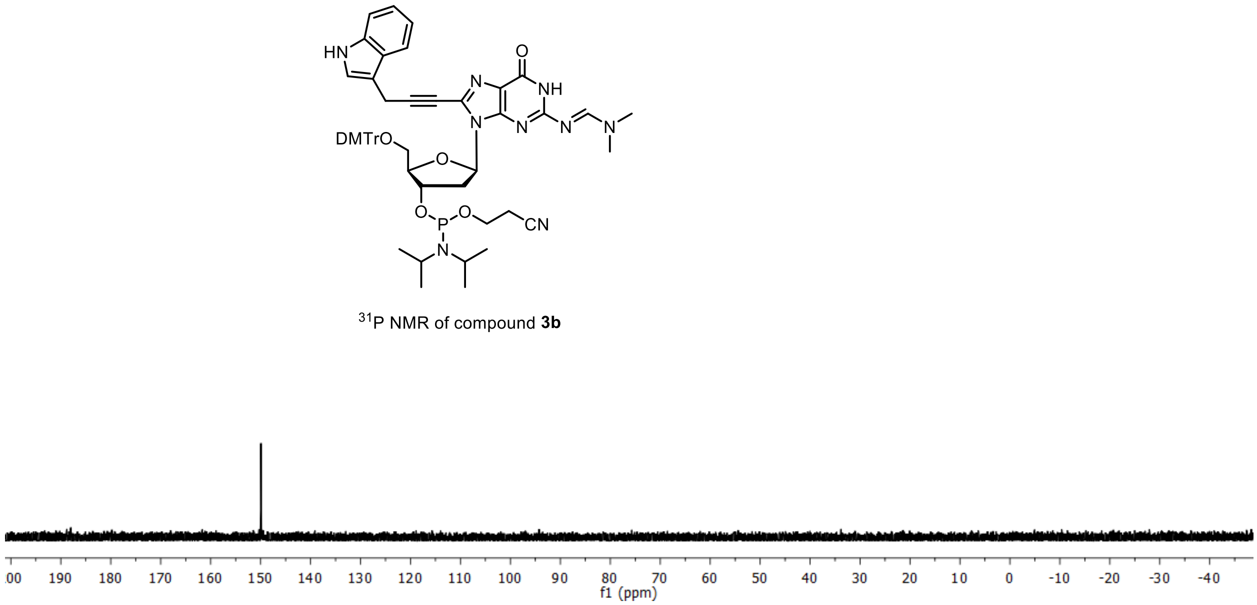


$^{31}$P NMR of compound **3b**

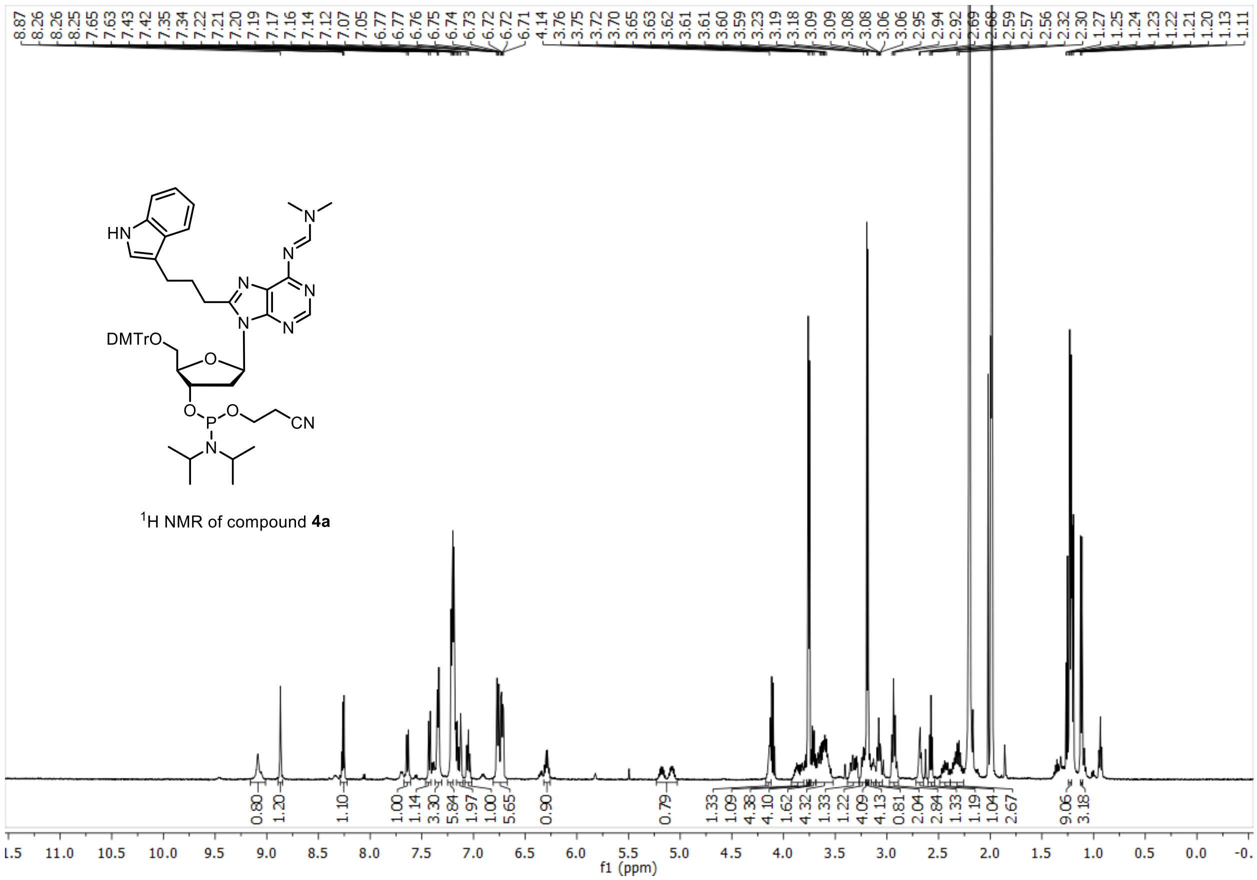


$^{1}$H NMR of compound **4a**

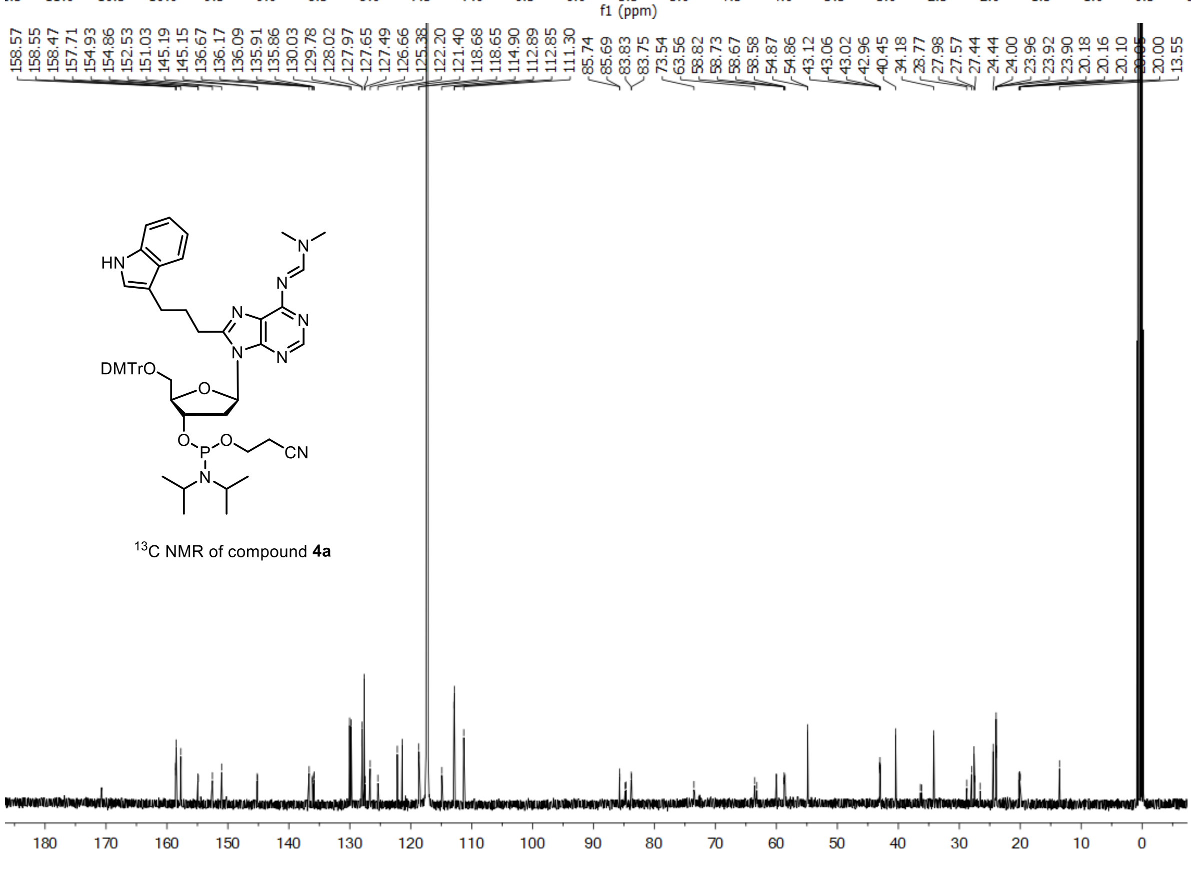


$^{13}$C NMR of compound **4a**

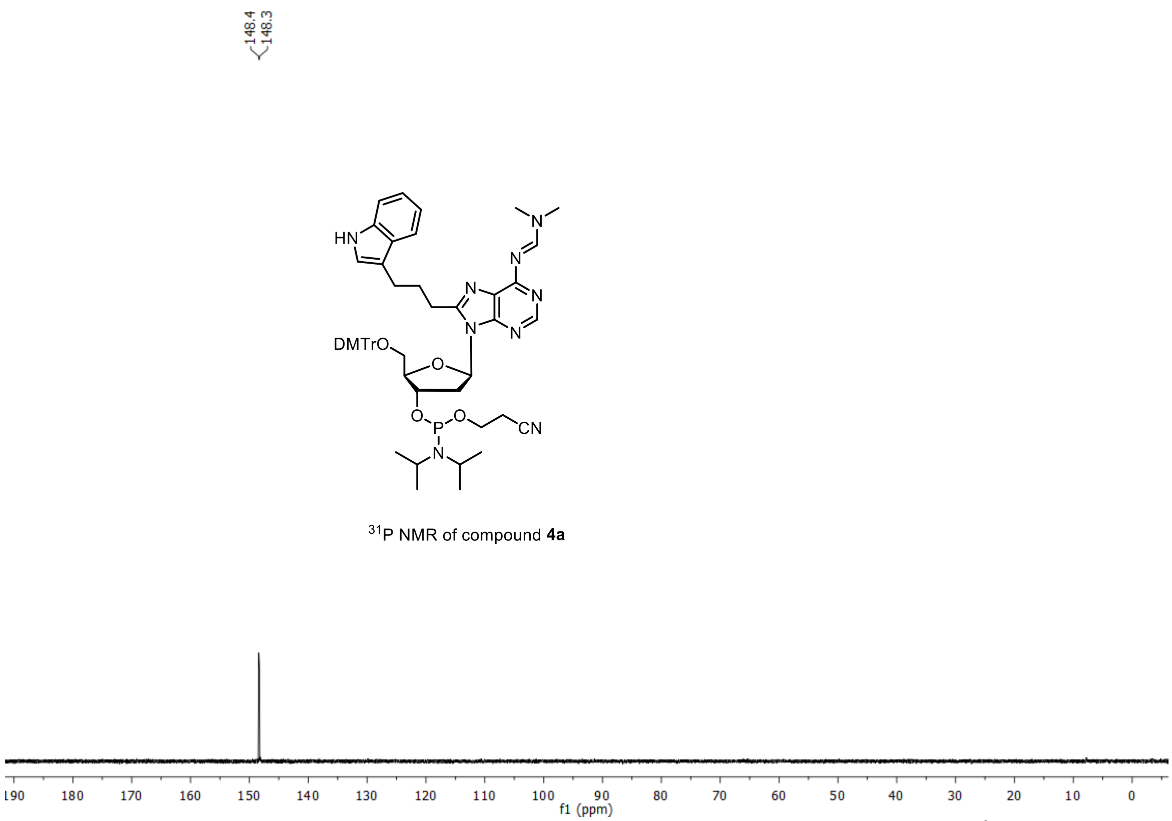


$^{31}$P NMR of compound **4a**

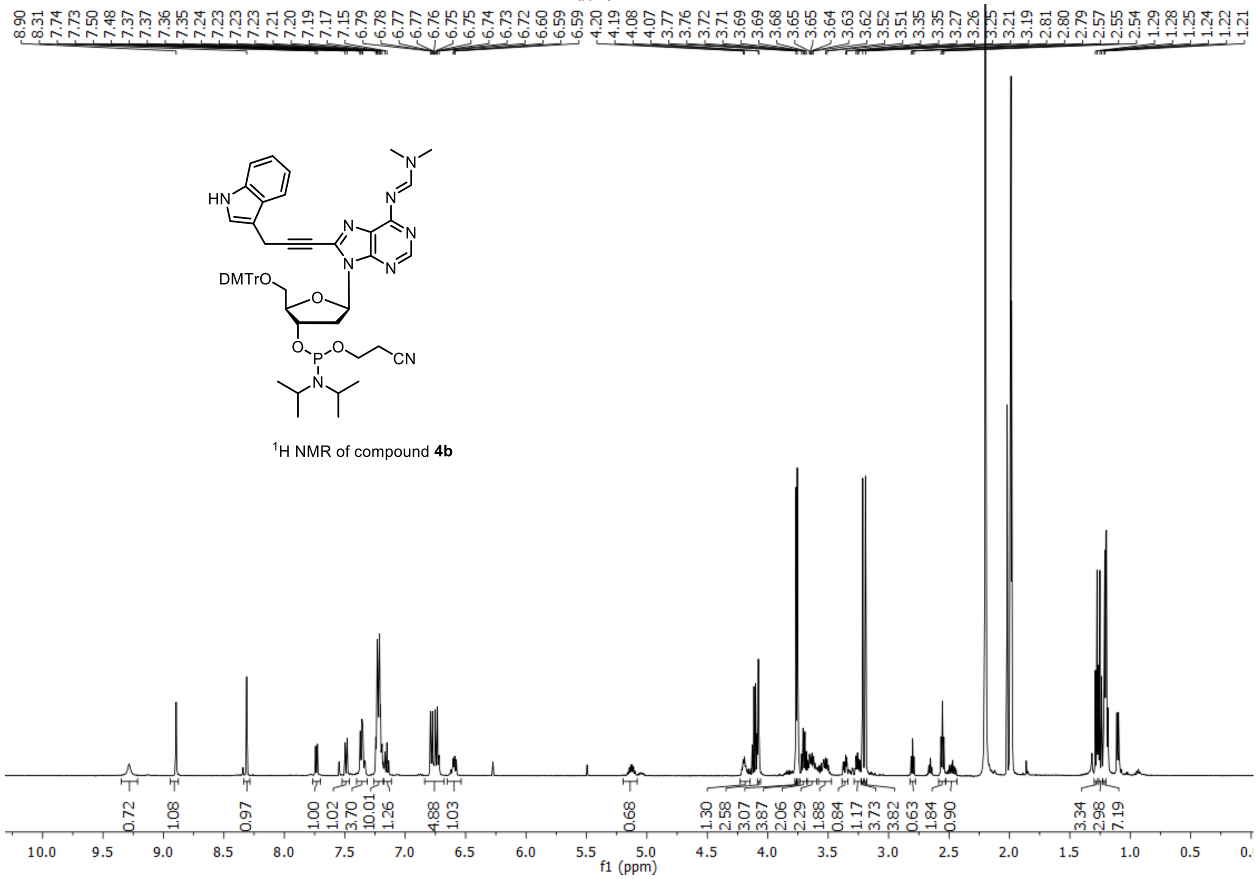


$^{1}$H NMR of compound **4b**

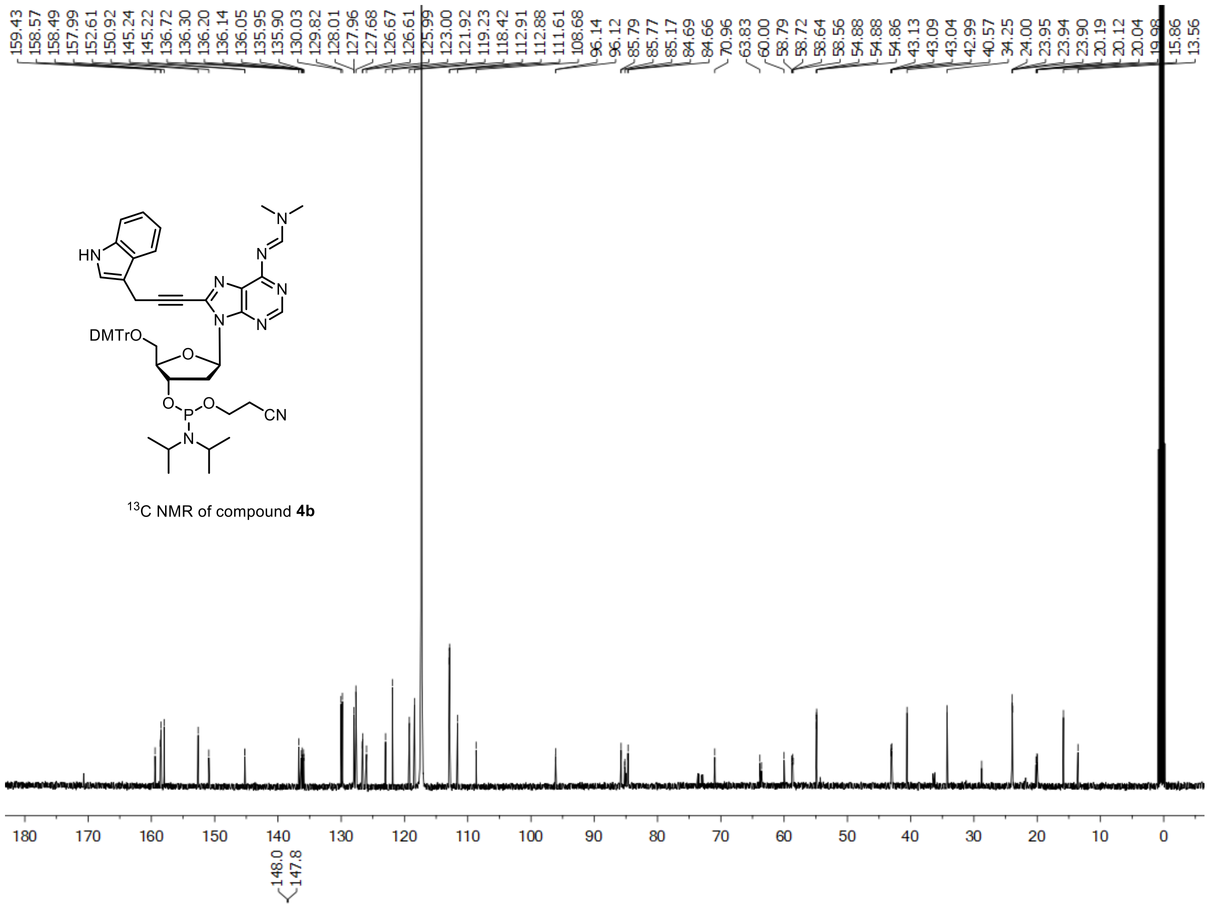


$^{13}C$ NMR of compound **4b**

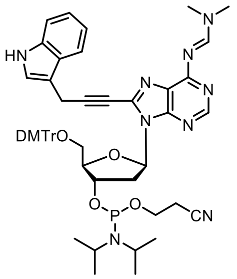


$^{31}P$ NMR of compound **4b**

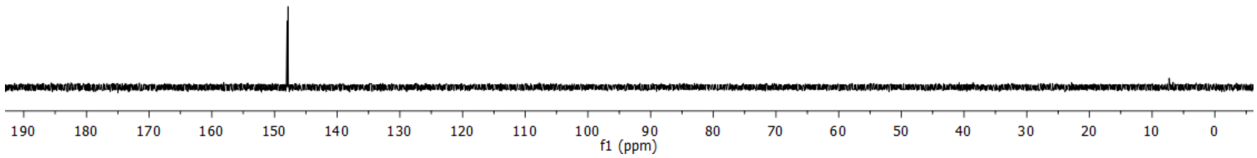

### 6. Mass Spectra of DNA sequences

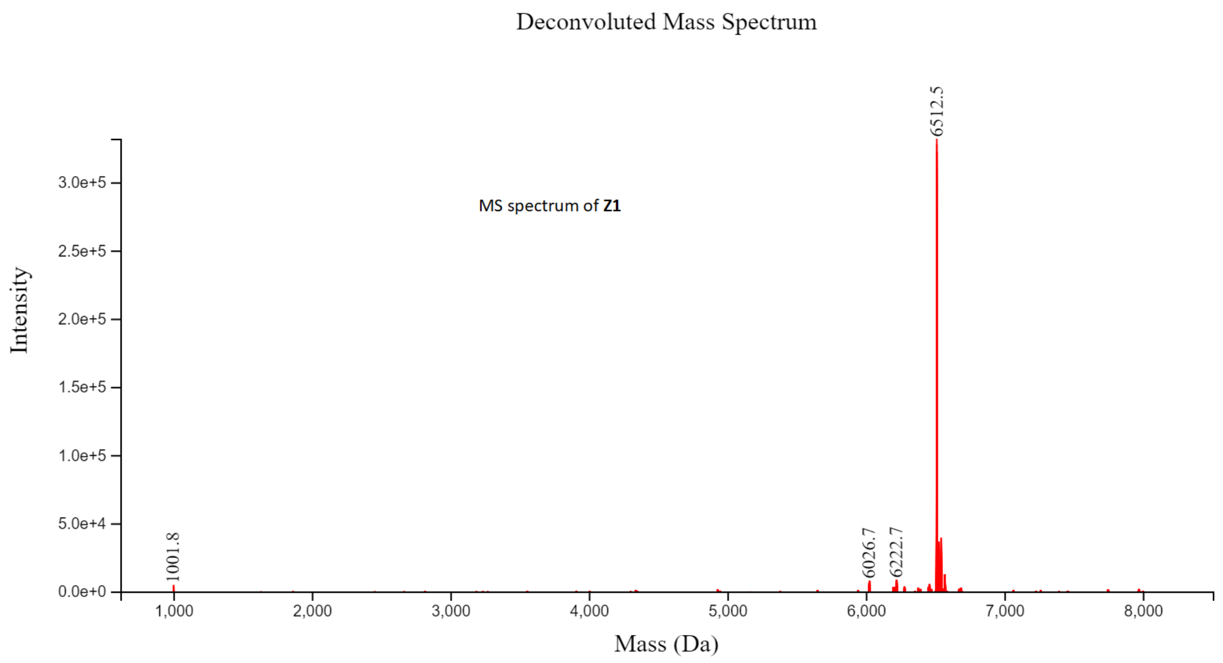


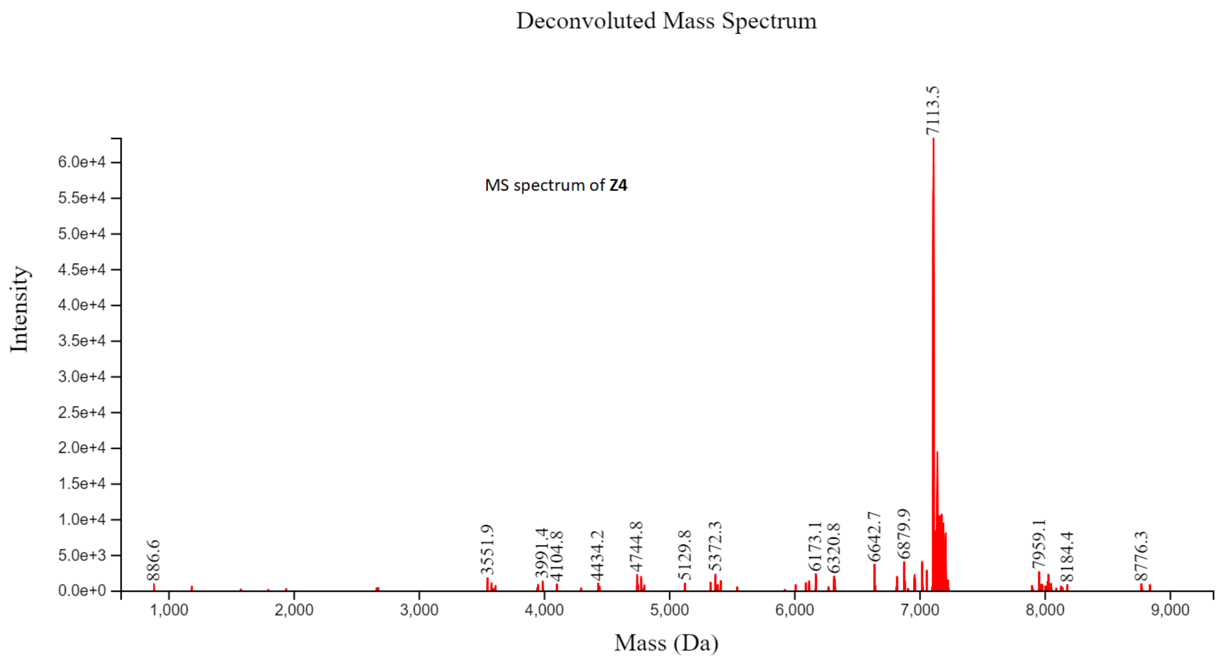

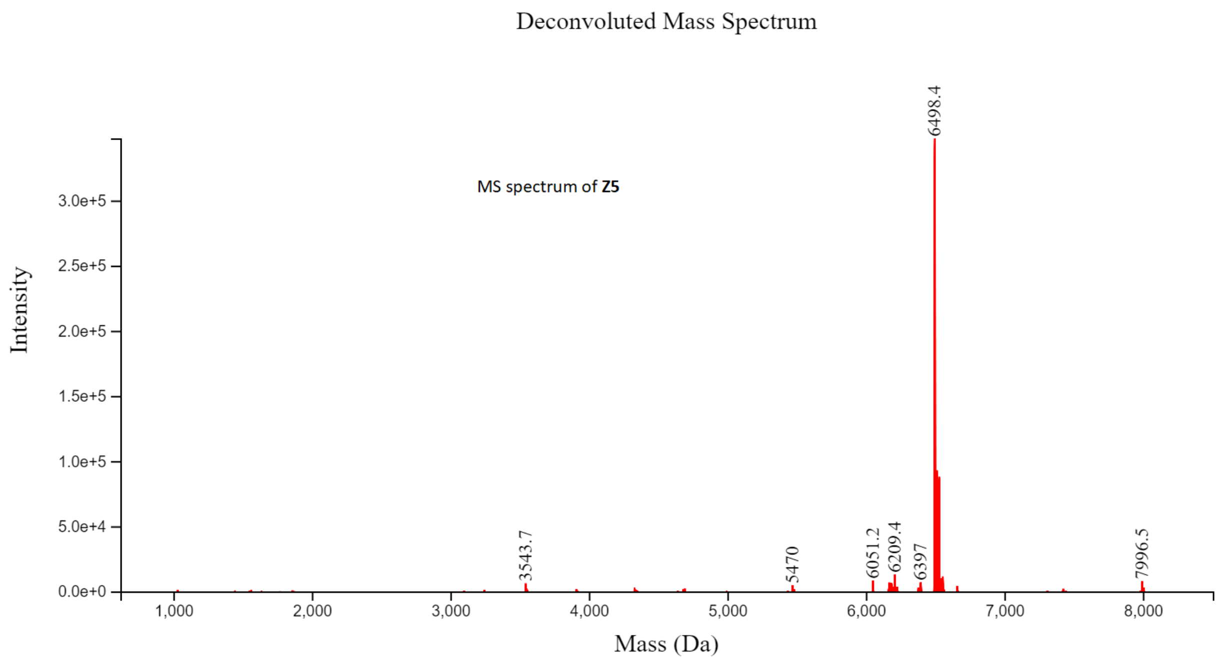
Deconvoluted Mass Spectrum
MS spectrum of **Z5**
Intensity
3.0e+5
2.5e+5
2.0e+5
1.5e+5
1.0e+5
5.0e+4
0.0e+0
6498.4
3543.7
5470
6051.2
6209.4
6397
7996.5
1,000
2,000
3,000
4,000
5,000
6,000
7,000
8,000
Mass (Da)

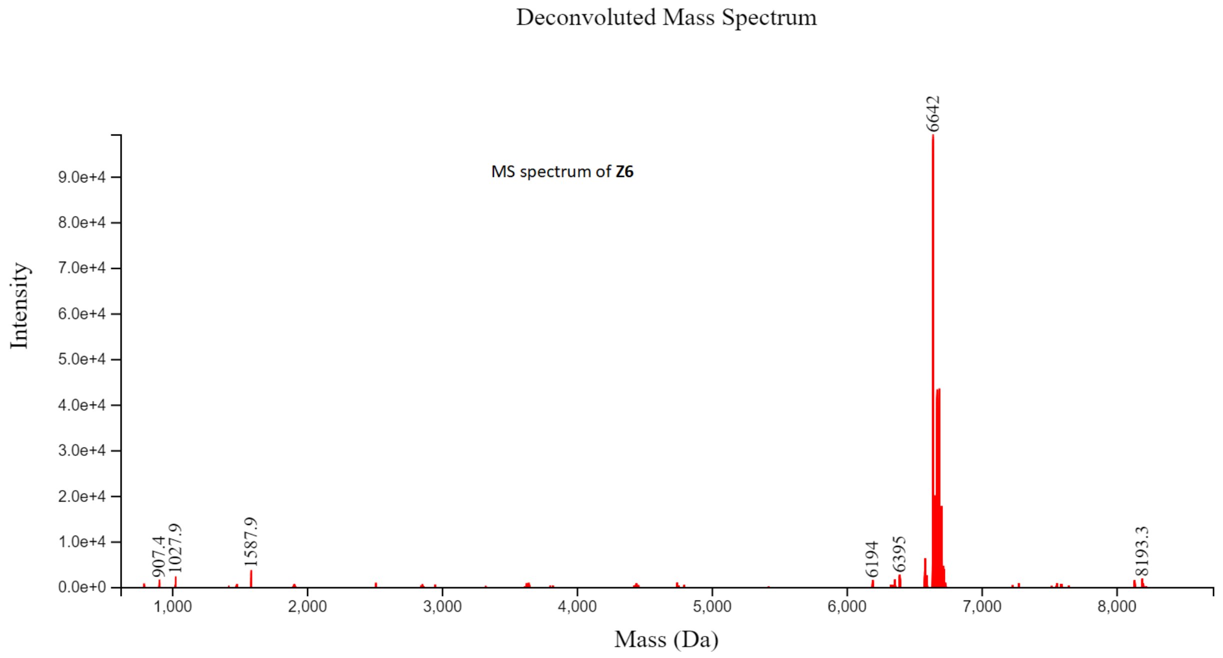
Deconvoluted Mass Spectrum
MS spectrum of **Z6**
Intensity
9.0e+4
8.0e+4
7.0e+4
6.0e+4
5.0e+4
4.0e+4
3.0e+4
2.0e+4
1.0e+4
0.0e+0
6642
907.4
1027.9
1587.9
6194
6395
8193.3
1,000
2,000
3,000
4,000
5,000
6,000
7,000
8,000
Mass (Da)

Deconvoluted Mass Spectrum

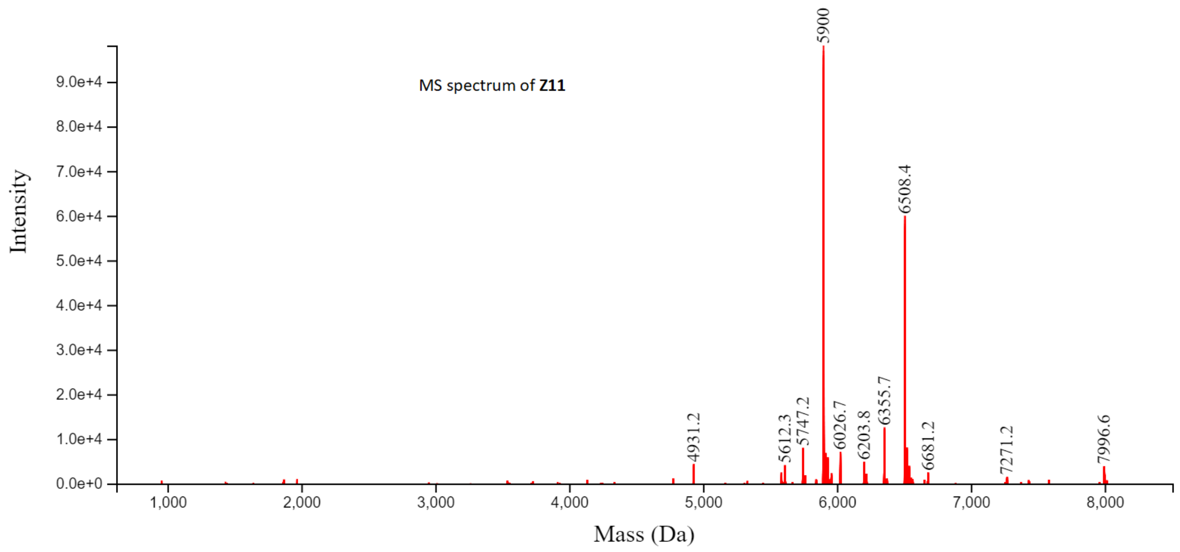


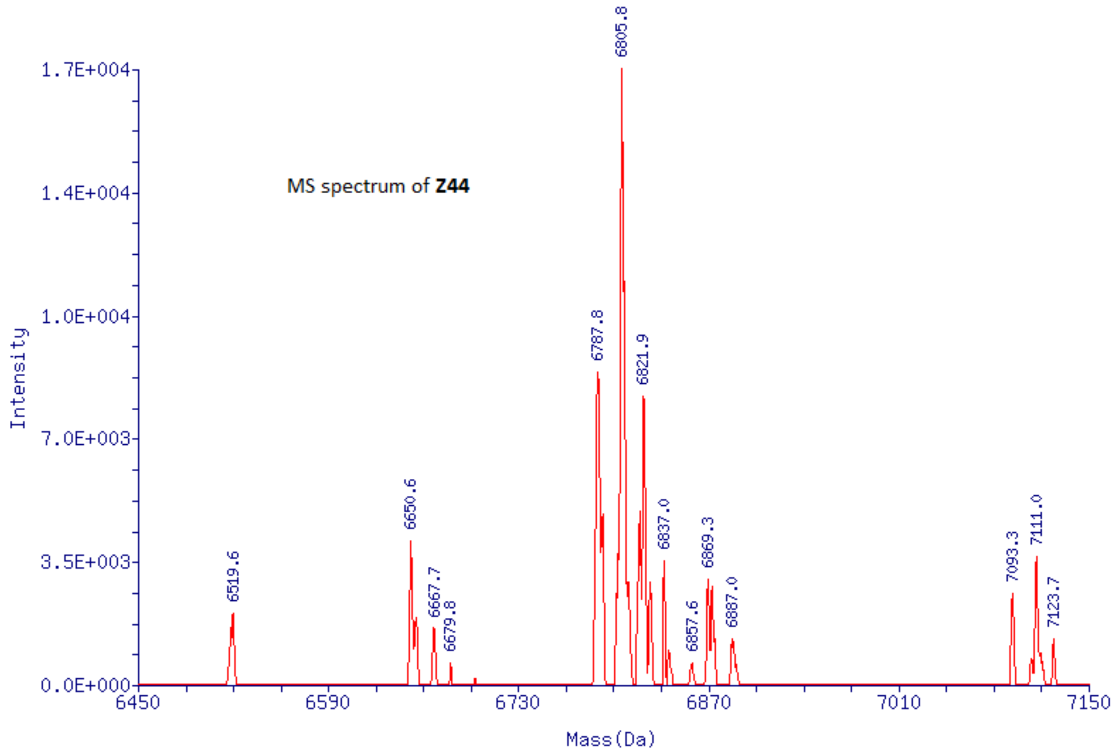

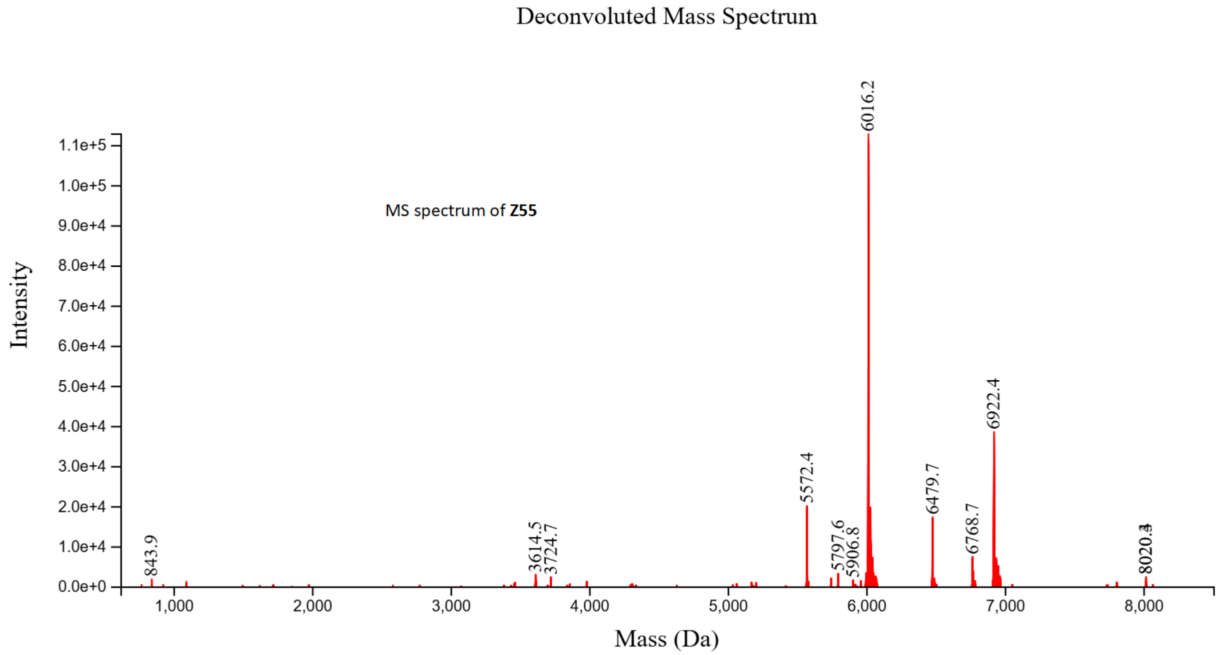
Deconvoluted Mass Spectrum
MS spectrum of **Z55**
Intensity
Mass (Da)
6016.2
6922.4
5572.4
6479.7
6768.7
5797.6
5906.8
3614.5
3724.7
843.9

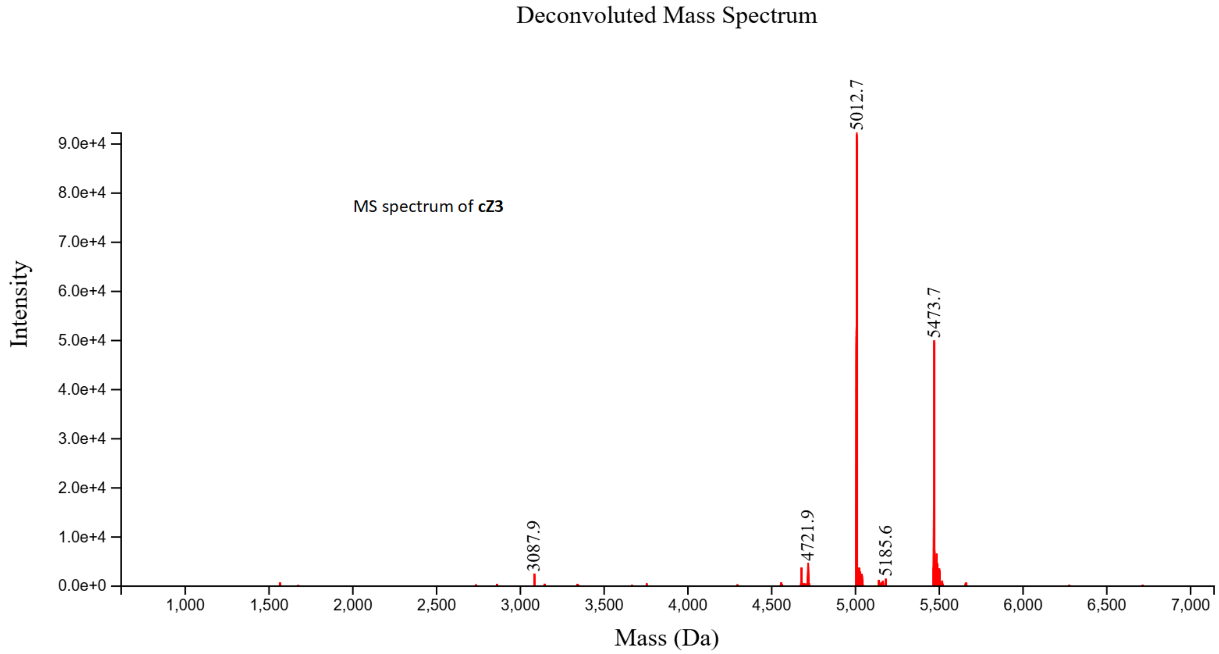
Deconvoluted Mass Spectrum
MS spectrum of **cZ3**
Intensity
Mass (Da)
5012.7
5473.7
4721.9
5185.6
3087.9

Deconvoluted Mass Spectrum

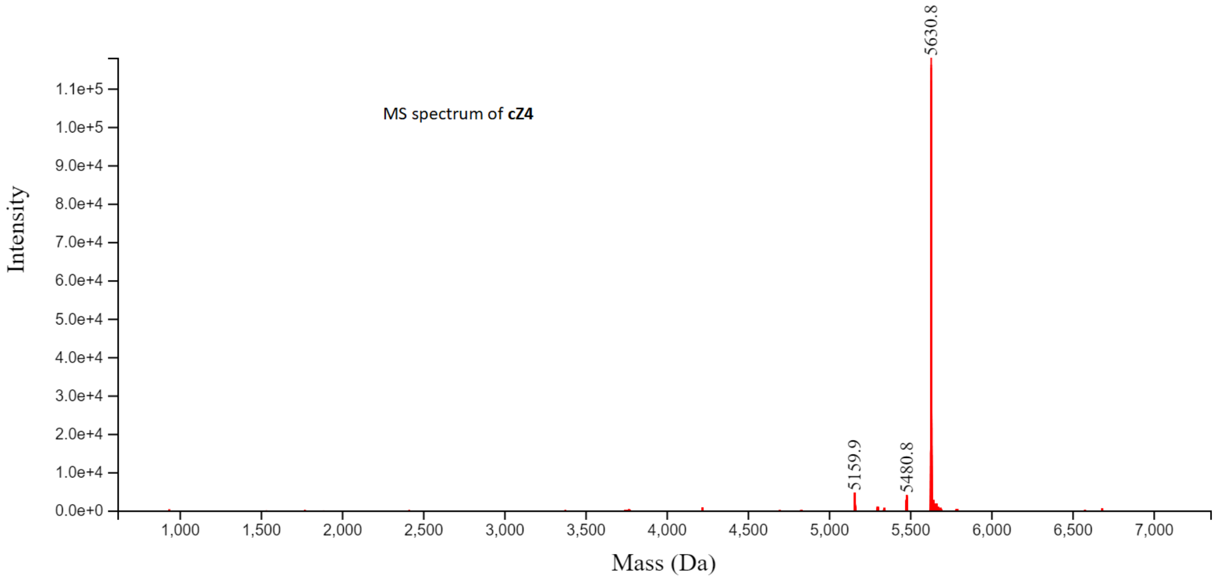


Deconvoluted Mass Spectrum

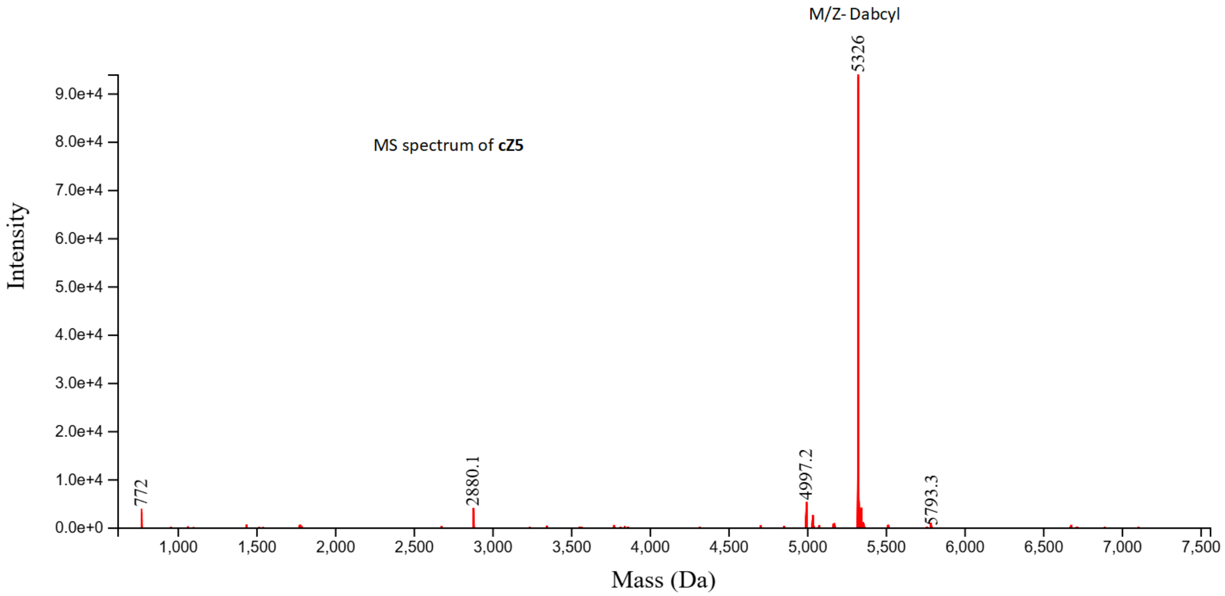


**7.** Reference:


1. Yu. T. Robert.; Friedman, R. K.; Rovis, T. *J. Am. Chem. Soc*. **2009**, *131*, 13250-13251.
